# nELAVL phosphorylation by CDKL5 regulates RNA metabolism and condensates communication to promote experience-dependent maturation of the visual cortex

**DOI:** 10.64898/2026.04.03.716270

**Authors:** Shiyang Yuan, Yao Zhu, Zhongyu Zheng, Hei Matthew Yip, Maggie See Wing Chan, Zhongjie Zhang, Yue Chai, Kyle Robert Jenks, Katya Tsimring, Gregg Robert Heller, Jose Carlos Zepeda, Marco Celotto, Hung Kwok Hung, Yangyang Duan, Shun-Fat Lau, Crystal Wing Yan Ho, Hayley Wing Sum Tsang, Amy Kit Yu Fu, Hei Ming Lai, Andrew John Kwok, Ho Ko, Chunbin Gu, Billy Wai-Lung Ng, Jinyue Liao, Sin Hang Fung, Qin Cao, Stephen Kwok-Wing Tsui, Taeyun Ku, Liming Tan, Andrea Colarusso, Keren Lasker, Yihui Wan, Jinwei Zhu, Jacque Pak Kan Ip

**Affiliations:** School of Biomedical Sciences, The Chinese University of Hong Kong, Hong Kong, China; Department of Brain and Cognitive Sciences, Picower Institute for Learning and Memory, Massachusetts Institute of Technology, Cambridge, MA, USA; Department of Pharmacology, Vanderbilt University, Nashville, TN, USA; Institute for Neural Information Processing, Center for Molecular Neurobiology (ZMNH), University Medical Center Hamburg-Eppendorf (UKE), Hamburg, Germany; Division of Life Science, State Key Laboratory of Molecular Neuroscience and Molecular Neuroscience Center, The Hong Kong University of Science and Technology, Hong Kong, China; Department of Chemical Pathology, The Chinese University of Hong Kong, Hong Kong, China; Department of Medicine and Therapeutics, Faculty of Medicine, The Chinese University of Hong Kong, Hong Kong, China; Li Ka Shing Institute of Health Sciences, Faculty of Medicine, The Chinese University of Hong Kong, Hong Kong, China; Gerald Choa Neuroscience Institute, The Chinese University of Hong Kong, Hong Kong, China; Guangdong-Hong Kong-Macao Joint Laboratory for New Drug Screening, School of Pharmacy, Faculty of Medicine, The Chinese University of Hong Kong, Hong Kong, China; Peter Hung Pain Research Institute, Faculty of Medicine, The Chinese University of Hong Kong, Hong Kong, China; Department of Ophthalmology, LKS Faculty of Medicine, The University of Hong Kong, Hong Kong, China; Graduate School of Medical Science and Engineering, Korea Advanced Institute of Science and Technology, Daejeon, Republic of Korea; Shenzhen Key Laboratory of Neuropsychiatric Modulation, Shenzhen-Hong Kong Institute of Brain Science, Shenzhen Institutes of Advanced Technology, Chinese Academy of Sciences, Shenzhen, China; Guangdong Provincial Key Laboratory of Brain Connectome and Behavior, the Brain Cognition and Brain Disease Institute, Shenzhen Institutes of Advanced Technology, Chinese Academy of Sciences, Shenzhen, China; Department of Integrative Structural and Computational Biology, Scripps Research, La Jolla, California, USA; Bio-X Institutes, Key Laboratory for the Genetics of Developmental and Neuropsychiatric Disorders, Shanghai Jiao Tong University, Shanghai, China; Department of Nephrology, Shanghai Sixth People’s Hospital Affiliated to Shanghai Jiao Tong University School of Medicine, Shanghai, China; Kunming Institute of Zoology - The Chinese University of Hong Kong (KIZ-CUHK) Joint Laboratory of Bioresources and Molecular Research of Common Diseases, The Chinese University of Hong Kong; CUHK Gerald Choa Neuroscience Institute – Oujiang Laboratory Joint Laboratory for Neuroscience and Neurology, Wenzhou, Zhejiang Province, China

## Abstract

Mutations in Cyclin-Dependent Kinase-Like 5 (CDKL5) cause CDKL5 deficiency disorder (CDD), an X-linked neurodevelopmental condition. Through a phosphoproteomic screen, we identified the neuron-specific nELAVL family of RNA-binding proteins as direct activity-dependent substrates of CDKL5. In support of this regulatory axis, single-nuclei transcriptomics of *Cdkl5* knockout (KO) cortices revealed an enriched reduction in activity-dependent mRNAs. Mechanistically, we show that nELAVL proteins undergo phase separation to form biomolecular condensates, the size of which is gated by CDKL5 phosphorylation. Loss of CDKL5 leads to enlarged nELAVL condensates, which exhibit reduced binding affinity for the target mRNA *Fos*, resulting in its accelerated degradation. This disruption extends to inter-condensate communication: phosphodeficient nELAVL show diminished interaction with P-bodies, which themselves become enlarged in CDD mutant iNeurons. Functionally, the absence of nELAVL phosphorylation recapitulates the deficits in experience-dependent visual function observed in *Cdkl5* KO mice. Our findings establish a critical molecular mechanism by which CDKL5-mediated phosphorylation governs mRNA metabolism by tuning the properties of nELAVL condensates and their communication with other biomolecular condensates, ultimately promoting experience-dependent maturation of the visual cortex.

## Introduction

CDD is a severe, X-linked neurodevelopmental disorder caused by loss of function mutations in the *CDKL5* gene^1^. Initially categorized as an early-onset-seizure variant of Rett syndrome, CDD is now recognized as a distinct condition with its own ICD-10 code (G40.42). CDD is characterized by a range of neurodevelopmental features including early-onset epilepsy, motor dysfunction, learning disabilities, autistic features and severe neurodevelopmental delay^2,3^. CDKL5 is a serine/threonine kinase highly expressed in the brain^4^. *Cdkl5* knockout (KO) mice have been shown to exhibit autistic-like phenotypes, impairments in motor control and fear memory^5^, as well as reduced amplitude of visually evoked potentials and diminished visual acuity^6^. *Cdkl5* KO also exhibit impaired dendritic spine development and microtubule dynamics^7,8^. Phosphorylation targets of CDKL5 include microtubule-binding proteins EB2 and MAP1S, transcriptional regulators ELOA and SOX9, and the voltage-gated Ca^2+^ channel Cav2.3^7,9–12^. However, clinically relevant phenotypes of CDD rodent model such as motor deficit or cortical visual impairment were not fully recapitulated in the phospho-deficient knock-in mouse models of the abovementioned CDKL5 phosphorylation targets^7,13^, suggesting that critical CDKL5-mediated functions or phosphorylation targets remain undiscovered.

Here, we present evidence that CDKL5 regulates the refinement of binocular visual circuits in mouse V1 through a previously unidentified phosphorylation target. We performed an unbiased phosphoproteomic screen and identified a family of RNA-binding proteins (RBPs), the neuronal Embryonic Lethal, Abnormal Vision-Like proteins (nELAVLs), as direct substrates of CDKL5. The nELAVL proteins, identified as being recruited into ribonucleoprotein (RNP) granules^14–16^, play crucial roles in regulating RNA stability, localization, and activity-dependent translation^17,18^.

Considering that post-translational modifications, such as phosphorylation, can impact the phase transition of RNP granules, we postulated that CDKL5-mediated phosphorylation would modulate the propensity of nELAVLs to form condensates. Supporting this hypothesis, we found that nELAVL positive condensates are aberrantly enlarged in *Cdkl5* KO neurons. Furthermore, we observed that CDKL5-mediated phosphorylation also affects the size of other biomolecular condensates, including those labelled with Processing (P)-body marker GW182 and stress granules marker TIA1, suggesting that CDKL5 is a key gatekeeper for fine-tuning condensate properties. Consistent with this notion, we found that mRNA levels of various activity-dependent genes were reduced in the V1 of *Cdkl5* KO mice, linking these molecular changes to the impaired binocular matching of visual responses. Mechanistically, phospho-deficient mutant of ELAVL3 exhibited reduced binding affinity with its target mRNA *Fos* resulting in faster degradation. Functionally, we demonstrate that the orientation selectivity and binocular matching of visual responses is impaired in *Cdkl5* KO mouse V1 neurons with the ipsilateral eye responses more severely affected. Furthermore, we showed that AAV-mediated expression of nELAVL phospho-deficient in mouse bV1 recapitulated the deficits observed in *Cdkl5* KO mice.

Taken together, these findings are consistent with a model of cortical visual impairment in CDD in which enlarged nELAVL condensates hampers activity-dependent mRNA regulation, thereby impairing orientation selectivity and binocular matching of V1 neurons.

## Results

### CDKL5 phosphorylates nELAVLs in an activity-dependent manner

Cortical visual impairment is a phenomenon common amongst neurodevelopmental disorders^6,19–25^. In particular, amongst the 75% of individuals diagnosed with Cyclin-Dependent Kinase-Like 5 (CDKL5) deficiency disorder (CDD)^4,6,21,26–30^, an X-linked neurodevelopmental disorder attributed by loss of function mutations in the *CDKL5* gene^1^, cortical visual impairment correlates with delayed developmental milestones and has therefore been proposed to be a potential biomarker^21^. Given the clinical relevance of the primary visual cortex (V1), we first sought to identify relevant direct substrates of CDKL5 by applying an unbiased phosphoproteomic screen using V1 tissue from WT and *Cdkl5* KO mice. We used Tandem Mass Tag-based phosphoproteomics analysis and identified 9048 phosphopeptides on 3146 phosphoproteins in V1 (Figure 1A; table S1). Among them, 119 phosphopeptides were upregulated and 105 downregulated in *Cdkl5* KO mice, corresponding to 88 and 73 phosphoproteins, respectively (Figure 1B). The number and distribution of phosphorylation sites on all identified proteins were analyzed (Figure S1A & B). Gene Ontology (GO) analysis showed that the downregulated phosphopeptides in V1 were predominantly related to synaptic function or structural constituent of the postsynaptic density (Figure 1C). Seven of the downregulated phosphorylated proteins contained the CDKL5 consensus phosphorylation sequences, Arg-Pro-X-Ser-Ala (RPXS*A) or Arg-Pro-X-Ser (RPXS*)^31^, suggesting they are potentially direct substrates of CDKL5 (Figure S1C). nELAVLs were the most significantly down-regulated phosphorylated proteins among these, and the phosphorylation sites on ELAVL2 (S119), 3 (S119) and 4 (S131) are evolutionary conserved among human, mouse, and rat (Figure 1D). ELAVL2, ELAVL3 and ELAVL4 have 80% sequence homology, are predominantly expressed in the nervous system and are therefore collectively referred as nELAVLs^32^. nELAVLs bind to AU-rich elements in a wide range of target mRNAs to regulate RNA stability, localization, and translation^17,18,33^.

**Figure 1.**
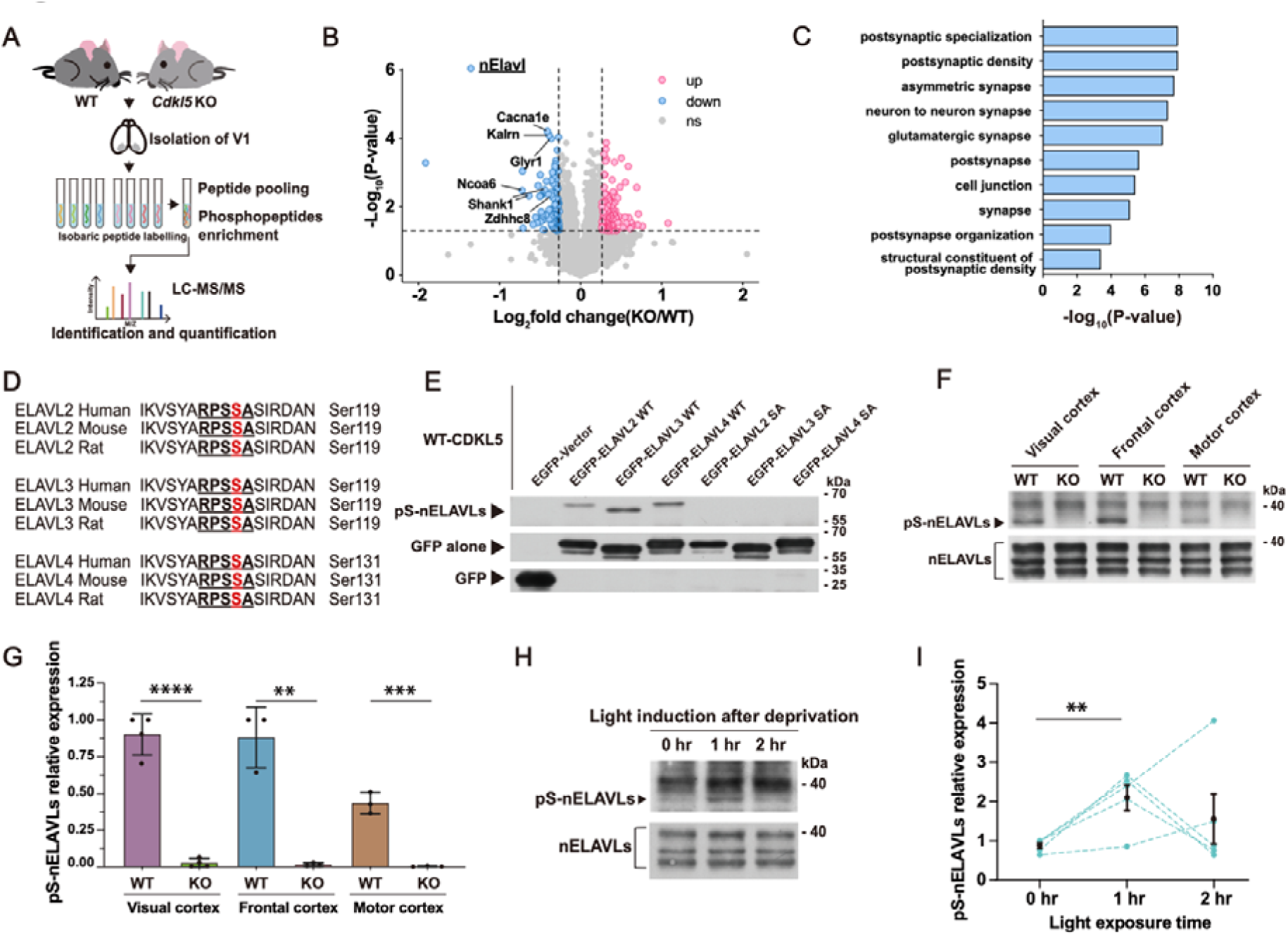
nELAVLs are phosphorylation substrates of CDKL5. (A) Workflow of phosphoproteomic screening in WT and *Cdkl5* KO primary visual (V1) cortices. V1 tissues were harvested for isobaric peptide labelling, followed by peptide pooling and phosphopeptide enrichment. The samples were then quantitatively identified by liquid chromatography-tandem mass spectrometry (LC-MS/MS). (B) Volcano plot showing the log_2_ (fold change) and -log_10_ (p-value) of each gene comparing WT and *Cdkl5* KO mice. Differentially expressed phosphopeptides with fold change > 1.2 and < 0.83 are highlighted in pink and blue, respectively. Dashed lines indicate p < 0.05 and a fold change > 1.2 or < 0.83. Each point represents one peptide, those with the CDKL5 consensus motif (RPXS*A/RPXS*) are labeled with gene name. (C) Gene Ontology (GO) term enrichment of down-regulated differentially expressed genes between WT and *Cdkl5* KO mice. (D) Conserved consensus serine phosphorylation sequence in ELAVL2 (119), ELAVL3 (119) and ELAVL4 (131) in human, mouse and rat. (E) Immunoblotting of immunoprecipitates for phosphorylated serine of ELAVL2, ELAVL3 and ELAVL4 in CDKL5-expressing HEK 293T cell. The mutation of serine to alanine at the RPXS*A motif prevented the phosphorylation. (F & G) Levels of pS-nELAVLs in visual, frontal and motor cortices were virtually undetectable in *Cdkl5* KO mice. Visual: n=4 mice, frontal and motor: n=3 mice. Unpaired t-test (two-tailed). (H & I) Increased pS-nELAVLs in visual cortex in WT mice after dark environment (16hr) with 0hr, 1hr and 2hr light induction. n=5 mice. Unpaired t-test (two-tailed). Dot lines indicate the same batch of experiments. Data are presented as mean ± SEM, ** p < 0.01. *** p < 0.001. **** p < 0.0001.

To validate that CDKL5 can phosphorylate nELAVLs, we generated a phosphospecific antibody targeting these identified sites. We also obtained a wild-type (WT) EGFP-nELAVLs expression construct and generated with the putative serine phosphorylation site replaced by alanine. We confirmed the specificity of nELAVLs antibodies via Western Blot (Figure S1D). Co-expression of CDKL5 and EGFP-ELAVL2, 3 or 4 in HEK 293T cells all showed nELAVL phosphorylation, while phosphorylation was absent when co-expressing CDKL5 and the phospho-deficient mutants (Figure 1E). We also found that nELAVL phosphorylation was consistently reduced in the visual, motor and frontal cortex of Cdkl5 KO mice (Figure 1F & G). These results show that nELAVL proteins are previously unidentified and bona fide CDKL5 phosphorylation targets.

Visual experience plays an essential role in V1 development, including the refinement and matching of input from the contra and ipsi eyes^34^. To test whether phosphorylation of nELAVLs by CDKL5 is experience-dependent, mice were kept in the dark for 16 hours and re-exposed to light for 0, 1 or 2 hours. nELAVLs phosphorylation was significantly higher after 1-hour of light re-exposure but not after 2-hour light re-exposure, suggesting the level of nELAVLs phosphorylation is experience-dependent within a tight time window (Figure 1H & I). Is experience-dependent phosphorylation in V1 specific to nELAVLs, or is this a general feature of CDKL5 substrates? EB2 and MAP1S are two previously reported CDKL5 substrates that are involved in microtubule regulation^10^. We measured the phosphorylation level of MAP1S and EB2 in V1, and found that the phosphorylation level of MAP1S and EB2 were indeed decreased in *Cdkl5* KO mice compared to WT (Figure S1E & G). However, MAP1S and EB2 phosphorylation levels did not show similar changes as nELAVLs phosphorylation after 1-hour and 2-hour light re-exposure, suggesting MAP1S and EB2 phosphorylation in V1 is not experience-dependent *in vivo* (Figure S1F & H).

### CDKL5 deletion alters expression of activity-dependent genes in V1

Given that nELAVLs are RNA binding proteins involved in the regulation of mRNA stability and translation^33^, an important question is how the aberrant phosphorylation of nELAVLs observed in CDKL5 deficiency impacts the expression of relevant mRNAs within mouse V1. Thus, we first used bulk RNA sequencing to profile the transcriptomic differences between WT and *Cdkl5* KO mouse V1. We identified 218 downregulated genes and 272 upregulated genes in *Cdkl5* KO mice when compared to WT mice (Figure 2A & S2A; table S2). Eight selected differentially expressed genes (DEGs) identified by bulk RNA sequencing, including Arc, Fos, Npas4, Pcdh19, Homer1, Gad1, Sst, and Pvalb, were validated using real-time qPCR (Figure S2B). All the DEGs were submitted to STRING to construct a Protein-Protein interaction (PPI) network. The MCC algorithm in CytoHubba plugin was used to identify hub genes in the PPI network. The top ten genes with the highest MCC scores, including *Fos*, *Nr4a1*, *Egr1*, *Fosb*, *Junb*, *Dusp1*, *Egr2*, *Egr3*, *Fosl2* and *Btg2*, were visualized (Figure 2B). These ten genes are all immediate early genes, suggesting that immediate early genes may play an important role in CDD pathophysiology. Consistently, the Gene Set Enrichment Analysis suggests that many activity-dependent neuronal functions, which are known to be driven by visual experience in V1, are impaired in *Cdkl5* KO mice, including regulation of synapse organization, glutamatergic synapses and dendritic spine development (Figure S2C).

**Figure 2.**
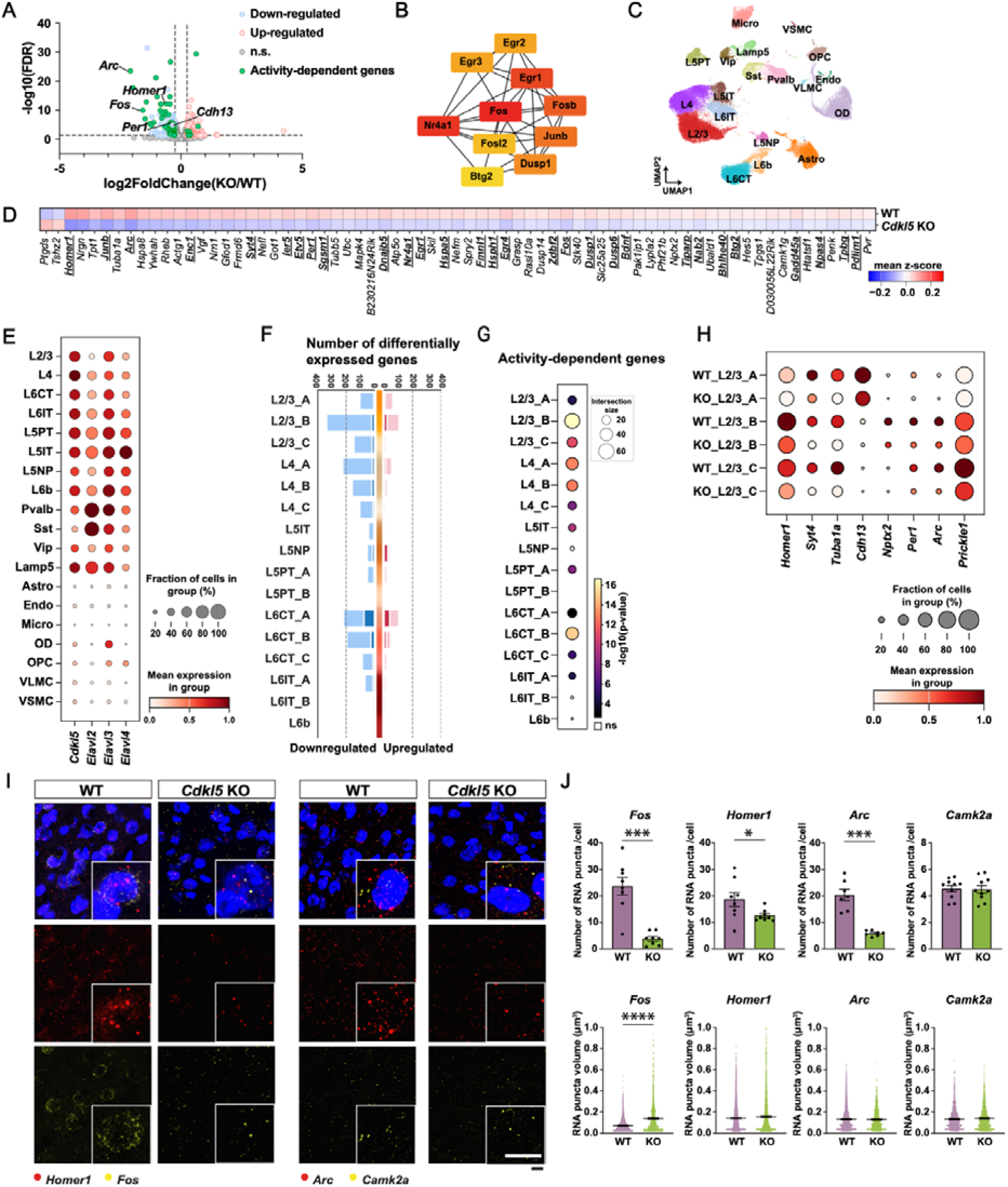
CDKL5 deletion alters activity-dependent gene expression. (A) Volcano plot showing the log_2_ (fold change) and -log_10_ (FDR-adjusted p-value) of each gene comparing WT and *Cdkl5* KO mouse V1. Differentially expressed genes (DEGs) with log_2_ (fold change) >C0.25 and < -0.25 are highlighted in pink and blue, respectively. Activity-dependent genes are highlighted in green. The dashed line indicates FDR-adjusted p-value <C0.05 and log_2_ (fold change) >C0.25 and < -0.25. (B) Hub genes of DEGs between WT and *Cdkl5* KO mouse V1 were identified from PPI network by maximal clique centrality (MCC) algorithm. Edges represent the protein-protein associations. The shading from red to yellow depicts MCC scores from high to low. (C) UMAP plot visualizing annotated subclasses of WT and *Cdkl5* KO mouse V1. (D) Matrix plot showing activity-dependent DEGs identified between WT and *Cdkl5* KO mouse V1. 29 overlapped activity-dependent DEGs between bulk and snRNA sequencing are underlined. (E) Dot plot showing *Cdkl5* and *nElavls* gene expression across subclasses of WT and *Cdkl5* KO mouse V1. (F) Number of DEGs between WT and *Cdkl5* KO mouse V1 in individual cell types of glutamatergic neurons, genes up- and down-regulated in *Cdkl5* KO compared with WT shown in pink and blue bars, respectively. DEGs were defined as genes with |log_2_ fold change| > 0.25 (light colored bars) or > 0.58 (dark colored bars) and FDR-adjusted p-value < 0.05. (G) Dot plot visualizing enrichment results of activity-dependent genes with DEGs. Enrichment was calculated separately for DEGs identified in individual cell types of glutamatergic neurons. Dot size is the number of intersected genes. Color is shaded by -log_10_ (P value) (one-sided Fisher’s exact test). ‘ns’ indicates P valueC≥C0.05. (H) Dot plot visualizing *Homer1, Syt4, Tuba1a, Cdh13, Nptx2, Per1, Arc and Prickle1* gene expression levels in sublayers of Layer 2/3 glutamatergic neurons in WT and *Cdkl5* KO mouse V1. (I) FISH images of mRNA expression of *Fos, Homer1, Arc, Camk2a* in WT and *Cdkl5* KO mouse V1. Scale bars, 20μm. (J) (Top) Bar plot quantifying *Fos, Homer1, Arc, Camk2a* mRNA expression levels. The quantification of mRNA numbers was normalized to the number of DAPI+ cells per image. (Bottom) Quantification of *Fos, Homer1, Arc, Camk2a* mRNA punctum volume. *Fos* and *Homer1*: n=4 mice, *Arc and Camk2a*: n=3 mice. Mann-Whitney test (two-tailed). Data are presented as mean ± SEM, * p < 0.05. *** p < 0.001. **** p < 0.0001.

To validate whether CDKL5 deficiency could lead to dendritic spine deficits in V1, sparsely labeled layer 2/3 and layer 5 excitatory neurons were analyzed by applying epitope-preserving Magnified Analysis of Proteome (eMAP) (Figure S3A-F). In layer 2/3 neurons, we found that the total dendritic spine density of basal and apical dendrites was lower in *Cdkl5* KO mice. In contrast, in layer 5 neurons, we only observed a significant difference in the spine density of basal dendrites (Figure S3G-J). To further investigate the changes in spine morphology, we compared the density of mushroom-shaped, thin, and stubby spines as well as filopodia between WT and *Cdkl5* KO mice. In layer 2/3 neurons, we found that mushroom-shaped spine density was reduced in *Cdkl5* KO mice on both apical and basal dendrites. In layer 5 neurons, only the density of mushroom-shaped spines on basal dendrites decreased in *Cdkl5* KO mice while on apical dendrites, the density of filopodia was higher in *Cdkl5* KO (Figure S3G-J). These findings indicate that *Cdkl5* KO mice have altered spine density and morphology on excitatory neurons, with a more pronounced effect observed in layer 2/3 neurons.

To investigate potential differential effects on specific neuronal layers like layers 2/3 and 5, we conducted single nucleus (sn) RNA sequencing to compare the transcriptomes of WT and *Cdkl5* KO V1. Nuclei from 6 snRNA libraries of V1 were collected from 6 WT and 6 *Cdkl5* KO mice and the gene expression matrices were filtered to remove low-quality cells and cells containing a high proportion of mitochondrial transcripts (>5%). Doublets were identified using Scrublet^35^ and removed in the downstream analysis (Figure S4A) and Harmony^36^ was applied for batch correction (Figure S4B). We obtained 62051 high-quality cells from the 6 snRNA libraries and combined them for the downstream analysis. Dimensionality reduction and Leiden clustering were applied to generate different cell classes, subclasses and types (Figure 2C & S4E-G). Classes consisted of glutamatergic neurons (37289 cells), GABAergic neurons (7318 cells) and non-neuronal cells (17444 cells). The UMAP of overlapping WT and *Cdkl5* KO libraries showed no gross change in cell types (Figure S4C & D).

Next, we performed differential gene expression analysis between the nuclei from WT and *Cdkl5* KO V1. Interestingly, we identified 69 activity-dependent genes among the 458 total DEGs in *Cdkl5* KO mice (Figure 2D; table S3), and ∼50% of these overlapped with activity-dependent DEGs identified in bulk RNA sequencing (Figure S4O), and among these 29 activity-dependent DEGs, 22 are overlapped with nELAVL-binding mRNAs identified previously^33^ (Figure S4P), again suggesting that activity-dependent plasticity might be disrupted by loss of CDKL5. We found that CDKL5, and its phosphorylation targets the nELAVLs, are mainly expressed in glutamatergic neurons (Figure 2E & S4H) which make up the majority of cells in our dataset (Figure S4D). Therefore, we further analyzed glutamatergic subclasses (Figure S4I), and plotted the number of DEGs in the individual glutamatergic cell types (Figure 2F; table S4). L2/3, L4 and L6 cell types have the most DEGs in *Cdkl5* KO mice. We intersected DEGs in different subtypes with activity-dependent genes and found that significant enrichment among DEGs in L2/3 sublayers (Figure 2G).

L2/3 glutamatergic neurons exhibit spatial segregation and display graded gene expression across three sublayers (A, B, and C). L2/3_A is adjacent to the pia, L2/3_C borders L4 and L2/3_B is positioned between L2/3_A and L2/3_C. This segregation represents an activity-dependent maturation of the layer and can be disrupted by dark rearing^37^. The laminar distribution of these sublayer neurons has been suggested to related to different downstream projecting targets and functions^38,39^. Given the deficits in layer 2/3 neurons identified through dendritic spine analysis, we were intrigued to investigate whether transcriptomic maturation of layer 2/3 neurons is influenced by the deletion of CDKL5. Therefore, we separated L2/3 glutamatergic neurons from WT and *Cdkl5* KO V1 into three types (A, B, and C) (Figure S4J & K) according to the sublayer-specific marker *Cdh13*, *Trpc6*, and *Chrm2*^37^. We found that the cell proportions across three sublayers of neurons was unchanged (Figure S4L), but significant transcriptomic alterations were observed. Again, we highlighted several activity-dependent downregulated DEGs (Figure 2H) including *Homer1, Syt4, Tuba1a, Nptx2, Per1, Arc, Prickle1,* as well as the L2/3_A sublayer marker *Cdh13,* which are all documented to associate with neuronal plasticity^40–47^. Among the highlighted DEGs, *Homer1, Syt4, Tuba1a* were found to be common across all sublayers of L2/3. *Cdh13* was uniquely identified as a DEG in sublayer L2/3_A, as expected, while *Nptx2* and *Per1* were specific DEGs in L2/3_B. Arc was found to be significantly downregulated in L2/3_B and L2/3_C whereas *Prickle1* was exclusively identified as a DEG in L2/3_C. These findings suggest distinct molecular signatures associated with each sublayer, indicating potential disruptions in the specialized functional roles within L2/3 sublayers following CDKL5 deletion, without affecting the overall cell proportions of layer 2/3 neurons.

In addition, we also analyzed inhibitory neurons and found their composition also did not show any significant difference between groups (Figure S4M). However, when computing DEGs in different types of inhibitory neurons, we found that most DEGs were specific to Pvalb+ inhibitory neurons (Figure S4N).

To validate whether the activity-dependent DEGs identified in bulk and snRNA sequencing are decreased in *Cdkl5* KO mice, we used fluorescence in-situ hybridization (FISH). We found that mRNA expression levels of *Fos*, *Homer1* and *Arc* were indeed significantly reduced in *Cdkl5* KO mice while *Camk2a*, an excitatory neuron marker, was unaltered (Figure 2I & J). Immunostaining results also showed that the expression of c-fos was decreased significantly in layers 2/3, 4 and 6 in *Cdkl5* KO mice (Figure S4Q & R). Of note, the *Fos* mRNA punctum volume in *Cdkl5* KO mice appeared larger than those in WT mice (Figure 2I & J).

Taken together, the structural abnormalities in dendritic spines (Figure S3) align with the decreased expression of activity-dependent genes and excitatory synaptic mRNA as revealed in our transcriptomic analysis (Figure 2A). Notably, 21 out of 29 dysregulated activity-dependent genes (highlighted in Figure 2D), including *Fos* and *Homer1*, are direct binding targets of nELAVLs identified through CLIPseq analysis^33^. These observations led us to postulate that the CDKL5-mediated phosphorylation of nELAVLs modulates the mRNA levels of activity-dependent genes, thereby influencing the refinement of the visual circuit.

### Loss of CDKL5-mediated phosphorylation alters nELAVLs phase separation

We next sought to investigate how phosphorylation of nELAVLs by CDKL5 affects mRNA expression in mouse V1. While CDKL5 itself was found to undergo phase separation^48^, we therefore evaluate the phase separation propensity of CDKL5’s kinase target nELAVLs. We employed the LLPhyScore algorithm^49^ to analyze a composite dataset comprising these target sequences alongside a benchmark set of experimentally verified LLPS-positive drivers (e.g., FUS, TDP-43, TIA-1) and a broad background of negative control sequences (Figure S5). The comparative ranking analysis indicates that nELAVL proteins possess a significant intrinsic potential for phase separation, distinct from non-phase-separating background proteins. While canonical “scaffold” proteins such as FUS and TDP-43 cluster at the apex of the distribution with near-maximal scores (≈1.0) and top-tier rankings (Rank < 200), the nELAVLs occupy a prominent intermediate-to-high probability tier. nELAVLs exhibit scores ranging from approximately 0.70 to 0.85, positioning them within the top quartile of the distribution (approximate ranks 700–1300) (Figure S5). Although they do not reach the saturation levels of primary drivers, their significant deviation from the tail of the distribution suggests they encode sufficient multivalent interaction valency to participate in phase separation, likely functioning as client proteins recruited into condensates or driving phase transitions under specific physiological conditions rather than acting as constitutive scaffolds. For example, post-translational modification like phosphorylation can finetune the phase separation features of proteins^50^. To examine whether nELAVLs show different protein distribution with or without CDKL5, we employed expanded V1 tissue of WT and *Cdkl5* KO mice by eMAP. We immunolabelled nELAVLs using a pan-nELAVLs antibody and ELAVL3 using a ELAVL3 specific antibody available (Figure S1D). We found that the immunolabelled signals formed cytoplasmic puncta in V1 neurons (Figure 3A). Although the number of puncta in each neuron showed no significant change, the volume of these puncta were significantly increased in volume in *Cdkl5* KO V1 neurons (Figure S6A). By fraction analysis, we also showed that the volume fraction distribution of these puncta in each neuron has significantly right shifted in *Cdkl5* KO mice, providing evidence that each of the KO V1 neurons contains larger nELAVL punta as compared to WT V1 neurons (Figure 3B). A similar phenotype was observed in the *CDKL5* knockdown primary neurons (Figure S6B & C). These unexpected findings suggested that nELAVLs are involved in droplet-like puncta in cells, and the punctum size is regulated by phosphorylation of nELAVLs by CDKL5.

**Figure 3.**
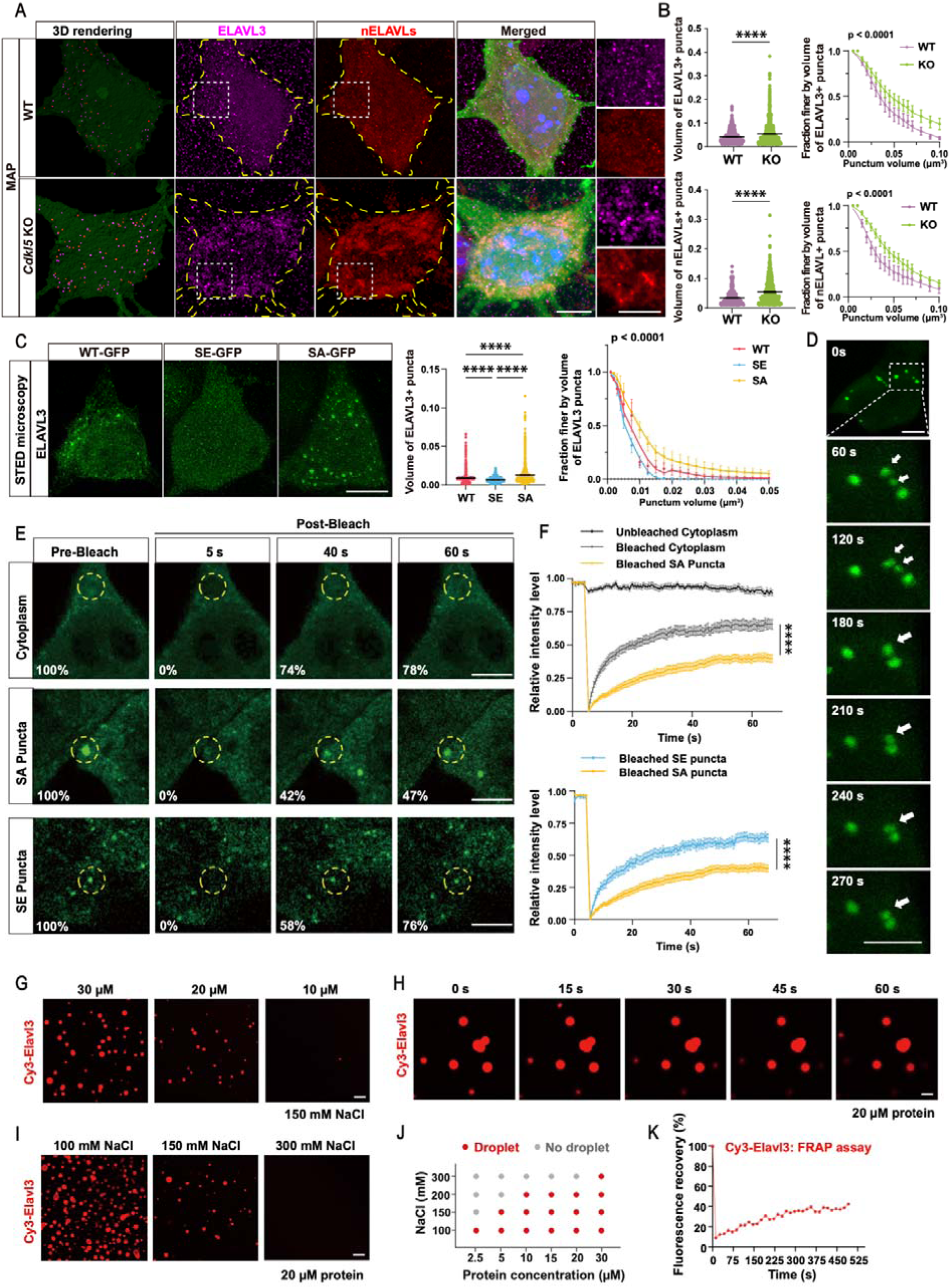
Loss of phosphorylation by CDKL5 alters nELAVL phase separation. (A & B) With epitope-preserving Magnified Analysis of Proteome (eMAP), ELAVL3 (magenta) and nELAVL (red) puncta in *Cdkl5* KO mice showed large-shifted punctum volume distribution among neurons. Scale bar, 10 μm and 3 μm for zoomed-in images (right) of ELAVL3 and nELAVL. WT: n=21 neurons from 3 mice; *Cdkl5* KO: n=27 neurons from 3 mice. Left: Mann-Whitney test (two-tailed); Right: Two-way ANOVA with Sidak’s multiple comparisons test on main effect of WT and *Cdkl5* KO. (C) Super-resolution STED microscopy of overexpressed WT and SE/SA mutant of ELAVL3 in rat primary hippocampal neurons. Compared to SE, SA mutant showed large-shifted punctum volume distribution among neurons. Scale bar, 5 μm. n=4 cells from 3 batches of experiments. Left: One-way ANOVA followed by Turkey’s multiple comparisons test; Right: Two-way ANOVA with Sidak’s multiple comparisons test on main effect of SE and SA. (D) Time-lapse images showing that ELAVL3 droplets fused with each other over time in ELAVL3-SA-overexpressing HEK 293T cells. Scale bar, 3 μm. (E) Confocal images showing fluorescence recovery between ELAVL3-SA and -SE puncta. Bleaching area radius ∼ 2.5 μm. Images represent the ELAVL3 puncta before bleaching and right after bleaching (5s, 40s and 60s). Scale bar, 5μm (F) Fluorescence recovery rate of bleached SA puncta was slower than those of bleached cytoplasm or bleached SE puncta. n=20 (unbleached cytoplasm), 12 (bleached cytoplasm), 24 (bleached SA puncta) and 17 (bleached SE puncta) cells from 3 batches of experiments. Two-way ANOVA with Sidak’s multiple comparisons test. (G) Purified full length ELAVL3 protein underwent phase separation at indicated concentrations. ELAVL3 was sparsely labelled by Cy3 fluorophore at 1%. Scale bar, 5μm. (H) Time-lapse images showing that ELAVL3 droplets fused with each other over time. Scale bar, 5 μm. (I) The number of ELAVL3 droplets decreased when NaCl concentration increased. Scale bar, 5 μm. (J) Phase separation diagram showing phase separation properties within demonstrated NaCl concentration and ELAVL3 concentration. The highlighted red dots: phase separation; grey dots: no phase separation. (K) Fluorescence recovery analysis showing that ELAVL3 in condensed droplets dynamically exchanged with those in the dilute phase. Data are presented as mean ± SEM, **** p < 0.0001.

To test the dependence of droplet-formation on nELAVL phosphorylation by CDKL5, we employed phospho-mimetic and phospho-deficient mutations of nELAVLs (SE and SA mutants with the consensus CDKL5 phosphorylation motif of -RPSSA-was substituted with -RPSEA- and -RPSAA-, respectively), to evaluate their capacity to form the droplet-like puncta. We overexpressed GFP-tagged nELAVL-SE and nELAVL-SA in both primary neurons and HEK 293T cells (Figure 3C & S6D, G & I). We first verified that the exogenous nELAVL proteins showed equal expression by immunoblotting (Figure S6E). Subsequently, through stimulated emission depletion (STED) microscopy, we visualized the cellular distribution of nELAVL proteins and showed that the volume of ELAVL3-SA and ELAVL4-SA puncta were significantly larger than that of ELAVL3-SE and ELAVL4-SE punta, respectively, in both primary neurons and HEK 293T cells (Figure 3C & S6F-J), supporting a critical role of CDKL5-mediated phosphorylation in regulating nELAVL punctum size.

The liquid-like nature of the nELAVL puncta was corroborated through live-cell imaging. As we observed similar droplet-like puncta among three nELAVLs, we selected ELAVL3 as a key example to demonstrate their biophysical property. Smaller ELAVL3-SA puncta were found to rapidly coalesce into larger ones over time, suggesting a liquid-like nature (Figure 3D). To evaluate the fluid-like property of the nELAVL puncta, fluorescence recovery after photo-bleaching (FRAP) were performed in HEK 293T expressing ELAVL3 SA and SE mutants (Figure 3E & F). The FRAP assays showed that both the SA and SE mutants within these puncta were capable of exchanging with the surrounding cytoplasm. It is worth noting that the recovery rate of SA was much slower than that of SE; within 40 seconds, the recovery rate of SA was approximately 35%, while that of SE was around 60% (Figure 3E & F). These data indicated that nELAVL is a member of cytosolic condensates and that its phosphorylation state modulates the properties of this condensate.

We next purified the recombinant full-length ELAVL3 protein and sparsely labeled the purified ELAVL3 with the Cy3 fluorophore. Cy3-labeled ELAVL3 protein formed spherical droplets in solution in a concentration-dependent manner (Figure 3G). These droplets fused quickly with each other over time (Figure 3H). The number of ELAVL3 droplets decreased when the salt concentration was elevated (Figure 3I), indicating that electrostatic interaction may contribute to the droplet’s formation. The phase diagram demonstrated that the droplets could form at the minimal concentration of 2.5μM, which is close to physiological condition (Figure 3J). The *in vitro* FRAP experiment showed that the ELAVL3 droplets exhibited recovery subsequent to photobleaching through protein exchange between the condensed droplets and surrounding solution (Figure 3K), suggesting that the ELAVL3 droplets possess properties characteristic of a phase separated state.

Taken together, these biophysical features of nELAVLs are reminiscent of biomolecular condensates in biological systems, indicating that nELAVLs can undergo phase separation *in vitro* and in cells. Moreover, their phase separation characteristics can be modulated by the CDKL5-dependent phosphorylation.

### CDD patient iPSC-derived neurons form aberrant nELAVLs condensates and altered crosstalk with P-bodies and stress granules

To investigate whether nELAVL condensates behavior is altered in the context of CDD, we aimed to characterize the nELAVL condensates in female patient-derived iPSC lines carrying the pathogenic variant 1648C>T (p.R550*) of CDKL5. As controls, we utilized a WT allele-expressing control line from the same patient, as well as another base editing-corrected isogenic control line (Figure 4A). We confirmed that CDKL5 expression was absent in the R550* line-derived neurons (iNeurons) (Figure 4B), rendering it a comparable model to KO. CDKL5 expression was restored in the isogenic control line (Figure 4C). As expected, the phosphorylation levels of nELAVLs were significantly reduced in R550* iNeurons and rescued in the isogenic line (Figure 4D & E). STED images of iNeurons revealed that the volume of ELAVL3 and nELAVL condensates were both significantly increased in the R550* iNeurons than in the control line, and this phenotype was rescued in the isogenic line (Figure 4F & G & S7A). These results demonstrated that nELAVL condensates is regulated by CDKL5-mediated phosphorylation in human CDD iNeurons, suggesting that our observations from CDKL5 deficient mouse neurons are translatable.

**Figure 4.**
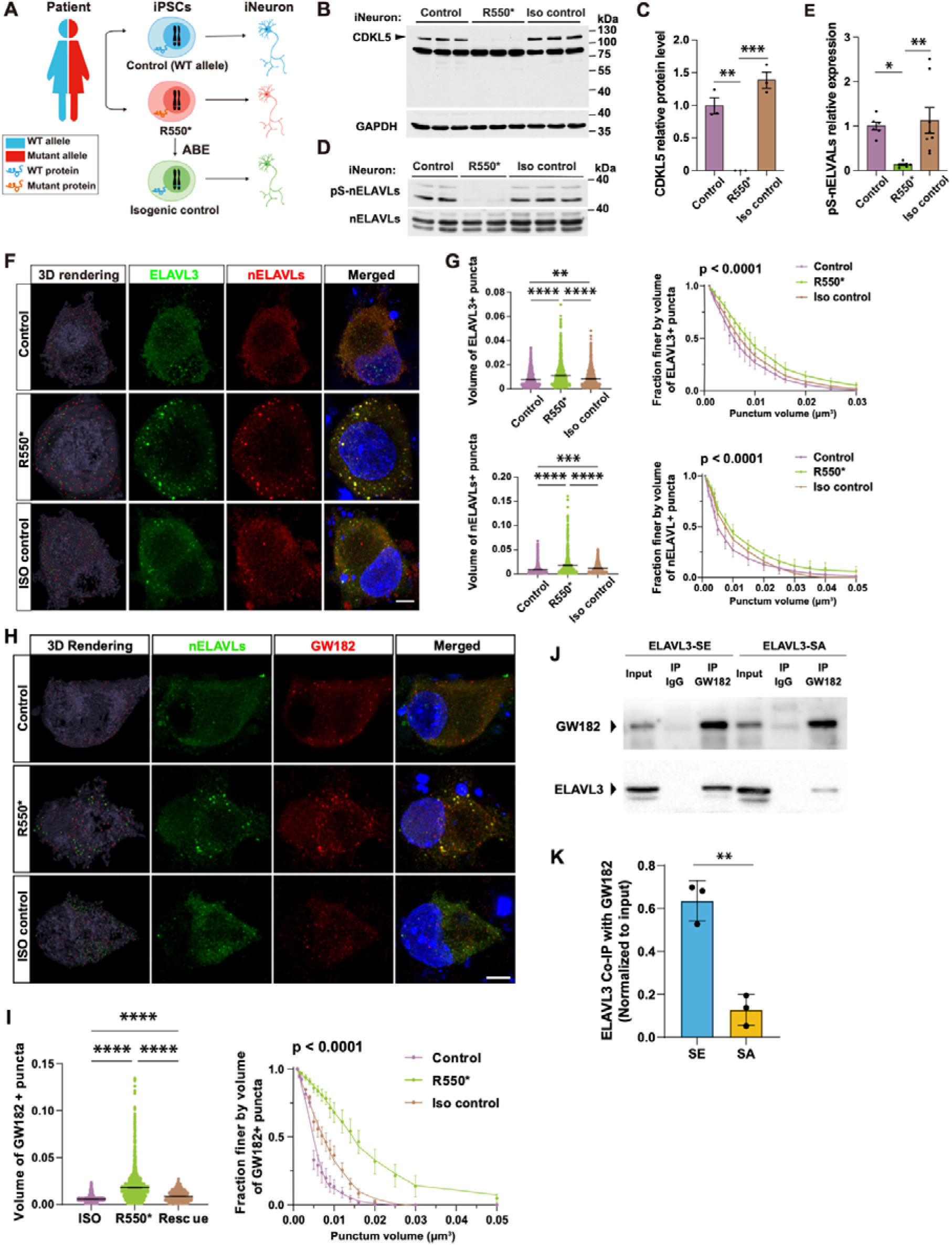
CDD patient iPSC-derived neurons forms aberrant nELAVL condensates and altered crosstalk with P-bodies. (A) An illustration of CDD patient iPSC-derived cortical neurons (iNeurons). iPSCs lines: R550* = CDKL5 deficient, control = wildtype (WT) and isogenic control = R550* that has undergone adenine base editing (ABE) to restore CDKL5 expression. (B & C) CDKL5 expression levels were comparable between control and isogenic control iNeurons but absent in R550* iNeurons. n=3. One-way ANOVA followed by Tukey’s multiple comparisons test. (D & E) pS-nELAVLs levels were comparable between control and isogenic control iNeurons but reduced in R550* iNeurons. Control: n=6, R550*: n=7, isogenic control: n=8. One-way ANOVA followed by Tukey’s multiple comparisons test. (F & G) STED microscopy showed large-shifted punctum volume distribution of ELAVL3 (green) and nELAVLs (red) in R550* iNeurons, in comparison to control and isogenic control iNeurons. Scale bar, 5 μm. n=6 cells from 3 batches of experiments. Left: One-way ANOVA with Tukey’s multiple comparisons test; Right: Two-way ANOVA with Sidak’s multiple comparisons test on main effect of Control, R550* and Iso control; Multiple comparisons: top: p < 0.0001 for Control vs R550*, p < 0.0001 for R550* vs Iso control; bottom: p < 0.0001 for Control vs R550*, p < 0.0001 for R550* vs Iso control. (H & I) STED microscopy showed large-shifted punctum volume distribution of GW182 (red) in R550* iNeurons, in comparison to control and isogenic control iNeurons. Scale bar, 5μm. n=6 cells from 3 batches of experiments. Left: One-way ANOVA with Tukey’s multiple comparisons test; Right: Two-way ANOVA with Sidak’s multiple comparisons test on main effect of Control, R550* and Iso control; Multiple comparisons: top: p < 0.0001 for Control vs R550*, p < 0.0001 for R550* vs Iso control; bottom: p < 0.0001 for Control vs R550*, p < 0.0001 for R550* vs Iso control. (J) Co-immunoprecipitation of GW182 and ELAVL3-SE/SA from cytoplasmic extracts of HEK 293T cells. Immunoprecipitation was performed using an anti-GW182 antibody. ELAVL3 bands were revealed with an anti-ELAVL3 antibody. (K) Quantification results of Co-immunoprecipitation, the GW182 capture more ELAVL3-SE than ELAVL3-SA. n=3 batches of experiments. Unpaired t-test (two-tailed). Data are presented as mean ± SEM, * p < 0.05; ** p < 0.01; *** p < 0.001; **** p < 0.0001.

RNP granules have emerged as crucial regulators of many cellular processes, including transcription and translation^51^. Among the RNP granules, P-bodies and stress granules are well documented to regulate RNA processing and are dysregulated in human diseases, including several neurodevelopmental disorders^52^. In addition, aberrant RBPs co-condensation has been reported in neurodegenerative conditions^14^. We therefore asked whether dysregulated nELAVL condensates may trigger aberrations in other RNP condensates, such as P-bodies and stress granules to influence RNA processing. To test this, we co-immunolabelled nELAVLs with P-body marker GW182 and stress granules marker TIA1 in CDD iNeurons. We found that P-bodies and stress granules volumes were significantly increased in R550* group and such abnormal increase was rescued in isogenic control group (Figure 4H & I & S7B-D).

To further investigate the impact of nELAVL phosphorylation on P-bodies and stress granules dynamics, we co-expressed ELAVL3-SA with GW182 or TIA1 in HT22 cells (Figure S7E & G). Similar to the effects observed in iNeurons, the expression of ELAVL3-SA led to an increased size of GW182 and TIA1 puncta, while the expression of ELAVL3-SE led to decreased size (Figure S7E-H). We next hypothesized that the phosphorylation status of ELAVL3 modulates its interaction with GW182. To test this, we performed co-immunoprecipitation and found that the interaction between GW182 and ELAVL3-SE was significantly stronger than that with the phospho-dead ELAVL3-SA mutant (Figure 4J & K). These results suggest that phosphorylation of nELAVLs not only mediates the formation of their own condensates but also modulates their interaction with other biomolecular condensates, such as P-bodies.

### CDKL5 modulate mRNA processing by regulating nELAVL condensates

P-bodies and stress granules are RNP granules known to regulate mRNA storage, decay, and translation^52^, and dynamically sequester mRNAs as a regulatory site during cellular processes^53–55^. To study the impact of nELAVL phosphorylation on RNA processing, we employed FISH assay and puro-PLA assay to examine the mRNA level and translation of *Fos*, a known binding target of nELAVLs^56^. We found reduced *Fos* mRNA puncta in ELAVL3-SA expressed primary neurons compared to ELAVL3-SE expressed neurons (Figure 5A& B). From the puro-PLA assay, we dual-labelled puromycin and Fos protein after the puromycin treatment and found reduced *Fos* translation in ELAVL3-SA expressed primary neurons compared to ELAVL3-SE expressed neurons (Figure S7I & J).

**Figure 5.**
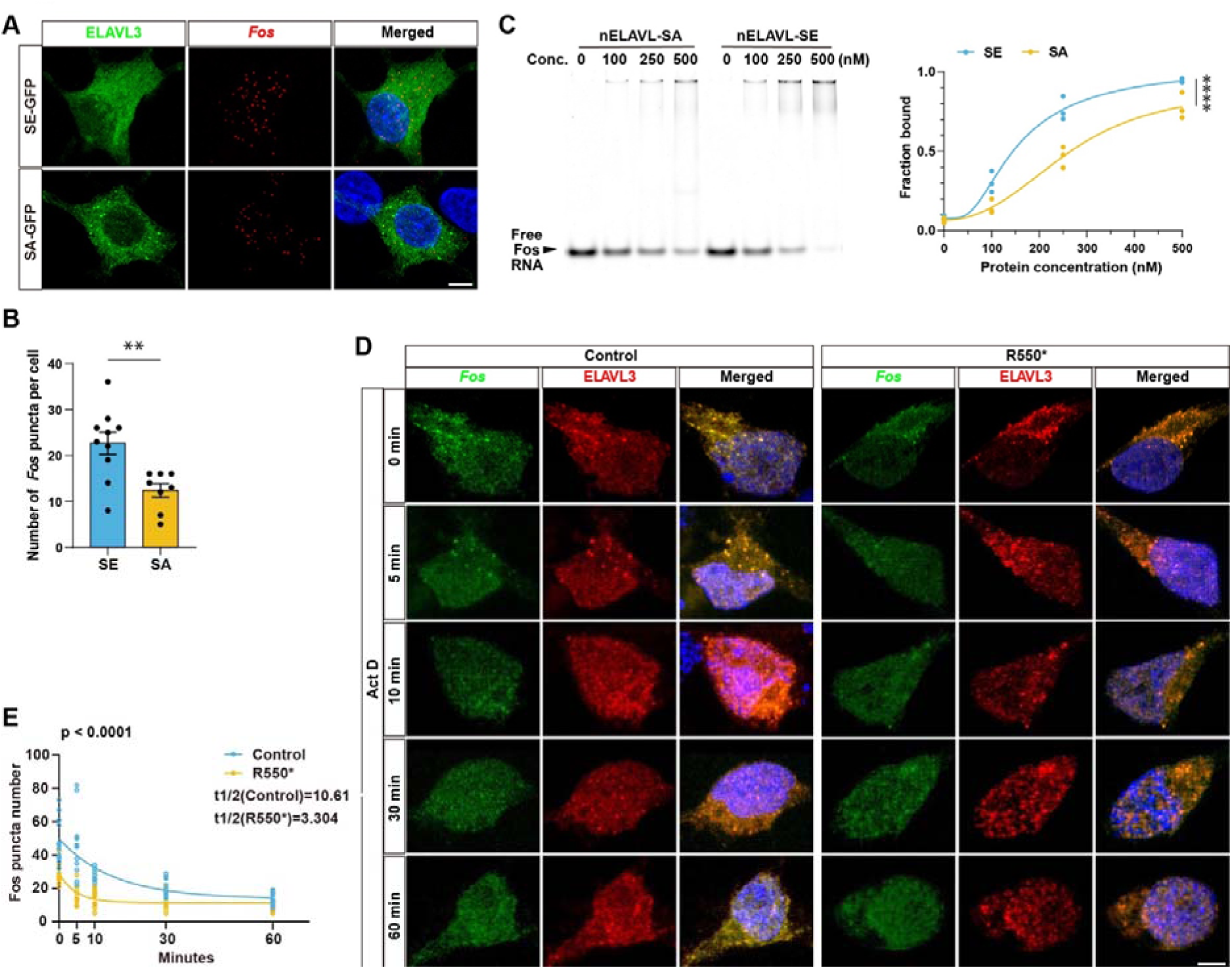
CDKL5 modulate mRNA processing by regulating nELAVL condensates. (A & B) FISH images of mRNA expression of *Fos* in ELAVL3-SE/SA expressing rat primary neurons. Compared to SE, ELAVL3-SA expressing neurons showed less *Fos* RNA puncta. Scale bar, 5μm. n=10 (SE) and 8 (SA) from 4 batches of experiments. Mann-Whitney test (two-tailed). (C) EMSA showing different concentration of nELAVL-SE and -SA protein binding with *Fos* RNA. At higher protein concentration, the *Fos* RNA bound fraction in SE group is significantly higher than SA group. n=3. Two-way ANOVA with Sidak’s multiple comparisons test. (D & E) FISH images of mRNA expression of *Fos* in Control and R550* iNeurons after 2 μg/mL Actinomycin D (Act D) treatment. Compared to control group, CDD iNeurons showed shorter *Fos* mRNA half-life. Scale bar, 5μm. n=12 iNeurons from 3 batches of experiments. Two-way ANOVA with Sidak’s multiple comparisons test on main effect of Control and R550*. Data are presented as mean ± SEM, ** p < 0.01; **** p < 0.0001.

Cell-free Electrophoretic Mobility Shift Assay (EMSA) reveals different binding affinities between nELAVL-SE and SA to *Fos* RNA. The recombinant nELAVL proteins formed protein-RNA complexes with *Fos*-Cy5 RNA (Figure 5C). The binding affinity of nELAVL-SE to *Fos* was significantly higher than that of nELAVL-SA to *Fos*, indicating that the phosphorylation status of nELAVLs modulates their binding affinity to target RNA (Figure 5C).

To test whether the mRNA stability was altered, we used Actinomycin D to inhibit the transcription at different time point (0, 5, 10, 30, 60 mins) before FISH assay on iNeurons. We found that in R550* iNeurons, the half-life of *Fos* mRNA was significantly shorter than in Control iNeurons (Control: 10.61 vs. R550*: 3.304), suggesting increased mRNA degradation in CDD iNeurons (Figure 5 D & E).

Taken together, these findings provide evidence for a mechanistic link between CDKL5-mediated nELAVL phosphorylation and the formation and function of RNP granules which plays important role in mRNA processing.

### CDKL5 deletion impairs visually guided behavior and visually evoked functional responses

To determine whether the *Cdkl5* KO mice exhibits behavioral deficits related to cortical visual impairment, we employed the visual cliff task^57,58^, which is designed to detect impairments in stereoscopic depth perception. In this task, animals were placed on a central platform within an apparatus containing two distinct areas: a “safe side” and a “cliff side” (the later consisting of a glass plate positioned above a sharply dropped, patterned floor) (Figure 6A, left). WT mice with intact stereoscopic vision consistently avoided the “cliff” side, descending approximately 80% from the “safe” side (Figure 6A, right). On the other hand, we found that *Cdkl5* KO mice descended significantly more times from the ‘cliff’ side, suggesting impaired stereoscopic depth perception (Figure 6A, right).

**Figure 6.**
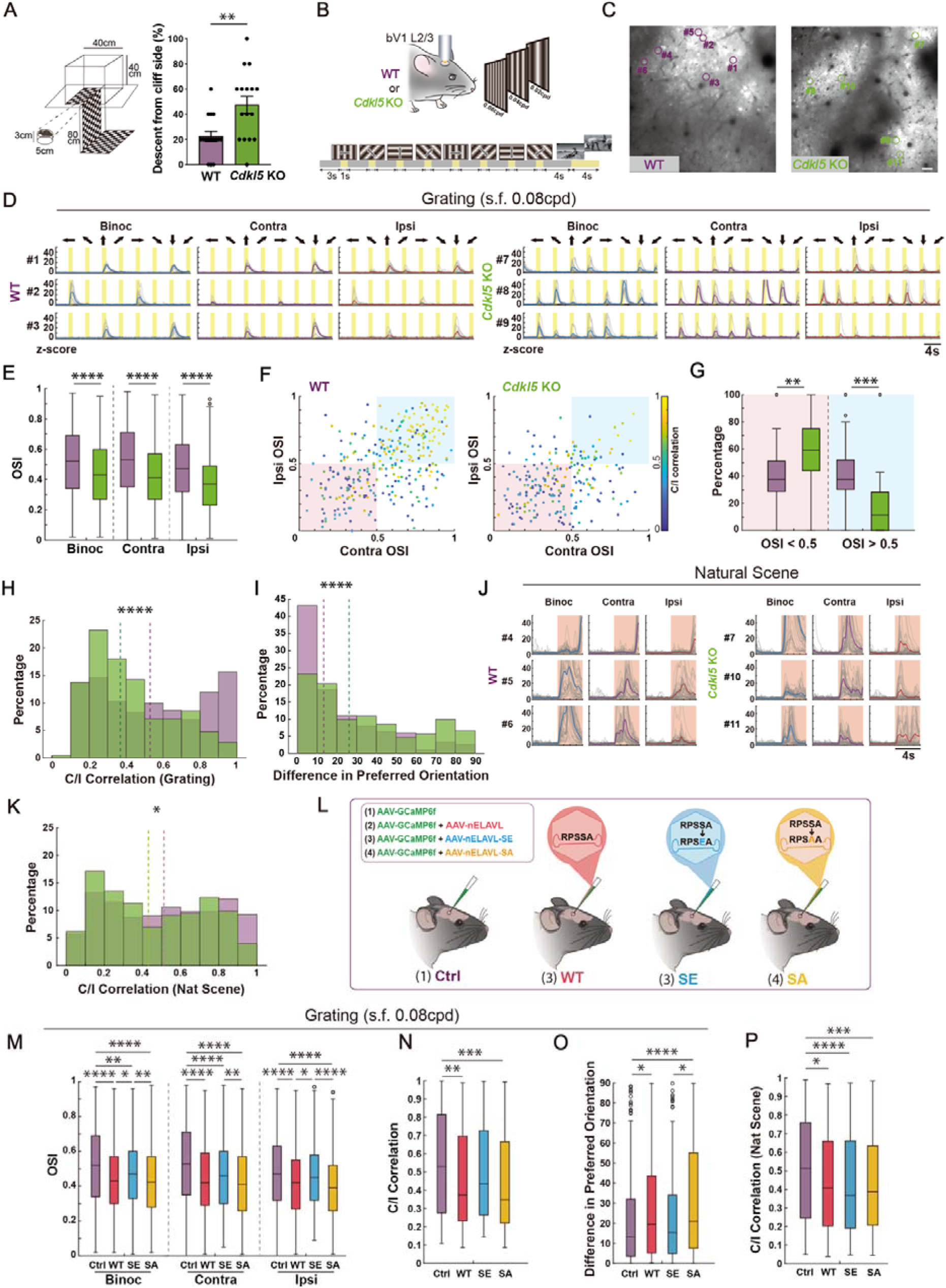
CDKL5 deletion leads to impaired visual function and nELAVL phosphorylation is necessary for visual processing. (A) Graphical design of the visual cliff task with measurements. Animals were placed on a platform (3 cm height) in the middle of the open field (left). Percentage of descents from the cliff side per mouse (5 trials per mouse) between WT and Cdkl5 KO mice (right). WT: n=16 mice, *Cdkl5* KO: n=16 mice. Mann-Whitney U test (two-tailed). (B) Head-fixed wildtype control (WT) mice and *Cdkl5* KO were presented, either to one eye or both eyes, with visual stimuli of different directional grating patterns. In the binocular visual cortex (bV1), calcium signal in layer 2/3 (L2/3) neurons were live-imaged with two-photon laser microscopy. (C) Field of views (FOVs) showing grating and natural scene responsive neurons in WT (left) and *Cdkl5* KO (right) mice. Scale bar, 20 μm. (D) Traces showing grating responsive neurons in WT and *Cdkl5* KO mice across 3 viewing conditions, which are binocular (Binoc), contralateral (Contra) and ipsilateral (Ipsi) viewing relatively to the brain hemisphere imaged. Yellow regions indicate grating presentation periods. Gray lines represent the Z-score from all trials and the colored lines show trial average. (E) Boxplots showing the orientation selective index (OSI) of *Cdkl5* KO mice were reduced across Binoc, Contra and Ipsi conditions under grating (s.f. 0.08cpd). Binoc OSI: 870 neurons from 6 WT and 698 neurons from 6 *Cdkl5* KO mice. Contra OSI: 794 neurons from 6 WT and 693 neurons from 6 *Cdkl5* KO. Ipsi OSI: 629 neurons from 6 WT and 476 neurons from 6 *Cdkl5* KO. The centerlines represent median values, and the whiskers connect the nonoutlier minimum and maximum values to 0.25 and 0.75 quartiles respectively. Outliers are values greater than 1.5 interquartile range away from the quartiles. Mann-Whitney U test (two-tailed). (F) Scatter plots comparing OSI and correlation of grating (s.f. 0.08cpd) responsive neurons in WT mice and *Cdkl5* KO mice. Color bar shows the correlation between tuning curves. 301 neurons from 6 WT and 211 neurons from 6 *Cdkl5* KO. Mann-Whitney U test (two-tailed). (G) Boxplots showing fraction of neurons per FOV within the blue region (OSI > 0.5) and red region (OSI < 0.5) from E. 26 FOV from 6 WT mice and 21 FOV from 6 *Cdkl5* KO mice. Mann-Whitney U test (two-tailed). (H) Distribution of Contra and Ipsi eye correlation between WT and *Cdkl5* KO under grating (s.f. 0.08cpd) stimuli. 301 neurons from 6 WT and 211 neurons from 6 *Cdkl5* KO. Mann-Whitney U test (two-tailed). (I) Distribution of difference in preferred orientation between WT and *Cdkl5* KO under grating (s.f. 0.08cpd) stimuli. 301 neurons from 6 WT and 211 neurons from 6 *Cdkl5* KO. Mann-Whitney U test (two-tailed). (J) Traces showing natural scene responsive neurons in WT and *Cdkl5* KO mice across 3 viewing conditions. Pink regions indicate natural scene presentation periods. Gray lines represent the Z-score from all trials and the colored lines show trial average. (K) Distribution of contra and ipsi eye correlation between WT and *Cdkl5* KO under natural scene stimuli. 529 neurons from 6 WT and 274 neurons from 6 *Cdkl5* KO. Mann-Whitney U test (two-tailed). (L) Mice were virally injected to express GCaMP6f (Ctrl), GCaMP6f and nELAVL (WT), GCaMP6f and nELAVL-SE (SE) or GCaMP6f and nELAVL-SA (SA). Head-fixed mice were presented, either to one eye or both eyes, with visual stimuli of different directional grating patterns and natural scene. In the binocular visual cortex (bV1), calcium signal in layer 2/3 (L2/3) neurons were live-imaged with two-photon laser microscopy. (M) Boxplots showing OSI were reduced in WT, SE and SA mice across Binoc, Contra and Ipsi conditions under grating (s.f. 0.08cpd). Binoc OSI: 870 neurons from 6 Ctrl, 670 neurons from 5 WT injected, 532 neurons from 4 SE injected and 782 neurons from 5 SA injected mice. Contra OSI: 794 neurons from 6 Ctrl, 715 neurons from 5 WT injected, 500 neurons from 4 SE injected and 852 neurons from 5 SA injected mice. Ipsi OSI: 629 neurons from 6 Ctrl, 574 neurons from 5 WT injected, 384 neurons from 4 SE injected and 613 neurons from 5 SA injected mice. Kruskal-Wallis test followed by Tukey’s multiple comparisons test. (N) Boxplots showing Contra and Ipsi correlation under grating (s.f. 0.08cpd) stimuli in Ctrl, WT injected, SE injected and SA injected mice. 301 neurons from 6 Ctrl, 253 neurons from 5 WT injected, 159 neurons from 4 SE injected and 275 neurons from 5 SA injected mice. Kruskal-Wallis test followed by Tukey’s multiple comparisons test. (O) Boxplots showing difference in preferred orientation under grating (s.f. 0.08cpd) stimuli in Ctrl, WT injected, SE injected and SA injected mice. 301 neurons from 6 Ctrl, 253 neurons from 5 WT injected, 159 neurons from 4 SE injected and 275 neurons from 5 SA injected mice. Kruskal-Wallis test followed by Tukey’s multiple comparisons test. (P) Boxplots showing Contra and Ipsi correlation under natural scene stimuli in Ctrl, WT injected, SE injected and SA injected mice. 529 neurons from 6 Ctrl, 185 neurons from 5 WT injected, 428 neurons from 4 SE injected and 443 neurons from 5 SA injected mice. Kruskal-Wallis test followed by Tukey’s multiple comparisons test. Data are presented as mean ± SEM, * p < 0.05. ** p < 0.01. *** p < 0.001. **** p < 0.0001.

To investigate how CDKL5 deletion influences neuronal responses and computations in the mouse binocular visual cortex (bV1), we conducted two-photon calcium imaging of layer 2/3 neurons expressing the calcium indicator GCaMP6f. The visual stimuli were drifting sinusoidal gratings (8 directions, with 45° increments^59^, 0.08, 0.04 and 0.02 cycle per degree (cpd) at 2Hz temporal frequency) with a 3-second grey screen between grating stimuli, followed by 4-second of natural scenes (Figure 6B). During imaging, mice viewed drifting grating and natural scenes stimuli presented independently to each eye (contralateral [contra] or ipsilateral [ipsi] only eye viewing) or to both eyes (binocular viewing) (Figure 6C & D & J). Consistent with established findings, we observed bV1 neurons exhibited varying responses and preferences for stimulus direction to binocular, ipsi eye, or contra eye viewing (Figure 6C & D & J)^60,61^. WT and *Cdkl5* KO mice did not differ in their percentage of binocularly responsive or monocularly responsive neurons (Table S5). Individual neurons in V1 are often preferentially activated by specific orientations of grating. Therefore, we next measured these preferences, quantified as an orientation selectivity index (OSI). *Cdkl5* KO mice showed robust and significant decrease in OSI across all three imaging conditions (binocular, contra eye only and ipsi eye viewing) at high spatial frequency (0.08cpd) as compared to WT mice, suggesting that they are overall less selective (Figure 6E). A significant difference in OSI was observed across all three tested spatial frequencies, with 0.08cpd having the largest OSI decrease in *Cdkl5* KO mice compared to WT (Figure 6E, S8A & K). There is no difference in neuron response amplitude between WT and *Cdkl5* KO mice except ipsi eye viewing condition with 0.02cpd (Table S6).These results suggested that the cortical visual deficits in *Cdkl5* KO V1 neurons are more sensitive to higher spatial frequency (Figure S8K).

Neurons in bV1 that receive inputs from both eyes (contra+ipsi responsive) match specific visual responses from each eye to integrate their visual information, thereby establishing an accurate representation of binocular visual space. The binocular neurons are crucial to visual functions such as disparity and depth perception^62^, and the matching of visual information is highly sensitive to disruption of visual experience or plasticity in development^63,64^. We therefore examined if binocular matching was altered upon CDKL5 deletion. We first examined the relationship between OSI and orientation matching (measured by pairwise signal correlations of their calcium responses between the contra eye and ipsi eye) in WT mice (Figure 6F, left). We observed that neurons with a high OSI (OSI>0.5) in both their contra eye and ipsi eye responses exhibited stronger contra-ipsi orientation matching (blue region in Figure 6F, left). On the other hand, neurons with a low OSI (OSI<0.5) in both their contra eye and ipsi eye responses exhibited weaker contra-ipsi orientation matching (red region in Figure 6F, Left), consistent with published findings^60,65^. In *Cdkl5* KO mice, we observed a significantly reduced fraction of neurons with high OSI and high contra-ipsi orientation matching (blue region in Figure 6F & G) and an increased fraction of neurons with low OSI and low contra-ipsi orientation matching (red region in Figure 6F & G) as compared to WT mice. This suggested that there was a specific impairment of ipsi eye alignment to contra and binocular orientation preferences in *Cdkl5* KO bV1 neurons. Additionally, we examined the difference in preferred orientation between contra and ipsi responses among binocular neurons in WT and *Cdkl5* KO mice. Of note, contra-ipsi (C/I) response correlation and difference in preferred orientation between contra and ipsi responses are two negatively correlated measurements to examine binocular matching (Figure S8B). We observed that *Cdkl5* KO mice showed significant decrease in C/I responses correlation and significant increase in difference in preferred orientation in grating stimuli at spatial frequency 0.08cpd (Figure 6H & I). Consistent with the OSI results, a milder binocular matching deficit was observed in *Cdkl5* KO bV1 neurons at spatial frequency 0.04cpd (Figure S8C-F), and no significant difference was observed at 0.02cpd (Figure S8G-J). In addition, bV1 neurons in *Cdkl5* KO mice exhibited decreased C/I correlation in response to natural scene stimuli, which represent a more complex visual stimulus class (Figure 6K). In sum, the deletion of CDKL5 results in decreased orientation selectivity and contra-ipsi response matching in the bV1 neurons. These deficits suggest there is impaired integration of ipsi eye input, which matures after contra eye inputs are already established and is experience dependent^66^.

### CDKL5 deletion disrupts circuit-wide visual information processing

To investigate how these single-neuron changes affected the binocular circuit, we computed the information encoding at the network level. We applied a supervised support vector machine (SVM) decoder to estimate the information encoding at a populational level, discriminating all four orientations or two natural scenes in our imaging paradigm. The decoding accuracy in the *Cdkl5* KO mice was consistently lower than in the WT mice at fixed population sizes (Figure S8M), suggesting a reduction in visual information in *Cdkl5* KO mice. To evaluate the binocular matching in the population level, we also trained a decoder based on contra only responses and applied this decoder to the ipsi only responses from the same neuronal populations (Figure S8L). At a spatial frequency of 0.08 and 0.04cpd, contra trained decoding accuracy of ipsi responses was significantly lower in *Cdkl5* KO mice than in WT mice, suggesting that visual information contained in the contra responses was less well matched to ipsi responses in *Cdkl5* KO mice. Thus, the deletion of CDKL5 impairs the encoding of visual information at network level, with more pronounced deficits in ipsi eye responses and matching.

### Fine balance of nELAVLs phosphorylation level is necessary for normal visual processing

Finally, to determine if cortical visual deficits in *Cdkl*5 KO mouse bV1 could be recapitulated by altering nELAVL phosphorylation, we again performed *in vivo* two-photon calcium imaging in layer 2/3 neurons in the bV1 of WT mice injected with virus containing the nELAVL-WT (WT), the nELAVL-SE phospho-mimetic (SE) or nELAVL-SA phospho-deficient (SA), and also compared with the group without expressing exogenous nELAVL proteins (Ctrl) (Figure 6L & S9A). We found no change in the overall proportion of responsive neurons across the different viewing conditions (Table S5). However, OSI was decreased in each viewing condition for nELAVL-WT, SE and SA injected mice when compared to Ctrl, with OSI lower in SA than in SE injected mice for all viewing conditions in grating stimuli at spatial frequency 0.08, 0.04 and 0.02cpd (Figure 6M & S9B & E), while neuron response amplitude is no difference between SE and SA injected mice (Table S6). Intriguingly, WT injected mice showed higher OSI than SA mice but lower than the OSI of SE mice, suggesting nELAVL phosphorylation level modulates orientation selectivity. We next examined binocular matching in these mice. We observed that both WT and SA injected mice showed decrease in C/I response correlation and increase in difference in preferred orientation in grating stimuli at spatial frequency 0.08cpd, as compared to Ctrl (Figure 6N & O). On the other hand, no significant difference was observed between SE-injected mice and Ctrl group (Figure 6N & O). These suggested that overexpression of nELAVL-WT or their phosphodeficient mutant, but not the phosphomimetic mutant, led to impairment of orientation matching. Similar to *Cdkl5* KO mice, a mild effect was observed at spatial frequency 0.04cpd and no significant difference was observed at spatial frequency 0.02cpd (Figure S9C-D & F-G). For the more complex natural scene stimuli, C/I response correlation was decreased in nELAVL-WT, SE and SA injected mice when compared to Ctrl (Figure 6P), suggesting that binocular matching of complex stimuli is more sensitive to the expression level and phosphorylation status of nELAVLs. At the population level, decoding accuracy was decreased across viewing conditions in WT, SE and SA injected mice at spatial frequency 0.08cpd and natural scene when compared to Ctrl (Figure S9H & I). In sum, expression of nELAVL proteins with different phosphorylation status in V1 are sufficient to replicate the cortical visual deficits observed in *Cdkl5* KO mice.

## Discussion

Sensory processing deficits are a prevailing feature in numerous syndromic developmental disorders. Within the spectrum of developmental disorders, individuals with CDD are notable for their high rate of cortical visual impairment. In this study, utilizing a CDD mouse model, we delineated visual processing deficits of orientation tuning, interocular and monocular-binocular matching, and information encoding in the bV1. Our findings indicate that fundamental features of cortical visual processing, both at the single-neuron and network levels, is compromised in CDD mice, potentially impacting higher-order processing stages encompassing recognition, identification, and memory for visual patterns^67^. The binocular orientation mismatch observed in CDD mice likely diminishes their ability to differentiate similar stimuli and hinders the accurate representation of visual space.

At eye-opening in mice at P13-14, bV1 is predominantly composed of contra responsive cells which are already orientation selective^34^, while ipsi and contra+ipsi responsive cells emerge after eye-opening. The contra+ipsi responsive population later becomes orientation matched by experience-dependent plasticity during the ocular dominance critical period^61^. Our work reveals that in adult *Cdkl5* KO mice, both orientation selectivity and contra-ipsi orientation matching compromised, with more severe deficits in ipsi driven responses. L2/3 excitatory neurons in bV1 have been shown to undergo structural synaptic remodelling during the critical period dependent on experience-driven changes in gene expression^68^. We found alterations in L2/3 synaptic density and morphology, as well as the transcriptomic profile of *Cdkl5* KO mouse V1, with gene expression changes most prominent in L2/3 (Figure 2F-H). While the overall cell composition of V1 did not change, mRNA for the canonical immediate early gene *Fos*, a marker for neuronal activation and a direct binding target of nELAVLs, was identified as a hub gene for downregulated activity-dependent DEGs in *Cdkl5* KO mice (Figure 2B). Together, our data suggests that L2/3-mediated circuit impairment in *Cdkl5* KO mice is caused by dysregulated activity-dependent gene expression through the loss of CDKL5-dependent phosphorylation of nELAVLs. The deficits in contra-ipsi matching towards the contra eye further suggest that CDKL5-dependent phosphorylation of nELAVLs is most crucial during the developmental ocular dominance critical period. Given that cortical visual impairment is a well-reported clinical feature in CDD patients and has been suggested to be employed as a functional biomarker of CDD^6,26^, it will be of clinical interest to assess whether CDD patients also experience deficits in interocular integration.

Despite the identification of several physiological substrates, it remained unclear how CDKL5 dysfunction leads to cortical visual impairment at the molecular, cellular and neural circuit levels. Here we identified nELAVLs as direct substrates of CDKL5 and demonstrated that CDKL5-mediated phosphorylation can affect the propensity of nELAVLs to undergo phase separation. Our FRAP assay revealed a decreased exchange rate within ELAVL3 condensates in the absence of CDKL5-mediated phosphorylation, suggesting a more compact internal structure and less dynamic property of the condensates. CDKL5-dependent phosphorylation introduces negatively charged phosphate groups into the RPSS*A motif on nELAVLs, modulating charge-patterning critical for phase separation. This charge redistribution disrupts the balance of attractive (e.g., cation-π, π-π) and repulsive forces within nELAVLs, akin to phosphorylation-driven phase separation regulation in other RBPs including FUS and TDP-43 are known to undergo phase separation^69^. The negative charge introduced by phosphorylation may also disrupt hydrogen bonding with adjacent arginine/lysine residues^70^, altering RNA-binding domain conformation. Additionally, as activity-dependent DEGs, *Fos* and *homer1* are direct nELAVLs binding targets^33^, and our data demonstrated lower mRNA level of these DEGs, low RNA:RBP ratios may also modulate condensate size^71^. Given that impaired nELAVL phosphorylation was also observed in the frontal cortex and motor cortex of Cdkl5 KO mice, it will be of great interest to examine the roles of nELAVLs dysregulation in other clinically relevant phenotypes of CDKL5 deficiency, including motor and social deficits.

Our data highlight a CDKL5-mediated phosphorylation switch of nELAVLs that regulates the crosstalk between the nELAVL condensates and other biomolecular condensates. Stress granules and P-bodies are RNP granules that regulate mRNA processing^52–55^. The mRNAs sequestered within stress granules or P-bodies can be translationally repressed and these mRNAs may either re-enter the translation process after granule disassembly^53,54,72,73^ or undergo RNA decay^74–77^. In CDD models as well as in scenarios where nELAVLs are non-phosphorylated, we observed reduced levels of nELAVLs-binding mRNA concomitant with an augmentation in the volume of nELAVLs, stress granules, and P-bodies puncta. These results suggest that the loss of CDKL5 function in CDD patients leads to a downregulation of nELAVL phosphorylation, which in turn results in abnormal phase separation of nELAVLs. Such aberrant nELAVL phase separation then contributes to dysregulated stress granules and/or P-bodies, which sequester and prevent the activity-dependent translation of key mRNAs essential for synaptic plasticity, ultimately contributing to impaired refinement of neural circuits during the pathological, developmental progression of CDD (Figure S10). To elucidate how aberrant RNP condensates—including P-bodies and stress granules—coordinately dysregulate mRNA expression and translation in CDKL5 deficiency will be our future direction.

Altered dynamics of stress granules and P-bodies has been suggested to contribute to the pathology of several neurodegenerative conditions including amyotrophic lateral sclerosis, frontotemporal dementia, Alzheimer’s disease and Parkinson’s disease^78–80^. Genetic and functional studies also suggest that nELAVLs, stress granules, and P-bodies are strongly implicated in neurodevelopmental disorders. ELAVL3 and ELAVL4 null mice display deficits in learning, motor function, and neuronal maturation^81–83^. Differential gene expression analysis in ELAVL2 knockdown primary human neurons also reveals autism related alterations^84^. ELAVL3 has also been identified as an autism-associated gene in multiple large-scale genetic studies^85–87^, and both ELAVL2 and ELAVL3 are listed as autism candidate genes by the Simons Foundation Autism Research Initiative. Similarly, pathogenic variants of genes encoding for stress granules and P-bodies core proteins such as G3BP1, UBAP2L and TNRC6B have been identified in patients with neurodevelopmental disorders^88,89^. Given the pivotal roles of nELAVLs in a wide range of neurological disorders, it is reasonable to speculate that the dysregulation of CDKL5-mediated phosphorylation of nELAVLs, and the subsequent formation of aberrant nELAVL condensates, may also contribute to the etiology of other diseases.

In summary, our findings shed light on the physiological significance of CDKL5-mediated nELAVLs phase separation, as well as their interplay with other biomolecular condensates, in the context of activity-dependent visual circuit refinement. These findings deepen our understanding of the link between CDKL5 loss of function and cortical visual impairment in CDD, and potentially other pathology. Crucially, our discovery also opens up possibilities for exploring how phosphorylation governs the crosstalk among various biomolecular condensates which have broad and profound implications for normal brain function as well as neurodevelopmental and neurodegenerative disorders.

## Acknowledgments

We thank Gavin Siu, Josie Lai, Joaquim Vong and Cara Kwong for their technical assistance. We also thank Core Laboratories, School of Biomedical Sciences, the Chinese University of Hong Kong as well as Biosciences Central Research Facility, the Hong Kong University of Science and Technology for their technical support. We thank Hei Man Kim Chow for sharing HEK 293T and HT22 cells. We thank Hovy Wong and Kwok-On Lai for critical comments on the manuscript.

## Author contributions

S.Y. performed surgeries, two-photon imaging, phosphoproteomic and eMAP experiments. Y.Z. performed RNA extraction, real-time PCR, FISH, bulk and snRNA-seq data analysis. Zhongyu Z. performed cell line, primary neuron and iPSC experiments, STED imaging and quantification. H.Y. processed and analyzed the two-photon imaging data. Zhongjie Z. carried out population analysis for two-photon imaging data. K.J., K.T., G.H., J.Z., Marco C. and L.T. contributed to two-photon imaging data analysis. Maggie C. and Y.C. performed Western blot, with inputs from H.H.. Y.D., S.L. and A.F. contributed to snRNA-seq experiments. C.H., H.T. and Zhongyu Z. performed FRAP experiments. H.L. and H.K. provided expertise on the FISH experiment. J.L., A.K., S.F., Q.C. and S.T. contributed to snRNA-seq analysis. T.K. provided expertise on eMAP experiments. A.C. and K.L. provided expertise on cellular condensates experiments. C.G. and B.N. computed the phase separation propensity of nELAVLS. Y.W. and J.Z. performed the in vitro phase separation assay. J.I. conceived and supervised all aspects of the experiments. S.Y., Y.Z., Zhongyu. Z., H.Y. and J.I. wrote the paper. All authors edited the manuscript.

## Methods

### Key Resources Table

| Antibodies | Source | Catlog # |
| --- | --- | --- |
| Rabbit anti-ELAVL2 | Proteintech | 14008-1-AP |
| Rabbit anti-nELAVL pS | Biomatik | custom-made |
| Rat anti-EB2 | Abcam | ab45767 |
| Rabbit anti-EB2 pS222 | Covalab | 00117741 |
| Rabbit anti-MAP1S | Sigma | HPA050934 |
| Rabbit anti-MAP1S pS812 | Covalab | 00118044 |
| Mouse monoclonal anti-GAPDH | Invitrogen | AM4300 |
| Mouse monoclonal anti-CDKL5 | Santa Cruz | sc-376314 |
| Rabbit monoclonal anti-c-fos | Synaptic Systems | 226008 |
| Chicken polyclonal anti-GFP | Invitrogen | A10262 |
| Rabbit polyclonal anti-ELAVL3 (HuC) | Proteintech | 55047-1-AP |
| Mouse monoclonal anti-ELAVL4 (HuD) | Santa Cruz | sc-48421/sc-28299 |
| Mouse monoclonal anti-GW182 | Santa Cruz | sc-56314 |
| Mouse monoclonal anti-TIA1 | Santa Cruz | sc-166247 |
| Goat anti-chicken, Alexa 488 | Invitrogen | A32931 |
| Goat anti-rabbit, Alex 555 | Invitrogen | A32732 |
| Goat anti-mouse, Alex 647 | Invitrogen | A32728 |
| Goat anti-rabbit, Alex 647 | Invitrogen | A32733 |
| Abberior STAR ORANGE | Abberior | STORANGE |
| Abberior STAR RED | Abberior | STRED |

| **Bacterial and virus strains** | **Source** | **Catlog #** |
| --- | --- | --- |
| rAAV-hSyn-GCaMp6f-WPRE-hGH polyA | Brain VTA | PT-0147 |
| AAV9-hSyn-nElavl-P2A-DsRed-WPRE-BGH-polyA | Vigene | / |
| AAV9-hSyn-nElavl(SE)-P2A-DsRed-WPRE-BGH-polyA | Vigene | / |
| AAV9-hSyn-nElavl(SA)-P2A-DsRed-WPRE-BGH-polyA | Vigene | / |
| pCAG-FLEX-EGFP-WPRE | Addgene | 51502 |
| pENN.AAV.CamKII 0.4.Cre.SV40 | Addgene | 105558 |

| **Experimental models: Organisms/strains** | **Source** | **Catlog #** |
| --- | --- | --- |
| Mouse: *Cdkl5* KO | The Jackson Laboratory | Strain #:021967;<br>RRID:IMSR_JAX:021967 |
| Mouse: C57BL/6J | Laboratory Animal Services Centre, The Chinese University of Hong Kong | / |

| **Software and algorithms** | **Source** | **Catlog #** |
| --- | --- | --- |
| MATLAB | Mathworks | https://www.mathworks.com/products/matlab.html |
| GraphPad Prism 9 | GraphPad Software Inc | https://www.graphpad.com/scientific-software/prism/ |
| Fiji | Open source | https://imagej.net/software/fiji/ |
| Suite2p | Open source | https://suite2p.readthedocs.io/en/latest/ |
| Imaris | Oxford Instruments | https://imaris.oxinst.com/ |
| Python | Anaconda3 | https://www.anaconda.com |
| R | R version 4.1.2 | https://www.r-project.org/ |

### Resource availability

#### Leading contact

Further information and requests for resources and reagents should be directed to and will be fulfilled by the lead contact.

#### Materials availability

This study did not generate new unique reagents.

#### Data and code availability

Any additional information to reanalyze the data reported in this paper is available upon request. All raw and processed snRNA-seq datasets reported in this study will be available before publication. Analysis code generated during this study are available upon reasonable request.

### Methods details

#### Mouse line

*Cdkl5* KO mouse line^5^ (B6.129(FVB)-*Cdkl5 tm1.1Joez /J,* strain no: 02196, Jackson Laboratory) was imported from the Jackson Laboratory. A colony was maintained at the Laboratory Animal Services Centre of the Chinese University of Hong Kong and housed in the Animal Holding Core of the School of Biomedical Sciences, the Chinese University of Hong Kong. *Cdkl5* heterozygous females (+/-) were used as breeders with C57BL/6 wildtype males to produce WT and *Cdkl5* KO littermates. Our study mainly focused on investigating hemizygous male mice as our model to avoid potential confounds induced by mosaic expression of CDKL5 in heterozygous females resulting from random X chromosome inactivation. C57BL/6 mice were used for nELAVLs, nELAVLs-SE, nELAVLs-SA virus injection for two-photon calcium imaging. Mice were group-housed (no more than five mice per cage) with a standard light/dark cycle of 12/12 hours with access to food and water *ad libitum*. All experimental procedures were approved by the Animal Experimentation Ethics Committee of The Chinese University of Hong Kong.

#### Genotyping

A PCR-based method was used to detect deletions in *Cdkl5* exon 6 for genotyping (Phire Tissue Direct PCR Master Mix, Thermo Fisher Scientific). Dilution Buffer and DNA Release from the genotyping kit were mixed for genomic DNA extraction for 5 minutes at 98 °C, 20 μl Dilution Buffer and 1 μl DNA Release for each sample. 1 μl of genomic DNA extracted from the ear tissue was used for PCR. Primers for genotyping are forward: 5’-CCACCCTCTCAGTAAGGCAGCAG-3’ and reverse: 5’-GTCCTTTTGCCACTCAATTCCATCC-3’^5^. PCR reactions contained 0.5 μl of each primer, 10 μl 2X Phire Tissue Direct Master Mix and 8 μl nuclease-free water. PCR was performed using the following program: 1) 95°C for 3 min, 2) 98°C for 20 s, 3) 60°C for 15 s, 4) 72°C for 1 min, 5) 72°C for 2 min, 6) 4°C for 10 minand finish. Repeat steps 2-4 for 27 times. The WT allele showed a 653 base pair product, and the *Cdkl5* KO allele showed a 305 base pair product.

##### Phosphoproteomics

V1 of 4 pairs WT and of *Cdkl5* KO littermates (8-week old) were processed for TMT labeling by Shanghai Applied Protein Technology Co., Ltd. For protein extraction and digestion, tissues were separately lysed in SDT buffer (4% SDS, 100 mM Tris–HCl, and 1 mM DTT [pH 7.6]) and quantified using the BCA Protein Assay Kit. Digested peptides of each sample were sonicated by C18 Cartridges and concentrated by vacuum centrifugation, followed by reconstituting in 40 µl of 0.1% (v/v) formic acid. 280 nm UV light spectral density was used to estimate the peptide content. The peptide mixture of each sample (100 μg) was labeled with TMT (Thermo Fisher Scientific) reagent. Phosphorylated peptides were subjected to TiO2 bead-based enrichment (90% enrichment efficiency) followed by LC-MS/MS analysis in Thermo Scientific Q Exactive HF mass spectrometer (Thermo Scientific) that was coupled to Easy nLC (Proxeon Biosystems, now Thermo Fisher Scientific) for 120 min. The peptides were loaded onto a reverse phase trap column (Thermo Scientific Acclaim PepMap100, 100 μm*2 cm, nanoViper C18) connected to the C18-reversed phase analytical column (Thermo Scientific Easy Column, 10 cm long, 75 μm inner diameter, 3μm resin) in buffer A (0.1% Formic acid) and separated with a linear gradient of buffer B (84% acetonitrile and 0.1% Formic acid) at a flow rate of 300 nl/min controlled by IntelliFlow technology. The mass spectrometer was operated in positive ion mode. MS data was acquired using a data-dependent top10 method dynamically choosing the most abundant precursor ions from the survey scan (300–1800 m/z) for HCD fragmentation. Automatic gain control (AGC) target was set to 3e6, and maximum inject time to 10 ms. Dynamic exclusion duration was 40.0 s. Survey scans were acquired at a resolution of 70,000 at m/z 200 and resolution for HCD spectra was set to 17,500 at m/z 200, and isolation width was 2 m/z. Normalized collision energy was 30 eV and the underfill ratio, which specifies the minimum percentage of the target value likely to be reached at maximum fill time, was defined as 0.1%. The instrument was run with peptide recognition mode enabled. For TiO2 enrichment, samples were reconstituted in pre-cooled IAP Buffer (1.4ml) and followed by adding TiO2 beads, shaken for 40 minutes and centrifuged to remove the supernatants. Beads were put into the tips and washed with the washing buffer 3 to 6 times. Phosphopeptides were eluted and concentrated using an elution buffer, followed by dissolving in formic acid (20 μl 0.1%) for MS analysis. For LC-MS/MS analysis, a reverse phase trap column connected to a C18-reversed phase analytical column was used to load the peptides. A positive ion mode was used for the mass spectrometer, and MS data were acquired using a data-dependent top10 method. Peptides and proteins were mapped by Proteome Discoverer 2.4 (Thermo Electron). The parameters set as follow: Enzyme: Trypsin; Max Missed Cleavages: 2; Variable modifications: Oxidation (M), Phospho (ST) / Phospho (Y); Peptide Mass Tolerance: ± 20 ppm; Fragment Mass Tolerance: 0.1Da; Database pattern: Decoy; Peptide FDR: ≤0.01; Protein FDR: ≤0.01; Experimental Bias: Normalizes all peptide ratios by the median protein ratio. The median protein ratio should be 1 after the normalization. Downregulated genes meeting the criterion of p < 0.05 were selected for GO analysis with all genes we identified in phosphoproteomics as background genes. Top 10 GO terms were mapped, and sequences were annotated using gProfiler.

#### Western blotting

Samples were collected and homogenized in RIPA lysis buffer supplemented with protease and phosphatase inhibitors, and were incubated for 30 min at 4°C. The resultant lysates were cleared by centrifugation at 20,000g for 20 min at 4°C. Proteins were denatured by 1x sample buffer supplemented with 0.1M DTT and incubated at 95°C for 5-7 min. Samples were then resolved in 10% Tri-glycine SDS-PAGE gel. Proteins were then transferred onto a PVDF membrane at 100V for 2 hr on ice. The membranes were blocked by 5% non-fat milk in TBS and probed with primary antibody diluted in 2% non-fat milk in TBST at 4°C overnight: nELAVL (1:4000, Proteintech), pS-nELAVL (1:1000, Biomatik, rabbit polyclonal phosphospecific antibodies were raised against SYARPS(pS)ASIRDAN), EB2 (1:2000, Abcam), pS222-EB2 (1:2000, Covalab), MAP1S (1:1000, Sigma Aldrich), pS812-MAP1S (1:1000, Covalab), GAPDH (1:7500, Life Technologies), CDKL5 (1:500, Santa Cruz). Following incubation with 1:5000 HRP-conjugated secondary antibody (2% in non-fat milk in TBST) for 1 hr at room temperature, membranes were washed with 1x TBST for five times, and 10 min for each wash. Signal was detected using enhanced chemiluminescent substrate. The expression of targeted proteins was quantified by ImageJ software.

#### Immunohistochemistry

Mice were perfused with ice-cold phosphate-buffered saline (PBS) and 4% paraformaldehyde transcardially. Brains were collected and incubated in 4% paraformaldehyde at 4^°^C overnight.

Brain was sectioned by vibratome and then subjected to immunostaining. Brain sections were blocked for 1 hr in PBS with 2% horse serum and 0.4% Triton-100 followed by an overnight incubation at 4^◦^C with rabbit anti-c-fos primary antibody (1: 2500, Synaptic Systems). Brain sections were washed and incubated with a secondary antibody (1: 1000, Invitrogen) for 1 hr at RT. After PBS rinsing, slices were mounted on glass slides. Confocal microscopy (TCS SP8, Leica) was used for imaging.

#### RNA extraction and real-time PCR

Total RNA from mouse V1 was extracted using NucleoSpin RNA kit (Macherey–Nagel), following the manufacturer’s instructions. Reverse transcription of RNA into cDNA was conducted using a PrimeScript RT Reagent Kit (RR037A, Takara Bio). Quantitative real-time PCR was conducted with a Premix Ex Taq Kit (RR390A, Takara Bio) using ABI ViiA7 Real Time PCR System (Applied Biosystems). The mRNA expression was normalized to that of *Gapdh*.

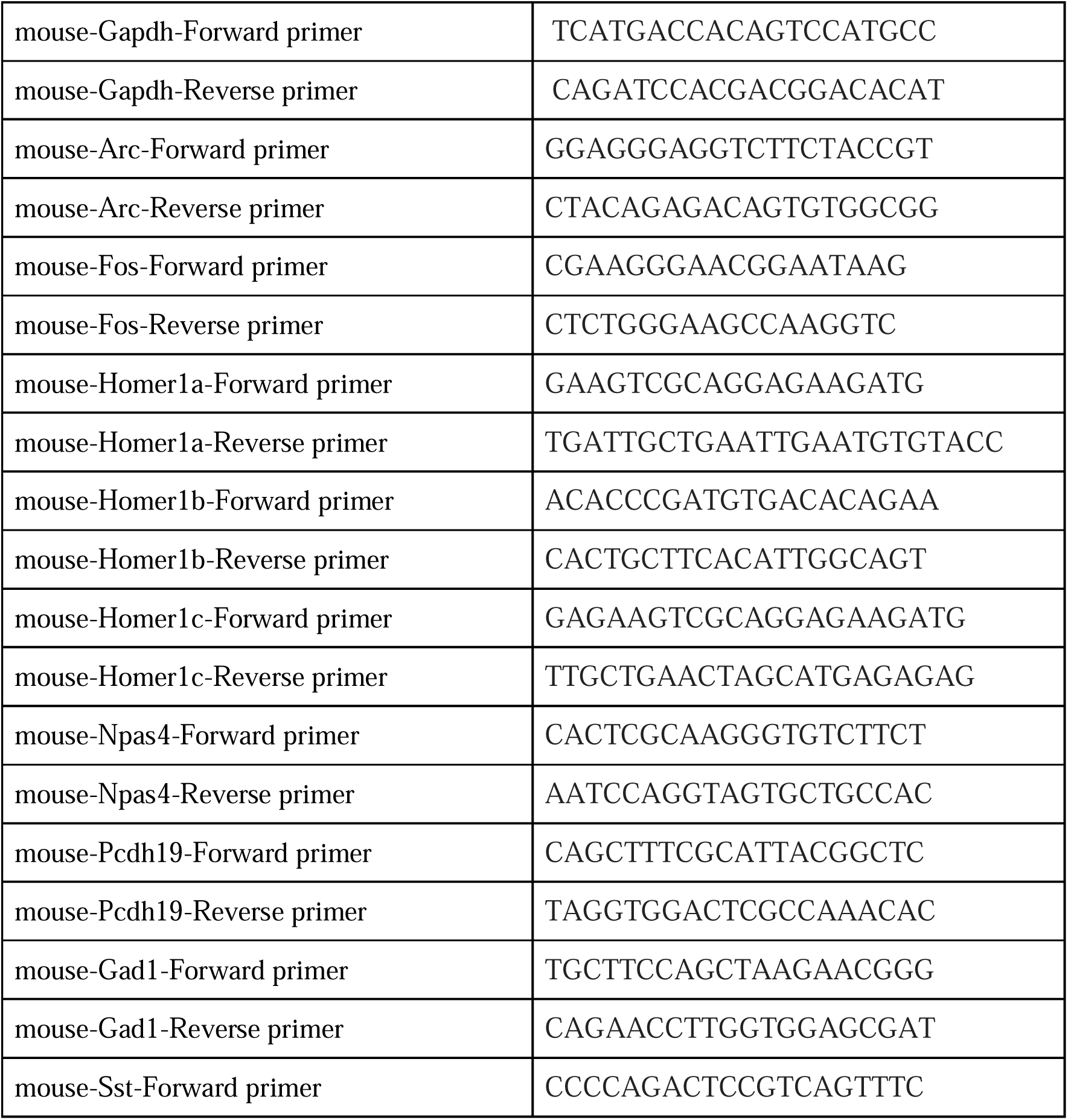
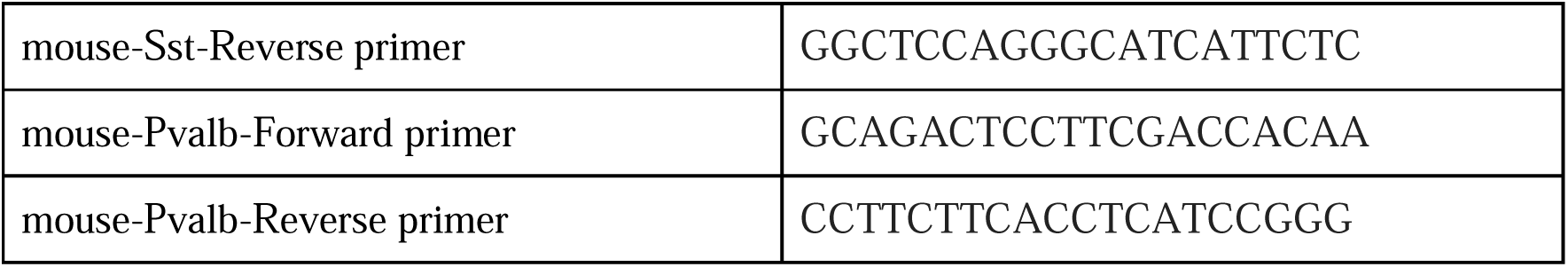

#### Bulk RNA sequencing and data analysis

RNA was extracted from V1 tissues of 6 pairs WT and *Cdkl5* KO mice (8-week-old) using NucleoSpin RNA column (Macherey–Nagel), and RNA integrity was assessed by Agilent 4200 TapeStation System. mRNA enrichment fragmentation and cDNA library construction were performed by Novogene Co. The transcriptomic profile for each sample was generated by Novogene Co utilizing the Illumina Novaseq 6000 platform, 150bp paired-end sequencing. For RNA sequencing, quality control of the datasets was firstly performed using FastQC^90^. Reads alignment was performed by HISAT2^91^ and counting was conducted by FeatureCounts^92^. Differential gene expression analysis was performed using R package DESeq2^93^. DEGs between WT and *Cdkl5* KO mice were selected using the following criteria: foldchange >C1.2 or <C−0.83 and *padj*C<C0.05. STRING was used to construct PPI network for DEGs. The PPI network with minimally required interaction score of high confidence (0.7) were visualized in Cytoscape (3.10.0). Maximal Clique Centrality (MCC) algorithm in CytoHubba was used to calculate and identify hub genes. DEGs with log2Foldchange were subjected to GSEA, which was performed by clusterProfiler^94^ that enabled GO term enrichment analysis.

#### Isolation of V1 nuclei for snRNA-seq

Nuclei from 6 pairs of frozen WT and *Cdkl5* KO mouse (8-week-old) V1 samples were isolated following previously established protocol^95^. WT and *Cdkl5* KO mice were euthanized as described above. Mouse V1 was dissected in ice-cold DPBS and immediately snap-frozen in liquid nitrogen. Frozen V1 tissues from two animals were pooled and transferred to a prechilled Dounce homogenizer containing ice-cold homogenization buffer. This buffer consisted of 250 mM sucrose, 25 mM KCl, 5 mM MgCl2, 20 mM tricine-KOH (pH 7.8), 1 mM dithiothreitol, 0.15 mM spermine, 0.5 mM spermidine, 5 μg/mL actinomycin, 0.16% Nonidet P-40, 0.04% bovine serum albumin, protease inhibitors, and 200 U/mL RNAsin. The tissues were gently homogenized on ice using a loose pestle until no visible solid fragments remained. The lysate was subsequently combined with OptiPrep (Sigma) at a ratio of 1:1 (v/v) and centrifuged at 10,000 × *g* for 20 min at 4 °C. The supernatant was removed and the nuclei were washed once with Dulbecco’s modified Eagle medium/F12 supplemented with 5% fetal bovine serum. The nuclei were resuspended in Dulbecco’s modified Eagle medium/F12 supplemented with 10% fetal bovine serum and 200 U/mL RNAsin. The resuspended nuclei were counted on a hemocytometer using Trypan blue labeling and diluted to 400 nuclei per microliter. Nuclei were recounted on an automated cell counter (RWD) before further processing.

#### snRNA-seq library preparation

snRNA-seq libraries of WT and *Cdkl5* KO mouse V1 samples were constructed by Chromium Single Cell 3′ Library Kit v3 (1000078; 10× Genomics). 40 μL suspension of diluted nuclei for each sample was loaded and processed according to manufacturer’s instructions. The libraries were subjected to 150bp paired-end sequencing on an Illumina NovaSeq 6000 system (Novogene Co), obtaining a minimum of 200 GB raw data for each library.

#### snRNA-seq analysis

The demultiplexed FASTQ files were mapped to the GRCm38 genome reference using Cell Ranger (version 7.1.0) with the default settings. The transcriptomic analysis was performed using a Python package called SCANPY^96^. Raw gene expression matrices from 6 snRNA-seq libraries (each library was derived from two animals) were combined for analysis. For quality control, the following criteria was used to exclude low-quality cells or outliers: n_genes > 6500 or < 700, percent_mito > 5%, and n counts > 40,000. Genes detected in less than 8 cells were removed^37^. Cells were normalized using *scanpy.pp.normalize_per_cell* with default setting, followed by log-transformation.

After quality control, highly variable genes (HVGs) were identified using *scanpy.pp.highly_variable_genes* with default setting. Harmony^36^ was employed for batch correction. The top 40 principal components (PCs) were used to compute a nearest-neighbor graph on the cells using *scanpy.pp.pca* function. Leiden algorithm^97^ was applied to do clustering with default setting and the graph was finally embedded in 2D via the Uniform Manifold Approximation and Projection (UMAP) algorithm^98^. Canonical marker genes which were reported in previous studies^37^ were used for annotation. Doublets were identified using Scrublet^35^. Clusters that expressed markers of more than two classes or contained more than 50% doublets were discarded.

To further identify types in each class, this combined gene expression matrix was divided into 3 matrices: glutamatergic neurons, GABAergic neurons and non-neuronal cells. The raw matrix of glutamatergic neurons was further processed for highly variable gene selection, batch correction, dimensionality reduction and clustering as described above and visualized by UMAP. Previously reported type-specific markers^37^ were used for type annotation of glutamatergic neurons.

Differential expression analysis was performed to assess the transcriptomic alterations between WT and *Cdkl5* KO mouse V1, and to identify markers in each subclass or type. Differential expression analysis in this study was achieved using *scanpy.tl.rank_genes_groups* function in the SCANPY package. Differentially expressed genes were defined as: adjusted P value < 0.05 and a log2 fold change > 0.25 or < −0.25.

Enrichment for activity-dependent genes in visual cortex^99^ was calculated using the R package GeneOverlap. The lists of DEGs for each cell type were tested, utilizing all genes existing in snRNA seq from this study as the background. *P* values from Fisher exact test were reported.

#### Fluorescent in situ hybridization (FISH)

WT and *Cdkl5* KO mice (8-week-old) were euthanized and perfused as described. The brains were removed and fixed in 4% PFA for 24hr at 4 °C, followed by sucrose gradient until dehydration. Mouse V1 tissues were dissected under microscope and embedded in optimal cutting temperature compound (OCT, Tissue-Tek), and frozen at -80°C. The V1 tissues were sectioned at 20 µm with a cryostat (Epredia Cryostar NX70 Cryostat). The sections were fixed with 4% PFA for 10 min, washed 3 times with PBS and dehydrated using 70% ethanol. The sections were permeabilized by 8% SDS in PBS for 15 min at room temperature. The sections were then labelled according to the manufacturer’s protocol^100^ (Molecular Instruments). DAPI was stained before imaging. Probes for *Fos* and *Arc* were commercially designed by the manufacturer (Molecular Instruments). Probe for *Homer1* and *Camk2a* were custom designed and the target sequences are listed below:

Homer1-1 GGCTATCAGCCCATTGGCCAAACTTTTGAGATGTTTTAGTAAATGTCATG

Homer1-2 GCCGAGCAGCTTCCTTAAATTCCTGAAACTTTTCTGCGAATTTTGAAAGA Homer1-3 GCTGAAGATAGGTTGCTCCCC

Homer1-4 GGTGTGATGGTGCTATTTATTATTGCCTTTGAGCCATCTAAACTG Homer1-5 AAGGGGTACTGGTCAGCTCC

Camk2a-1 CCACAGCGTGAGAAAGAGCAGCAT Camk2a-2 TTCCCCAGGGCCTCTGGTTCAA Camk2a-3 ATCTGCCATTTGCCGTCCCTGC

FISH assay on rat primary hippocampal neurons and iNeurons were performed with ViewRNA Fos probe following manufacturer’s protocol (Rat: Catalog number: VX-06, Assay ID: VC1-11292-VCP; Human: Catalog number: VX-01, Assay ID: VA1-11504-VC; Thermo Fisher Scientific).

Leica SP8 confocal microscope with a 40×, NA 0.85Cobjective (SBS core lab, CUHK) was used for imaging. V1 cortical areas were imaged with optical sectioning of 0.5 μm and 15 optical sections. Channels were imaged sequentially to avoid any optical crosstalk. The number and volume of puncta within cells were quantified using the Imaris reconstruction software (v.9.0, Bitplane).

#### Epitope-preserving Magnified Analysis of Proteome (eMAP), imaging and analysis

We performed eMAP as previously described^101,102^. Mice under anesthesia were perfused with PBS followed by 4% paraformaldehyde. Brain tissues were collected and post-fixed in 4% paraformaldehyde overnight at 4C. Brain tissues were then embedded in MAP solution (30% acrylamide, 10% sodium acrylate, 0.1% bisacrylamide, and 0.03% VA-044 in PBS) under hypoxic conditions. The embedded samples were then immersed in hydration solution (0.02% sodium azide in PBS) and sectioned at 60 μm using a Leica VT1000S vibratome. Lipid was removed by incubation in clearing solution (6% SDS, 0.1 M phosphate buffer, 50 mM sodium sulfite, 0.02% sodium azide in deionized water [pH 7.4]) for 4 hours at 37C. Samples were then washed with PBST (0.1% Triton X-100 and 0.02% sodium azide in PBS) and stained with primary anti-GFP (A10262, Invitrogen) antibody diluted in PBST overnight at 37C. After washing with PBST, samples were incubated with secondary antibodies diluted in PBST overnight at 37C and then washed again with PBST. The samples were expanded and mounted in 0.01× PBS immediately before imaging. Approximately 3× total linear expansion was achieved, and imaging was performed under a Leica SP8 confocal microscope using a 20X lens with a 0.75 numerical aperture or 63X water immersion lens with a 1.2 numerical aperture. For dendritic spine morphology analysis, FIJI (version 2.14) was used to measure head diameter (D_head_), neck diameter (D_neck_), and the length (L_length_) of dendritic spines according to dendritic spine classification criteria previously described^103^.

#### Site-directed mutagenesis

Wildtype sequences of ELAVL 2 to 4 were directly cut out by corresponding restriction enzymes and ligated to in-frame EGFP-C vectors. Then, by using them as templates, SE mutants of Elavls 2 to 4 were built using traditional Primer Extension Mutagenesis with two-stage nested PCR^104^. While SA mutants were built using one-stage PCR and Dpn1 digestion method^105^.

Addgene plasmids carrying wild-type sequences of ELAVL 2 to 4:

ELAVL 2: pENTRY4_HuB (65752);

ELAVL 3: pENTRY4_HuC (65753);

ELAVL 4: pENTRY4_HuD (65754).

Primers for SE mutants:

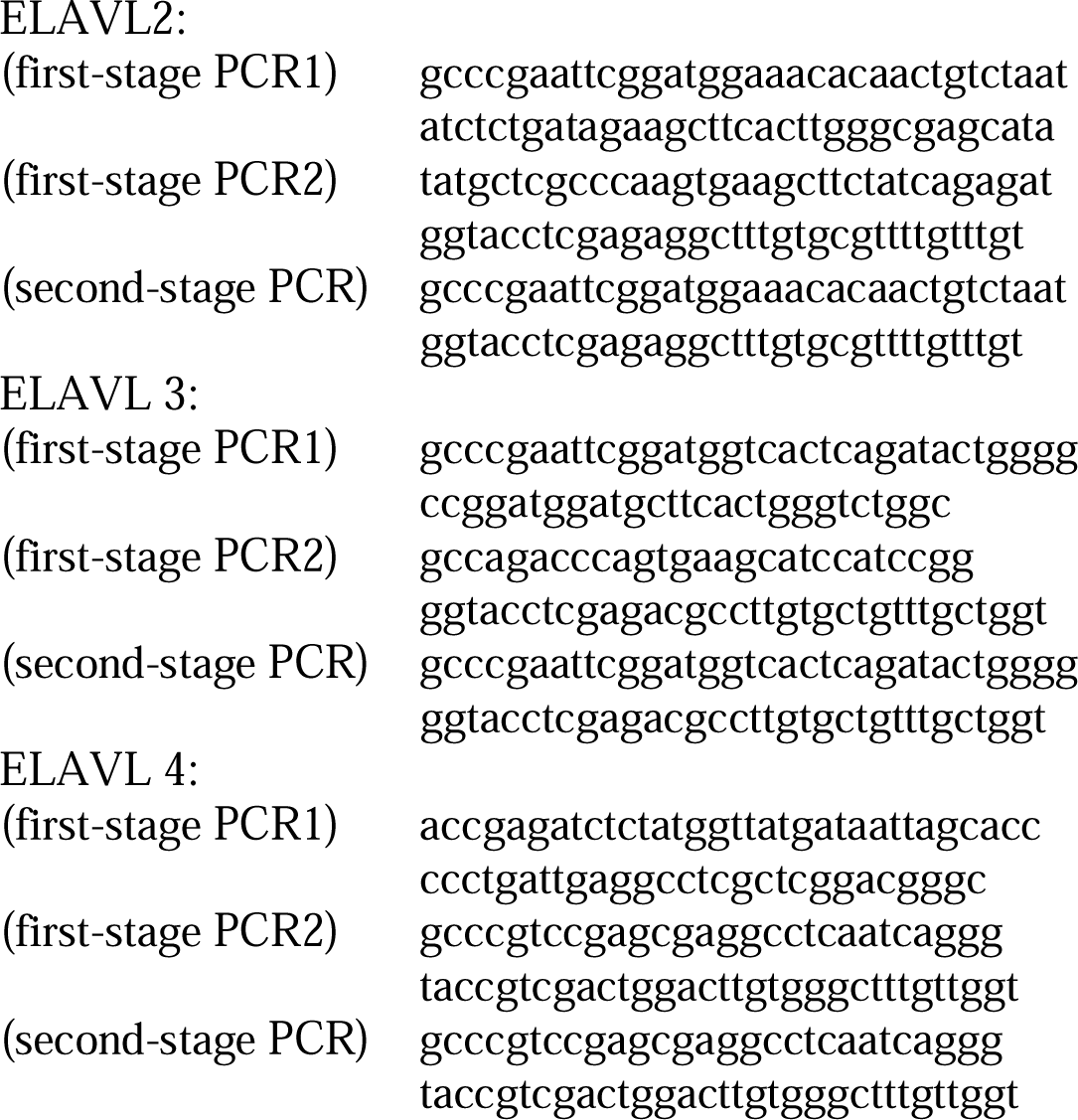

Primers for SA mutants:

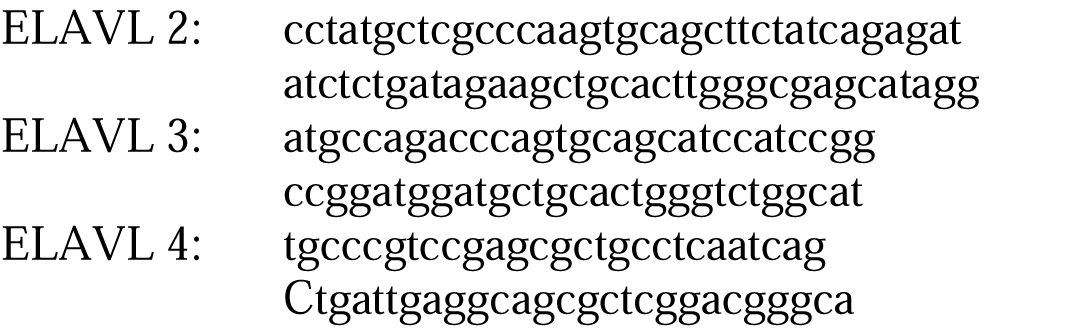

#### LLPhyScore algorithm

The algorithm computed a composite LLPS_score for each entry by integrating eight distinct biophysical features derived from PDB statistical potentials, including solvent accessibility and pi-pi interaction frequencies. We subsequently constructed a global rank-ordered distribution curve by sorting all sequences by their predicted scores. This framework allowed for a quantitative, comparative assessment of the ELAVL family’s biophysical driving forces relative to both established high-confidence scaffolds and the general baseline of the proteome.

#### Cell line culture, transfection and drug treatment

The primary hippocampal neurons were prepared from embryonic day 18.5 rat embryos. Hippocampal neurons (1 × 10^5^) were plated on 10 μg/ml poly-D-lysine (Sigma Aldrich) coated 18 mm coverslips in 12-well plates, fed with Neurobasal medium (21103-049, Gibco) supplemented with 2% B27 (17504-044, Gibco) and 1% GlutaMAX (35050-061, Gibco). We performed calcium phosphate transfection on div 11 and fixed the neuron for immunostaining following previously described protocol on div 17^106^. The HEK 293T and HT22 cells were grown in Dulbecco’s Modified Eagles Medium (DMEM) supplemented with 10% of fetal bovine serum and 1% of penicillin/streptomycin solution up to 40 passages. We used the general PEI transfection protocol by adding 0.4 μg DNA/coverslip, transfect overnight and perform immunostaining. For Actiomycin D treatment, we replenish the iNeurons with neurobasal medium containing 2 μg/mL Actinomycin D at different time point, and fixed them together for FISH assay.

#### iPSC culture

Human iPSC lines OR00005 (p.R550* mutant allele clone) and OR00006 (WT allele clone) were purchased from Coriell Institute. The two lines were reprogrammed from a heterozygous CDD female patient carrying a pathogenic variant 1648C>T (p.R550*) of CDKL5 in one allele as well as a WT allele. To generate adenine base editor (ABE)-corrected iPSCs, we electroporated 5 μg ABEmax plasmid with a guide RNA to correct the mutation using Human Stem Cell Nucleofector 1 (Lonza, VPH-5012) according to the manufacturer’s protocols. ABE-corrected iPSC clones were selected with puromycin and confirmed through sanger sequencing. iPSCs were cultured in Essential 8 (E8) Medium (Thermo Fisher Scientific, A1517001) on 6-well cell culture dishes (SPL, 30006) coated with Geltrex (Thermo Fisher Scientific, A1414402) diluted 1:100 in DMEM/F-12 (Thermo Fisher Scientific, 11330032). E8 medium was daily replenished. When iPSC colonies demonstrated mature morphology, the iPSCs were either clump passaged with EDTA (Thermo Fisher Scientific, 25300054) for routine maintenance or differentiation into cortical neurons. For clump passaging with EDTA, iPSCs were washed with Dulbecco’s PBS (DPBS; Millipore Sigma, D8537) and then incubated with EDTA for 5–7 min at 37C; the EDTA solution was then aspirated and replaced with E8; iPSC colonies were then gently detached and resuspended gently and centrifuged at 300 g for 3 min. Then the supernatant was discarded, and cell pellets were resuspended with E8 + Y-27632 (Sigma Aldrich, Y0503) and passaged at 1:10–1:40 dilutions in E8 + Y-27632, with Y-27632 removed the next day.

#### Two-dimensional neuron differentiation

For differentiation of iPSCs into cortical neurons, published protocols were followed with minor modifications^107^. iPSCs were plated on 6-well Geltrex-coated plates and suspended lentivirus with rtTA and Ngn2-GFP-puro vectors 2 hours later in E8 medium containing 10 μM Y-27632 and infected the iPSCs for 24 hours. After the colonies demonstrated mature morphology (Day 0), the culture medium was replaced with N2 medium (DMEM/F-12, 1 × N2 and 1 × NEAA; Thermo Fisher Scientific) containing 2 μg/mL doxycycline (Sigma Aldrich). On day 1, the culture medium was replaced with N2 medium containing 2 μg/mL doxycycline and 1 μg/mL puromycin (Thermo Fisher Scientific, A1113803). On day 3, the induced cells were dissociated with Accutase (Thermo Fisher Scientific, A1110501) and plated them onto 6-well PDL/laminin coated plates at 1.0 × 10^6^ cells per well or 24-well plates with coverslips at 5 × 10^4^ cells per coverslip in B27 medium (Neurobasal medium, 1 × B27, 1 × GlutaMAX and 0.2 μg/ml laminin, Thermo Fisher Scientific) containing 2 μg/mL doxycycline, 1 μg/mL puromycin. For further experiments, from day 7 until day 28, half of the medium was removed every 4 days and was replaced with B27 medium containing 0.5 μg/mL doxycycline, 10 μg/mL BDNF and 10 μg/mL GDNF.

#### Immunocytochemistry

Cells grown on coverslips or chamber slides were fixed with 4% paraformaldehyde (Sigma, F8775) for 15 min at room temperature. Then the cells were permeabilized with 0.1% Triton X-100 in DPBS for 5 min at RT and blocked with 1% BSA in DPBS for 20 min. Primary antibodies were incubated with the cells overnight at 4℃. The following primary antibodies were used: anti-Elavl2 (Proteintech, 14008-1), anti-Elavl3 (Proteintech, 55047-1), anti-Elavl4 (Santa Cruz, sc-48421 and sc-28299), anti-GFP (Invitrogen, A10262), anti-TIA1 (Santa Cruz, sc-166247), anti-GW182 (Santa Cruz, sc-56314). After washing three times in DPBS, the cells were incubated with secondary antibodies and DAPI for 1 hr at room temperature. The following secondary antibodies were used: Alexa Fluor 488 (Invitrogen, A32931), Alexa Fluor 555 (Invitrogen, A32732), Alexa Fluor 568 (Invitrogen, A21124), Alexa Fluor 594 (Invitrogen, A32742), Alexa Fluor 647 (Invitrogen, A32728, A32733), STAR Orange (Abberior, STORANGE-1001, STORANGE-1002) and STAR Red (Abberior, STRED-1001, STRED-1002). After washing for 3 times, the samples were mounted with anti-fade mounting medium.

Confocal images were acquired using Leica SP8 microscope with 63x objective, Z-series images were collected using z-serial scanning mode (20 optical sections at 0.3 μm steps). STED images were acquired using abberior STED microscopy system at 50 nm resolution, z-series images were collected using z-serial scanning mode (100 optical sections at 75nm per step).

#### Quantification of puncta number, volume and fraction distribution

The number and volume of puncta within cells were quantified using the Imaris reconstruction software (v.9.0, Bitplane). 3D reconstruction was performed on high-resolution confocal/STED images. The border of cells and nucleus was determined by surface module in the software, rendering with 0.1-μm smoothing, and disconnected processes were merged with the cell body to create a single surface. Then the spots module was used to measure the number and volume of the fluorescence puncta. The puncta within cells were limited by distance between puncta and cell surface/Nucleus surface. The fraction distribution analysis was done by fraction fining puncta within each cell by their volume, following by non-linear regression.

#### Protein expression and purification

Human *ELAVL3* (GenBank: NM_001420.4) gene was cloned into a modified pET32a vector with an N-terminal Trx-His_6_-tag. Full-length ELAVL3 was expressed in *Escherichia coli* BL21 (DE3) host cells at 37°C for 4 hr induced by 0.2 mM isopropyl-β-D-thiogalactoside (IPTG). Cells were harvested and resuspended in buffer containing 50 mM Tris, pH 8.0, 1 M NaCl, 5% Glycerol, 2 mM β-ME and 10 mM imidazole, and then lysed by high pressure homogenizer at 4°C. After centrifugation at 18,000 rpm for 20 min, the supernatants were loaded onto the Ni^2+^-NTA agarose affinity column and eluted with the buffer containing 50 mM Tris, pH 8.0, 1 M NaCl and 500 mM imidazole, and then purified by a size-exclusion chromatography in the buffer containing 50 mM Tris, pH 8.0, 1 M NaCl, 5% Glycerol, 1mM EDTA and 1 mM DTT.

#### Protein labeling with fluorophore

Purified ELAVL3 proteins (∼5 mg/mL) were dissolved in the buffer containing 300 mM NaCl, 100 mM NaHCO_3_, pH 8.3, 4 mM β-ME. Cy3 NHS ester (AAT Bioquest) was dissolved in DMSO and incubated with the corresponding protein (molar ratio ∼1:1) at room temperate for 1 hr. The labeling reaction was quenched by the buffer of 200 mM Tris, pH 8.2, and the labeled protein was purified with a HiTrap desalting column with the buffer containing 50 mM Tris, pH 8.0, 1 M NaCl, 5% Glycerol, 1mM EDTA and 1 mM DTT. Fluorescence labeling efficiency was determined by Nanodrop 2000 (Thermo Fisher).

#### In vitro phase separation assay

The stock ELAVL3 protein was centrifuged at 16,873 g for 10 min at 4°C to remove any precipitations and diluted to designed concentrations and buffer conditions. Subsequently, all the samples were placed on ice before phase separation assay. For microscope-based assays, each sample was injected into a home-made chamber for fluorescence imaging (Leica TCS SP8).

#### Fluorescence recovery after photo-bleaching assay

For FRAP of ELAVL3 SA and SE overexpressed HEK 293T cells, a circular ROI was bleached by 488 nm laser beam at 37C. The photobleaching was acquired with 50 cycles of 488 nm laser at 10% power and imaging every 0.5 s for 5 min after bleaching. The fluorescence intensity of the bleached area was normalized to the intensity of the same area before bleaching. The recovery curves were corrected with the region without photobleaching in the same frames.

For Cy3-labeled protein droplets, ROI was bleached by a 561 nm laser beam at room temperature. A dwell time of 1.2 s and 100% of laser power was used during bleaching. Fluorescence recovery was monitored for 300 s at 5 s intervals. For each experiment, the fluorescence intensity of a neighboring droplet with similar size to the bleached one was also recorded for intensity correction. Background intensity was subtracted before data analysis. The pre-bleaching ROI intensity was normalized to 100%.

#### Co-Immunoprecipitation

Protein extraction was performed in HEK 293T cells using a NP40 buffer (50 mM Tris-HCl buffer (pH 7.4), 150 mM NaCl, 2 mM MgCl, 1 mM EDTA, 0.5% (v/v) NP40, 1 mM DTT and complete EDTA-free protease inhibitor cocktail). Pre-clearing of protein extract was performed by 30 minutes R/T incubation with Protein A magnetic beads (Life Technologies).

Pre-cleared extract was incubated with 2 μg of GW182 antibody (sc-56314; Santa Cruz Biotechnology) for O/N at 4 ° C, then performed immunoprecipitation with Protein A magnetic beads by 30 minutes R/T incubation. Samples were washed 5 times using NP40 buffer and protein complex were released by resuspending the beads in 1x sample buffer supplemented with 0.1M DTT and incubated at 95°C for 5-7 min.

#### Electrophoresis Mobility Shift Assay (EMSA)

nELAVL SE and SA mutants were cloned and overexpressed in E.coli. Proteins were purified by Gene Universal Inc., (Chuzhou, Anhui). Fos-Cy5 au-rich RNA oligos (sequence: AUAUUUAUAUUUUUAUUUUAUUUUUUU) was synthesized by Techdragen. The binding buffer for EMSA consisted of 60 mM KCl, 300 mM NaCl, 2 mM MgCl2, 20 mM HEPES (pH 8.0) 1 mM EDTA, 1 mM DTT, 10 % Glycerol, 0.5 mg/mL of tRNA, and 2 fmol of Fos-Cy5 RNA per reaction. Reactions were equilibrated for 1 h at room temperature before loading on a Tris-glycine gel (to minimize complex dissociation), running 45 mins at 100 V. All gel shifts were done at least three times. Bands were quantified using ImageJ. Fraction bounds were calculated by plotting the ratio of RNA-protein complex against total RNA. Lines were obtained by nonlinear regression.

#### Puromycin Proximity Ligation Assay (Puro-PLA)

The Puro-PLA were performed on rat primary hippocampal neurons expressed with ELAVL3-SE/SA probing Fos protein and puromycin with kit (Duolink®C In Situ Red Starter Kit Mouse/Rabbit, catalog number: DUO92101, Sigma Aldrich) following manufacturer’s instructions.

#### Craniotomy and AAV injection

6-week-old mice were anesthetized with isoflurane mixed with air (3% for induction, 2% for surgery) and placed on a stereotaxic apparatus (RWD Instruments). A 38 °C heating pad was used for maintaining animal body temperature. Ophthalmic ointment (Bausch & Lomb corner gel) was applied to the eyes for eye lubrication. Ethanol and betadine swabs were used to sterilize the skin above the skull before making an incision. A ∼4 mm craniotomy window was made over the left V1 (center coordinates were 3.2 mm ML and 1 mm AP relative to lambda) by a dental drill as described previously^108^. A glass pipette (RWD Instruments) was pulled to a fine tip by a puller (PC-100, Narishige), and filled with mineral oil in the whole pipette and AAV in the front tip. The filled glass pipette was lowered into the V1. At a depth of 0.1-0.2 mm below the dura, virus was injected slowly (120nl in 5 min) into V1 using a microinjector (R480 Nanoliter injection pump, RWD Instruments). After the injection, pipette was kept for 5 minutes before being withdrawn to prevent the backflow of virus. A two-layer glass window was made by sticking a 3 mm and 5 mm-diameter cover glass (#1 thickness, Warner Instruments) using UV sensitive optical adhesive. The glued glass window was placed under slight downward pressure on the exposed brain. The 5 mm was attached to the skull and the 3 mm was fitted into the craniotomy. A tissue-friendly adhesive (3M Vetbond) was applied to seal around the surrounding skull. A custom-made stainless steel head plate was attached to the skull with dental adhesive resin cement. Dental cement was used to seal the rest of the exposed skull. Two-photon imaging was conducted 2 weeks after craniotomy window implantation.

#### Viral vectors

For *in vivo* calcium imaging, rAAV-hSyn-GCaMp6f-WPRE-hGH-polyA (titer: ∼5.7x10^12^vg/ml, BrainVTA Co.,Ltd) was used. For sparse labeling of neurons for eMAP, a mixture of diluted pENN.AAV.CamKII 0.4.Cre.SV40 (titer: 1.8x10^13^vg/ml, Addgene: 105558, 1:10000-20000 dilution in PBS) and cre-dependent EGFP virus (titer: 1x10^13^ vg/ml, Addgene: 51502) were used. nELAVL-WT AAV was generated with a mix (1:1:1) of ELAVL2-WT, ELAVL3-WT and ELAVL4-WT plasmid (titer: 2.45×10^13^vg/ml, Vigene Biosciences, lnc.). nELAVLs-SE AAV was generated with a mix (1:1:1) of ELAVL2-S119E, ELAVL3-S119E and ELAVL4-S131E plasmid (titer: 6.60×10^13^vg/ml, Vigene Biosciences, lnc.). nELAVLs-SA virus was generated with a mix (1:1:1) of ELAVL2-E119A, ELAVL3-E119A and ELAVL4-E131A plasmid (titer: 8.02×10^13^vg/ml, Vigene Biosciences, lnc.).

#### Two-photon in vivo imaging with visual stimuli

The Psychophysics Toolbox in MATLAB was used to generate drifting grating stimuli and natural scene^109,110^. For each stimuli trial, there was a 3s blank period (gray at mean luminance) followed by a 1s drifting sinusoidal grating (mean luminance: ∼50 lx) with three different spatial frequencies (0.08, 0.04 and 0.02 cycles per degree) at temporal frequency of 2 Hz followed a 4s blank period with 4s natural scene. Eight drifting directions were separated by 45 degrees, and ten trials were displayed. An LCD monitor (21.5-inch, 75 Hz refresh rate) was placed 18 cm centered in front of the mouse’s eyes to display the stimuli. The monitor covered a field of view of ∼106 degrees horizontal and ∼58 degrees vertical. Mice were head-fixed, and the left V1 was imaged by a two-photon microscope (Scientifica, U.K.) with a resonant scanning module controlled by ScanImage built-in Matlab software. The Coherent Discovery laser was used at 920 nm for excitation through a Nikon water immersion objective (16X, N.A. = 0.8, working distance: 3.0 mm). The power was less than 100 mW for all imaging. The signal was detected by a photomultiplier with bandwidth of 480-560nm. Images were acquired at ∼7.5 Hz (30 Hz averaged to 4) over 512x512 pixels of each view with bidirectional scanning. BNC DAQ (BNC-2110, National Instruments) was used to synchronize the two-photon microscope recording and visual stimuli display. Mice were habituated with the imaging set-up for two to three days (30 min daily) before data collection. Neurons of L2/3 at 100-250μm below dura were imaged. To record visually evoked responses from different eyes, an opaque black card blocker was placed in front of the eye according to imaging conditions. 3 to 6 fields of view (FOVs) were collected for each mouse.

#### Image processing and data analysis for two-photon microscopy

To extract the calcium signal of each neuron at different imaging conditions (i.e., spatial frequencies 0.02, 0.04 and 0.08cpd, and ways of eye blocking), all images of the same FOV were imported and analyzed by suite2p (version 0.10.1) at the same time. The parameters of suite2p were set as follow: tau of 0.7, denoise of 1, diameter of 9, anatomical only of 1, maximum iterations of 1, frames per second of 7.5 and 191 minimum neuropil pixels. To confirm the stereotaxic coordinates reliably targeted bV1, we ensured at least 10% of putative neurons (regions of interest (ROIs)) extracted from each imaging session elicited time-locked, visually evoked responses from the ipsi eye, similar to other published reports^60,65^.

The debris-like ROIs were removed manually (“*iscell*” from 1 to 0). The intensities of the remaining neurons were saved in .mat files for data analysis in MATLAB (version R2021a).

The raw intensities of neurons were adjusted with neuropil signal according to 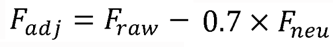^111^. Normalization of adjusted intensity was done by transforming into modified 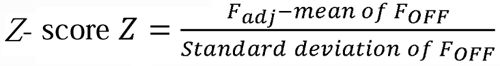, where *F_OFF_* is the 1-second adjusted signal in blank period just before each visual stimulus of that trial. The neuron with more than 20% of trials (i.e., > 2 trials in drifting grating and > 6 trials in natural scene) with Z > 2 at any stimulus, as well as the trial-average calculated for all time-point, Z_trial_avg_ >2 in that stimulus, was counted as stimulus-responsive. Tuning curve was obtained by averaging the Z_trial_avg_ in each grating direction. To analyze the orientation selectivity of grating-responsive neurons, the response R(θ_k_) of orientation *k* is calculated by averaging the Z_trial_avg_ of opposite directions. The orientation selectivity index (OSI) was defined as the norm of 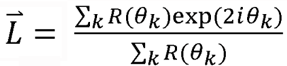 and the preferred orientation was half of the corresponding angle of 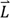 where *θ_k_* was expressed in radian^112^. The similarity of their signal was calculated as the distance correlation between their tuning curves for neuron responding to drifting grating stimuli in any two viewing conditions^113^. The confidence interval of mean difference of OSI distribution between WT and *Cdkl5* KO mice was estimated with bootstrapped distribution under significantly large sampling (i.e., 9999 times).

Considering that a neuron can contribute to population encoding regardless of their individual responses to the stimuli, all extracted putative neurons from suite2p were thus included in the population analysis. To quantify stimulus information encoded in activities of the neural populations, we measured the performance of a classifier in the prediction of different grating^114^. A support vector machine classifier was trained by 5-fold cross-validation^115^. Since the number of trials was limited in our study, the trained classifier would tend to overfit the training set, which could lead to an underestimation of the performance of the classifier for the validation set. Therefore, during training, the early stop was applied. Then the classifier’s performance was evaluated by accuracy on the validation set. To evaluate the binocular matching in the neural population, decoder was trained with contralateral data set and tested on both contra- and ipsilateral signal.

#### Visual cliff task

The visual cliff task assessed mice’s preference for descending a platform in an open field. The field consisted of an open field (40 x 40 x 40 cm) with transparent plexiglass floor positioned on the edge of a laboratory bench: half of the box rested on the tabletop (the “safe” side), while the other half extended over the floor 80 cm below (the “cliff” side)^57^. Both sides featured patterns visible through the plexiglass floor, a black and white checkerboard cover (3.5 x 3.5 cm squares) was placed on the bench top and extended vertically down to the floor, creating the illusion of a cliff. Prior to each trial, mice were transported to the testing room and habituated in their home cages for at least 30 minutes. At the start of each trial, a mouse was placed on a platform (3 cm in height, 5 cm in diameter) at the center of the box, facing the middle line between the two sides. The side from which the mouse descended was recorded by a camera. Each mouse underwent 5 trials, and the open field was cleaned between trials. The percentage of descents from the cliff side was calculated for each mouse.

## References

1 Montini, E. et al. Identification and characterization of a novel serine-threonine kinase gene from the Xp22 region. Genomics 51, 427–433, doi:10.1006/geno.1998.5391 (1998).

2 Kalscheuer, V. M. et al. Disruption of the serine/threonine kinase 9 gene causes severe X-linked infantile spasms and mental retardation. Am J Hum Genet 72, 1401–1411, doi:10.1086/375538 (2003).

3 Olson, H. E. et al. Cyclin-Dependent Kinase-Like 5 Deficiency Disorder: Clinical Review. Pediatr Neurol 97, 18–25, doi:10.1016/j.pediatrneurol.2019.02.015 (2019).

4 Leonard, H. et al. CDKL5 deficiency disorder: clinical features, diagnosis, and management. Lancet Neurol 21, 563–576, doi:10.1016/S1474-4422(22)00035-7 (2022).

5 Wang, I. T. et al. Loss of CDKL5 disrupts kinome profile and event-related potentials leading to autistic-like phenotypes in mice. Proc Natl Acad Sci U S A 109, 21516–21521, doi:10.1073/pnas.1216988110 (2012).

6 Mazziotti, R. et al. Searching for biomarkers of CDKL5 disorder: early-onset visual impairment in CDKL5 mutant mice. Hum Mol Genet 26, 2290–2298, doi:10.1093/hmg/ddx119 (2017).

7 Sampedro-Castaneda, M. et al. Epilepsy-linked kinase CDKL5 phosphorylates voltage-gated calcium channel Cav2.3, altering inactivation kinetics and neuronal excitability. Nat Commun 14, 7830, doi:10.1038/s41467-023-43475-w (2023).

8 Zhou, A., Han, S. & Zhou, Z. J. Molecular and genetic insights into an infantile epileptic encephalopathy - CDKL5 disorder. Front Biol (Beijing) 12, 1–6, doi:10.1007/s11515-016-1438-7 (2017).

9 Munoz, I. M. et al. Phosphoproteomic screening identifies physiological substrates of the CDKL5 kinase. EMBO J 37, doi:10.15252/embj.201899559 (2018).

10 Baltussen, L. L. et al. Chemical genetic identification of CDKL5 substrates reveals its role in neuronal microtubule dynamics. EMBO J 37, doi:10.15252/embj.201899763 (2018).

11 Khanam, T. et al. CDKL5 kinase controls transcription-coupled responses to DNA damage. EMBO J 40, e108271, doi:10.15252/embj.2021108271 (2021).

12 Kim, J. Y. et al. A kinome-wide screen identifies a CDKL5-SOX9 regulatory axis in epithelial cell death and kidney injury. Nat Commun 11, 1924, doi:10.1038/s41467-020-15638-6 (2020).

13 Lopes, A. T., Janiv, O., Claxton, S. & Ultanir, S. K. CDKL5’s role in microtubule-based transport and cognitive function. bioRxiv, 2024.2008.2028.610038, doi:10.1101/2024.08.28.610038 (2024).

14 De Santis, R. et al. Mutant FUS and ELAVL4 (HuD) Aberrant Crosstalk in Amyotrophic Lateral Sclerosis. Cell Rep 27, 3818–3831 e3815, doi:10.1016/j.celrep.2019.05.085 (2019).

15 Markmiller, S. et al. Context-Dependent and Disease-Specific Diversity in Protein Interactions within Stress Granules. Cell 172, 590–604 e513, doi:10.1016/j.cell.2017.12.032 (2018).

16 Bowles, K. R. et al. ELAVL4, splicing, and glutamatergic dysfunction precede neuron loss in MAPT mutation cerebral organoids. Cell 184, 4547–4563 e4517, doi:10.1016/j.cell.2021.07.003 (2021).

17 Bolognani, F., Contente-Cuomo, T. & Perrone-Bizzozero, N. I. Novel recognition motifs and biological functions of the RNA-binding protein HuD revealed by genome-wide identification of its targets. Nucleic Acids Res 38, 117–130, doi:10.1093/nar/gkp863 (2010).

18 Colombrita, C., Silani, V. & Ratti, A. ELAV proteins along evolution: back to the nucleus? Mol Cell Neurosci 56, 447–455, doi:10.1016/j.mcn.2013.02.003 (2013).

19 Ip, J. P. K., Mellios, N. & Sur, M. Rett syndrome: insights into genetic, molecular and circuit mechanisms. Nat Rev Neurosci 19, 368–382, doi:10.1038/s41583-018-0006-3 (2018).

20 Banerjee, A. et al. Jointly reduced inhibition and excitation underlies circuit-wide changes in cortical processing in Rett syndrome. Proc Natl Acad Sci U S A 113, E7287–E7296, doi:10.1073/pnas.1615330113 (2016).

21 Demarest, S. T. et al. CDKL5 deficiency disorder: Relationship between genotype, epilepsy, cortical visual impairment, and development. Epilepsia 60, 1733–1742, doi:10.1111/epi.16285 (2019).

22 Goel, A. et al. Impaired perceptual learning in a mouse model of Fragile X syndrome is mediated by parvalbumin neuron dysfunction and is reversible. Nat Neurosci 21, 1404–1411, doi:10.1038/s41593-018-0231-0 (2018).

23 Townsend, L. B., Jones, K. A., Dorsett, C. R., Philpot, B. D. & Smith, S. L. Deficits in higher visual area representations in a mouse model of Angelman syndrome. J Neurodev Disord 12, 28, doi:10.1186/s11689-020-09329-y (2020).

24 Galli, J. et al. Neurovisual profile in children affected by Angelman syndrome. Brain Dev 45, 117–125, doi:10.1016/j.braindev.2022.10.003 (2023).

25 Boggio, E. M. et al. Visual impairment in FOXG1-mutated individuals and mice. Neuroscience 324, 496–508, doi:10.1016/j.neuroscience.2016.03.027 (2016).

26 Olson, H. E. et al. Cerebral visual impairment in CDKL5 deficiency disorder: vision as an outcome measure. Dev Med Child Neurol 63, 1308–1315, doi:10.1111/dmcn.14908 (2021).

27 Brock, D. et al. Cerebral Visual Impairment in CDKL5 Deficiency Disorder Correlates With Developmental Achievement. J Child Neurol 36, 974–980, doi:10.1177/08830738211019284 (2021).

28 Lupori, L. et al. Site-specific abnormalities in the visual system of a mouse model of CDKL5 deficiency disorder. Hum Mol Genet 28, 2851–2861, doi:10.1093/hmg/ddz102 (2019).

29 Pizzo, R. et al. Lack of Cdkl5 Disrupts the Organization of Excitatory and Inhibitory Synapses and Parvalbumin Interneurons in the Primary Visual Cortex. Front Cell Neurosci 10, 261, doi:10.3389/fncel.2016.00261 (2016).

30 Quintiliani, M. et al. Cortical Visual Impairment in CDKL5 Deficiency Disorder. Front Neurol 12, 805745, doi:10.3389/fneur.2021.805745 (2021).

31 Eyers, P. A. A new consensus for evaluating CDKL5/STK9-dependent signalling mechanisms. EMBO J 37, doi:10.15252/embj.2018100848 (2018).

32 Mulligan, M. R. & Bicknell, L. S. The molecular genetics of nELAVL in brain development and disease. Eur J Hum Genet 31, 1209–1217, doi:10.1038/s41431-023-01456-z (2023).

33 Scheckel, C. et al. Regulatory consequences of neuronal ELAV-like protein binding to coding and non-coding RNAs in human brain. Elife 5, doi:10.7554/eLife.10421 (2016).

34 Crair, M. C., Gillespie, D. C. & Stryker, M. P. The role of visual experience in the development of columns in cat visual cortex. Science 279, 566–570, doi:10.1126/science.279.5350.566 (1998).

35 Wolock, S. L., Lopez, R. & Klein, A. M. Scrublet: Computational Identification of Cell Doublets in Single-Cell Transcriptomic Data. Cell Syst 8, 281–291 e289, doi:10.1016/j.cels.2018.11.005 (2019).

36 Korsunsky, I. et al. Fast, sensitive and accurate integration of single-cell data with Harmony. Nat Methods 16, 1289–1296, doi:10.1038/s41592-019-0619-0 (2019).

37 Cheng, S. et al. Vision-dependent specification of cell types and function in the developing cortex. Cell 185, 311–327 e324, doi:10.1016/j.cell.2021.12.022 (2022).

38 Kim, E. J. et al. Extraction of Distinct Neuronal Cell Types from within a Genetically Continuous Population. Neuron 107, 274–282 e276, doi:10.1016/j.neuron.2020.04.018 (2020).

39 O’Toole, S. M., Oyibo, H. K. & Keller, G. B. Molecularly targetable cell types in mouse visual cortex have distinguishable prediction error responses. Neuron 111, 2918–2928 e2918, doi:10.1016/j.neuron.2023.08.015 (2023).

40 Jenks, K. R. et al. Arc restores juvenile plasticity in adult mouse visual cortex. Proc Natl Acad Sci U S A 114, 9182–9187, doi:10.1073/pnas.1700866114 (2017).

41 Brakeman, P. R. et al. Homer: a protein that selectively binds metabotropic glutamate receptors. Nature 386, 284–288, doi:10.1038/386284a0 (1997).

42 Barber, C. F., Jorquera, R. A., Melom, J. E. & Littleton, J. T. Postsynaptic regulation of synaptic plasticity by synaptotagmin 4 requires both C2 domains. J Cell Biol 187, 295–310, doi:10.1083/jcb.200903098 (2009).

43 Aiken, J., Buscaglia, G., Bates, E. A. & Moore, J. K. The alpha-Tubulin gene TUBA1A in Brain Development: A Key Ingredient in the Neuronal Isotype Blend. J Dev Biol 5, doi:10.3390/jdb5030008 (2017).

44 Gu, Y. et al. Obligatory role for the immediate early gene NARP in critical period plasticity. Neuron 79, 335–346, doi:10.1016/j.neuron.2013.05.016 (2013).

45 Kwapis, J. L. et al. Epigenetic regulation of the circadian gene Per1 contributes to age-related changes in hippocampal memory. Nat Commun 9, 3323, doi:10.1038/s41467-018-05868-0 (2018).

46 Paradis, S. et al. An RNAi-based approach identifies molecules required for glutamatergic and GABAergic synapse development. Neuron 53, 217–232, doi:10.1016/j.neuron.2006.12.012 (2007).

47 Ban, Y. et al. Prickle promotes the formation and maintenance of glutamatergic synapses by stabilizing the intercellular planar cell polarity complex. Sci Adv 7, eabh2974, doi:10.1126/sciadv.abh2974 (2021).

48 Li, M. et al. CDKL5 modulates structural plasticity of excitatory synapses via liquid-liquid phase separation. *bioRxiv*, 2025.2003.2016.643581, doi:10.1101/2025.03.16.643581 (2025).

49 Cai, H., Vernon, R. M. & Forman-Kay, J. D. An interpretable machine learning algorithm to predict disordered protein phase separation based on biophysical interactions. bioRxiv, 2022.2007.2006.499043, doi:10.1101/2022.07.06.499043 (2022).

50 Li, J. et al. Post-translational modifications in liquid-liquid phase separation: a comprehensive review. Mol Biomed 3, 13, doi:10.1186/s43556-022-00075-2 (2022).

51 Ripin, N. & Parker, R. Formation, function, and pathology of RNP granules. Cell 186, 4737–4756, doi:10.1016/j.cell.2023.09.006 (2023).

52 Youn, J. Y. et al. Properties of Stress Granule and P-Body Proteomes. Mol Cell 76, 286–294, doi:10.1016/j.molcel.2019.09.014 (2019).

53 Hubstenberger, A. et al. P-Body Purification Reveals the Condensation of Repressed mRNA Regulons. Mol Cell 68, 144–157 e145, doi:10.1016/j.molcel.2017.09.003 (2017).

54 Wilbertz, J. H. et al. Single-Molecule Imaging of mRNA Localization and Regulation during the Integrated Stress Response. Mol Cell 73, 946–958 e947, doi:10.1016/j.molcel.2018.12.006 (2019).

55 Kodali, S. et al. RNA sequestration in P-bodies sustains myeloid leukaemia. Nat Cell Biol 26, 1745–1758, doi:10.1038/s41556-024-01489-6 (2024).

56 Sato, K. & Kotani, T. Visualizing the translational activation of a particular mRNA in zebrafish embryos using in situ hybridization and proximity ligation assay. STAR Protoc 5, 102951, doi:10.1016/j.xpro.2024.102951 (2024).

57. De Jesus-Cortes, H., et al. Using the visual cliff and pole descent assays to detect binocular disruption in mice. bioRxiv, doi:10.1101/2023.05.29.542767 (2024).

58 Boone, H. C. et al. Natural binocular depth discrimination behavior in mice explained by visual cortical activity. Curr Biol 31, 2191–2198 e2193, doi:10.1016/j.cub.2021.02.031 (2021).

59 Chen, T. W. et al. Ultrasensitive fluorescent proteins for imaging neuronal activity. Nature 499, 295–300, doi:10.1038/nature12354 (2013).

60 Tan, L., Tring, E., Ringach, D. L., Zipursky, S. L. & Trachtenberg, J. T. Vision Changes the Cellular Composition of Binocular Circuitry during the Critical Period. Neuron 108, 735–747 e736, doi:10.1016/j.neuron.2020.09.022 (2020).

61 Chang, J. T., Whitney, D. & Fitzpatrick, D. Experience-Dependent Reorganization Drives Development of a Binocularly Unified Cortical Representation of Orientation. Neuron 107, 338–350 e335, doi:10.1016/j.neuron.2020.04.022 (2020).

62 Tan, L. M., Ringach, D. L. & Trachtenberg, J. T. The Development of Receptive Field Tuning Properties in Mouse Binocular Primary Visual Cortex. Journal of Neuroscience 42, doi:10.1523/Jneurosci.1702-21.2022 (2022).

63 Espinosa, J. S. & Stryker, M. P. Development and plasticity of the primary visual cortex. Neuron 75, 230–249, doi:10.1016/j.neuron.2012.06.009 (2012).

64 Wang, B. S., Sarnaik, R. & Cang, J. Critical period plasticity matches binocular orientation preference in the visual cortex. Neuron 65, 246–256, doi:10.1016/j.neuron.2010.01.002 (2010).

65 Jenks, K. R. & Shepherd, J. D. Experience-Dependent Development and Maintenance of Binocular Neurons in the Mouse Visual Cortex. Cell Rep 30, 1982–1994 e1984, doi:10.1016/j.celrep.2020.01.031 (2020).

66 Smith, S. L. & Trachtenberg, J. T. Experience-dependent binocular competition in the visual cortex begins at eye opening. Nat Neurosci 10, 370–375, doi:10.1038/nn1844 (2007).

67 Ukita, J. Causal importance of low-level feature selectivity for generalization in image recognition. Neural Netw 125, 185–193, doi:10.1016/j.neunet.2020.02.009 (2020).

68 Sun, Y. J., Espinosa, J. S., Hoseini, M. S. & Stryker, M. P. Experience-dependent structural plasticity at pre- and postsynaptic sites of layer 2/3 cells in developing visual cortex. Proc Natl Acad Sci U S A 116, 21812–21820, doi:10.1073/pnas.1914661116 (2019).

69 Song, J. Molecular Mechanisms of Phase Separation and Amyloidosis of ALS/FTD-linked FUS and TDP-43. Aging Dis 15, 2084–2112, doi:10.14336/AD.2023.1118 (2024).

70 Liu, N., Guo, Y., Ning, S. & Duan, M. Phosphorylation regulates the binding of intrinsically disordered proteins via a flexible conformation selection mechanism. Commun Chem 3, 123, doi:10.1038/s42004-020-00370-5 (2020).

71 Maharana, S. et al. RNA buffers the phase separation behavior of prion-like RNA binding proteins. Science 360, 918–921, doi:10.1126/science.aar7366 (2018).

72 Di Stefano, B. et al. The RNA Helicase DDX6 Controls Cellular Plasticity by Modulating P-Body Homeostasis. Cell Stem Cell 25, 622–638 e613, doi:10.1016/j.stem.2019.08.018 (2019).

73 Courel, M. et al. GC content shapes mRNA storage and decay in human cells. Elife 8, doi:10.7554/eLife.49708 (2019).

74 Marcelo, A., Koppenol, R., de Almeida, L. P., Matos, C. A. & Nobrega, C. Stress granules, RNA-binding proteins and polyglutamine diseases: too much aggregation? Cell Death Dis 12, 592, doi:10.1038/s41419-021-03873-8 (2021).

75 Sheth, U. & Parker, R. Decapping and decay of messenger RNA occur in cytoplasmic processing bodies. Science 300, 805–808, doi:10.1126/science.1082320 (2003).

76 Dar, S. A. et al. Full-length direct RNA sequencing uncovers stress granule-dependent RNA decay upon cellular stress. Elife 13, doi:10.7554/eLife.96284 (2024).

77 Blake, L. A., Watkins, L., Liu, Y., Inoue, T. & Wu, B. A rapid inducible RNA decay system reveals fast mRNA decay in P-bodies. Nat Commun 15, 2720, doi:10.1038/s41467-024-46943-z (2024).

78 Hallacli, E. et al. The Parkinson’s disease protein alpha-synuclein is a modulator of processing bodies and mRNA stability. Cell 185, 2035–2056 e2033, doi:10.1016/j.cell.2022.05.008 (2022).

79 Bentmann, E., Haass, C. & Dormann, D. Stress granules in neurodegeneration--lessons learnt from TAR DNA binding protein of 43 kDa and fused in sarcoma. FEBS J 280, 4348–4370, doi:10.1111/febs.12287 (2013).

80 Wolozin, B. & Ivanov, P. Stress granules and neurodegeneration. Nat Rev Neurosci 20, 649–666, doi:10.1038/s41583-019-0222-5 (2019).

81 Ince-Dunn, G. et al. Neuronal Elav-like (Hu) proteins regulate RNA splicing and abundance to control glutamate levels and neuronal excitability. Neuron 75, 1067–1080, doi:10.1016/j.neuron.2012.07.009 (2012).

82 Akamatsu, W. et al. The RNA-binding protein HuD regulates neuronal cell identity and maturation. Proc Natl Acad Sci U S A 102, 4625–4630, doi:10.1073/pnas.0407523102 (2005).

83 DeBoer, E. M. et al. Prenatal deletion of the RNA-binding protein HuD disrupts postnatal cortical circuit maturation and behavior. J Neurosci 34, 3674–3686, doi:10.1523/JNEUROSCI.3703-13.2014 (2014).

84 Berto, S., Usui, N., Konopka, G. & Fogel, B. L. ELAVL2-regulated transcriptional and splicing networks in human neurons link neurodevelopment and autism. Hum Mol Genet 25, 2451–2464, doi:10.1093/hmg/ddw110 (2016).

85 Zhou, X. et al. Integrating de novo and inherited variants in 42,607 autism cases identifies mutations in new moderate-risk genes. Nat Genet 54, 1305–1319, doi:10.1038/s41588-022-01148-2 (2022).

86 Satterstrom, F. K. et al. Large-Scale Exome Sequencing Study Implicates Both Developmental and Functional Changes in the Neurobiology of Autism. Cell 180, 568–584 e523, doi:10.1016/j.cell.2019.12.036 (2020).

87 Rodin, R. E. et al. The landscape of somatic mutation in cerebral cortex of autistic and neurotypical individuals revealed by ultra-deep whole-genome sequencing. Nat Neurosci 24, 176–185, doi:10.1038/s41593-020-00765-6 (2021).

88 Jia, X. et al. De novo variants in genes regulating stress granule assembly associate with neurodevelopmental disorders. Sci Adv 8, eabo7112, doi:10.1126/sciadv.abo7112 (2022).

89 Eising, E. et al. A set of regulatory genes co-expressed in embryonic human brain is implicated in disrupted speech development. Mol Psychiatry 24, 1065–1078, doi:10.1038/s41380-018-0020-x (2019).

90 Andrews, S. (Babraham Bioinformatics, Babraham Institute, Cambridge, United Kingdom, 2010).

91 Kim, D., Paggi, J. M., Park, C., Bennett, C. & Salzberg, S. L. Graph-based genome alignment and genotyping with HISAT2 and HISAT-genotype. Nat Biotechnol 37, 907–915, doi:10.1038/s41587-019-0201-4 (2019).

92 Liao, Y., Smyth, G. K. & Shi, W. featureCounts: an efficient general purpose program for assigning sequence reads to genomic features. Bioinformatics 30, 923–930, doi:10.1093/bioinformatics/btt656 (2014).

93 Love, M. I., Huber, W. & Anders, S. Moderated estimation of fold change and dispersion for RNA-seq data with DESeq2. Genome Biol 15, 550, doi:10.1186/s13059-014-0550-8 (2014).

94 Yu, G., Wang, L. G., Han, Y. & He, Q. Y. clusterProfiler: an R package for comparing biological themes among gene clusters. OMICS 16, 284–287, doi:10.1089/omi.2011.0118 (2012).

95 Lau, S. F., Cao, H., Fu, A. K. Y. & Ip, N. Y. Single-nucleus transcriptome analysis reveals dysregulation of angiogenic endothelial cells and neuroprotective glia in Alzheimer’s disease. Proc Natl Acad Sci U S A 117, 25800–25809, doi:10.1073/pnas.2008762117 (2020).

96 Wolf, F. A., Angerer, P. & Theis, F. J. SCANPY: large-scale single-cell gene expression data analysis. Genome Biol 19, 15, doi:10.1186/s13059-017-1382-0 (2018).

97 Traag, V. A., Waltman, L. & van Eck, N. J. From Louvain to Leiden: guaranteeing well-connected communities. Sci Rep 9, 5233, doi:10.1038/s41598-019-41695-z (2019).

98 Becht, E. et al. Dimensionality reduction for visualizing single-cell data using UMAP. Nat Biotechnol, doi:10.1038/nbt.4314 (2018).

99 Hrvatin, S. et al. Single-cell analysis of experience-dependent transcriptomic states in the mouse visual cortex. Nat Neurosci 21, 120–129, doi:10.1038/s41593-017-0029-5 (2018).

100 Choi, H. M., Beck, V. A. & Pierce, N. A. Next-generation in situ hybridization chain reaction: higher gain, lower cost, greater durability. ACS Nano 8, 4284–4294, doi:10.1021/nn405717p (2014).

101 Ku, T. et al. Multiplexed and scalable super-resolution imaging of three-dimensional protein localization in size-adjustable tissues. Nat Biotechnol 34, 973–981, doi:10.1038/nbt.3641 (2016).

102 Park, J. et al. Epitope-preserving magnified analysis of proteome (eMAP). Sci Adv 7, eabf6589, doi:10.1126/sciadv.abf6589 (2021).

103 Zhao, J. et al. Specific depletion of the motor protein KIF5B leads to deficits in dendritic transport, synaptic plasticity and memory. Elife 9, doi:10.7554/eLife.53456 (2020).

104 Reikofski, J. & Tao, B. Y. Polymerase chain reaction (PCR) techniques for site-directed mutagenesis. Biotechnol Adv 10, 535–547, doi:10.1016/0734-9750(92)91451-j (1992).

105 Fisher, C. L. & Pei, G. K. Modification of a PCR-based site-directed mutagenesis method. Biotechniques 23, 570–571, 574, doi:10.2144/97234bm01 (1997).

106 Ip, J. P. et al. alpha2-chimaerin controls neuronal migration and functioning of the cerebral cortex through CRMP-2. Nat Neurosci 15, 39–47, doi:10.1038/nn.2972 (2011).

107 Ye, T. et al. Efficient manipulation of gene dosage in human iPSCs using CRISPR/Cas9 nickases. Commun Biol 4, 195, doi:10.1038/s42003-021-01722-0 (2021).

108 El-Boustani, S. et al. Locally coordinated synaptic plasticity of visual cortex neurons in vivo. Science 360, 1349–1354, doi:10.1126/science.aao0862 (2018).

109 Brainard, D. H. The Psychophysics Toolbox. Spat Vis 10, 433–436 (1997).

110 Pelli, D. G. The VideoToolbox software for visual psychophysics: Transforming numbers into movies. Spatial Vision 10, 437–442, doi:Doi 10.1163/156856897x00366 (1997).

111 Stringer, C., Pachitariu, M., Steinmetz, N., Carandini, M. & Harris, K. D. High-dimensional geometry of population responses in visual cortex. Nature 571, 361–365, doi:10.1038/s41586-019-1346-5 (2019).

112 Mazurek, M., Kager, M. & Van Hooser, S. D. Robust quantification of orientation selectivity and direction selectivity. Front Neural Circuits 8, 92, doi:10.3389/fncir.2014.00092 (2014).

113 Székely, G. J., Rizzo, M. L. & Bakirov, N. K. Measuring and testing dependence by correlation of distances. The Annals of Statistics 35, 2769–2794, 2726 (2007).

114 Kafashan, M. et al. Scaling of sensory information in large neural populations shows signatures of information-limiting correlations. Nat Commun 12, 473, doi:10.1038/s41467-020-20722-y (2021).

115 Pedregosa, F. et al. Scikit-learn: Machine learning in Python. the Journal of machine Learning research 12, 2825–2830 (2011).

