## Supplemental Figure for "nELAVL phosphorylation by CDKL5 regulates RNA metabolism and condensates communication to promote experience-dependent maturation of the visual cortex"

Supplementary Data File includes:

• Supplementary Figures S1-S10;

• Supplementary Tables S1-S5 title.

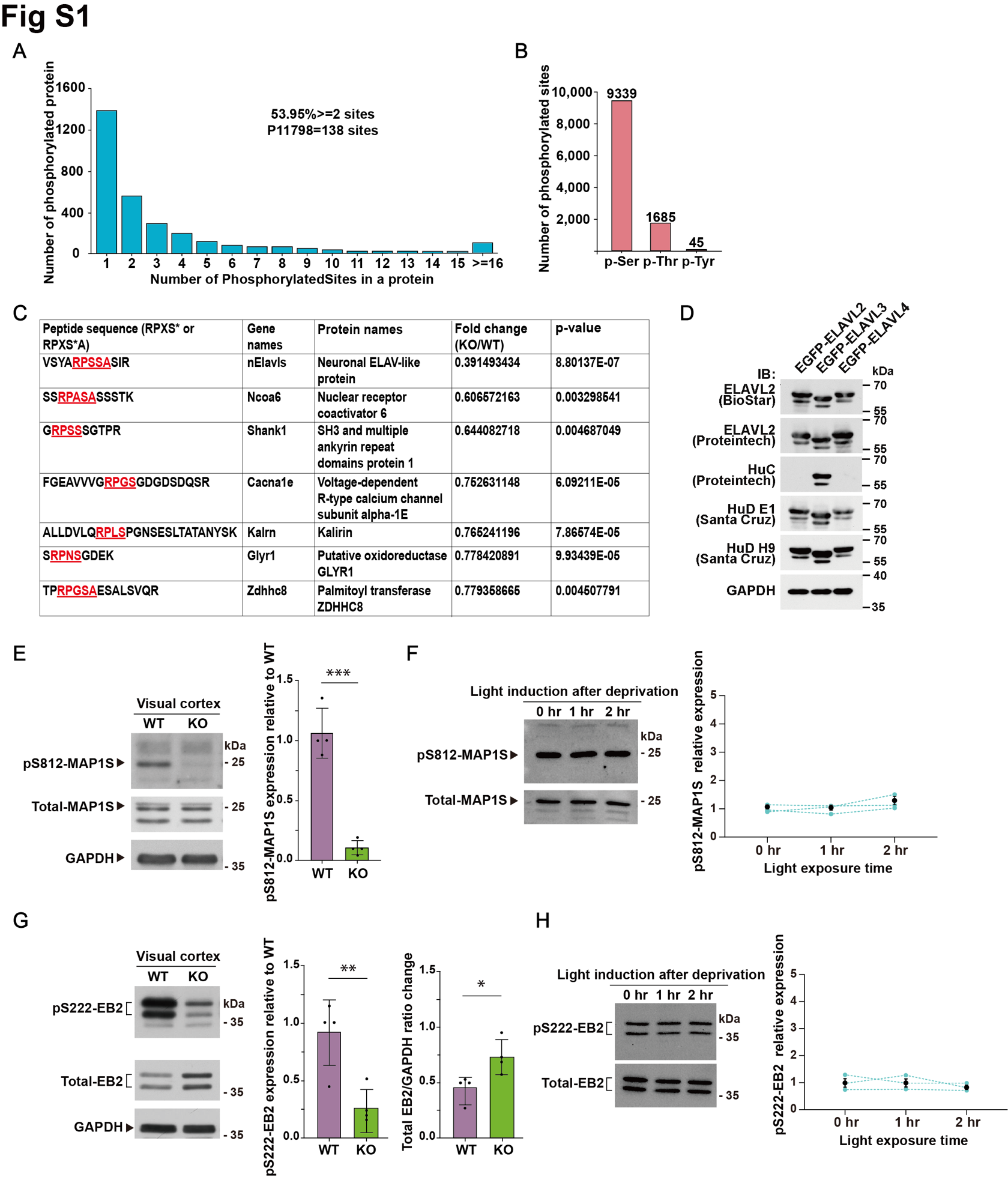

**Figure S1. Phosphoproteomic screening and validation in V1 of WT and *Cdkl5* KO mice.**

(A) Histogram showing the distribution of the number of phosphorylation sites per protein.

(B) The number of Ser, Thr, and Tyr phosphorylation sites in the identified phosphopeptides.

(C) The downregulated substrates with the RPXS*A/RPXS* motif were obtained through phosphoproteomic screening.

(D) ELAVL2 (HuB), ELAVL3 (HuC) and ELAVL4 (HuD) antibodies specificity validation by overexpression of ELAVL2, 3, 4 in HEK 293T cells. Note that we tested five commercially available ELAVL2, 3, and 4 antibodies and found that one of them is specific to ELAVL3, while the other antibodies target pan-nELAVLs.

(E) MAP1S phosphorylation was reduced in *Cdkl5* KO mice V1 lysates, as revealed with phospho-specific antibodies and quantification. n=4 mice, Unpaired t-test (two-tailed).

(F) Similar levels of MAP1S phosphorylation in dark environment (16 hr) with 0 hr, 1 hr and 2 hr light induction in visual cortex in WT mice and quantification. n=3 mice. Unpaired t-test (two-tailed). Dot lines indicate same batch of experiments.

(G) EB2 phosphorylation was reduced in *Cdkl5* KO mice V1 lysates, as revealed with phospho-specific antibodies and quantification. Quantification of phosphorylated EB2 is normalized to WT mice. n=4 mice, Unpaired t-test (two-tailed).

(H) Similar levels of EB2 phosphorylation in dark environment (16 hr) with 0 hr, 1 hr and 2 hr light induction in visual cortex in WT mice and quantification. n=3 mice. Unpaired t-test (two-tailed). Dot lines indicate same batch of experiments. Data are presented as mean ± SEM, * p < 0.05. ** p < 0.01. *** p < 0.001.

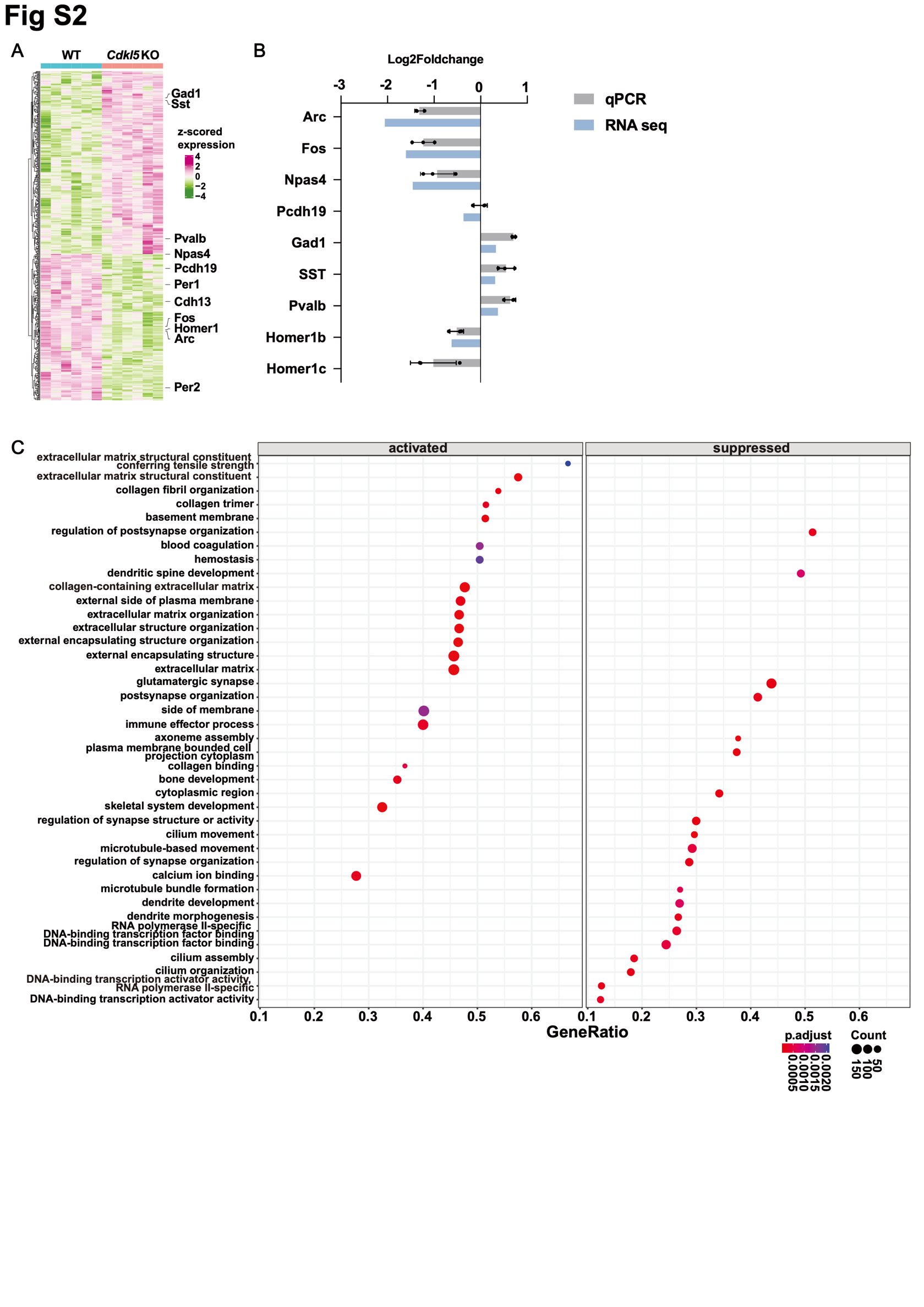

**Figure S2. Bulk RNA sequencing of WT and *Cdkl5* KO mice.**

(A) Heatmap showing the normalized expression of DEGs between WT and *Cdkl5* KO mouse V1. There were 272 upregulated and 218 downregulated genes. The DEGs of interest were labeled.

(B)Validation of eight DEGs detected by RNA sequencing using qRT-PCR. RNA sequencing values were shown in blue and qPCR values were shown in grey. Standard error of the mean was indicated by the black bars.

(C) GSEA enrichment plot showing top 20 terms that were significantly affected in *Cdkl5* KO mouse V1.

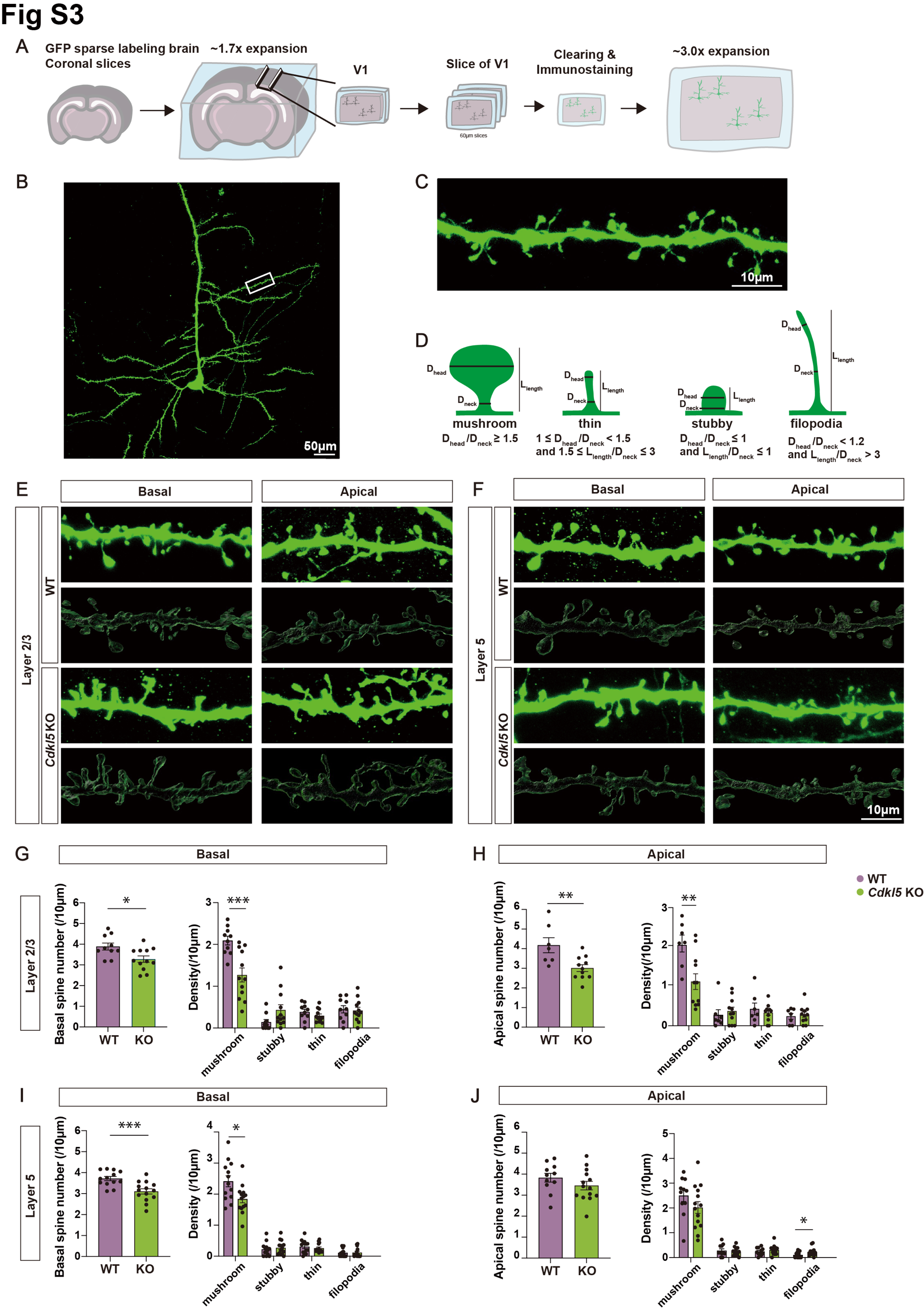

**Figure S3. Dendritic spine density and morphology analysis in WT and *Cdkl5* KO mice using eMAP.**

(A) Schematic diagram of tissue expansion, clearing and staining of eMAP.

(B) Confocal image of a layer 5 pyramidal neuron in WT. Scale bar, 50 μm.

(C) Magnified neuron dendritic segment in the white boxed in (B). Scale bar, 10 μm.

(D) An illustration of criteria to classify dendritic spines morphology. Dendritic spines were classified into four categories based on their morphology.

(E & F) GFP staining images and Imaris 3D reconstruction of basal and apical dendrites of layer 2/3 and layer 5 pyramidal neurons from WT and *Cdkl5* KO mice. Scale bar, 10 μm.

(G-J) The spine density in basal and apical dendrites of layer 2/3 and layer 5 pyramidal neurons from WT and *Cdkl5* KO mice. Data points indicate neurons from 3 mice per group. Layer 2/3: n=7-10 neurons from 3 WT mice and n=10-12 neurons from 3 WT mice. Layer 5: n=11-13 neurons from 3 WT mice and n=14 neurons from 3 WT mice. Unpaired t-test (two-tailed). Data are presented as mean ± SEM, * p < 0.05. ** p < 0.01. *** p < 0.001.

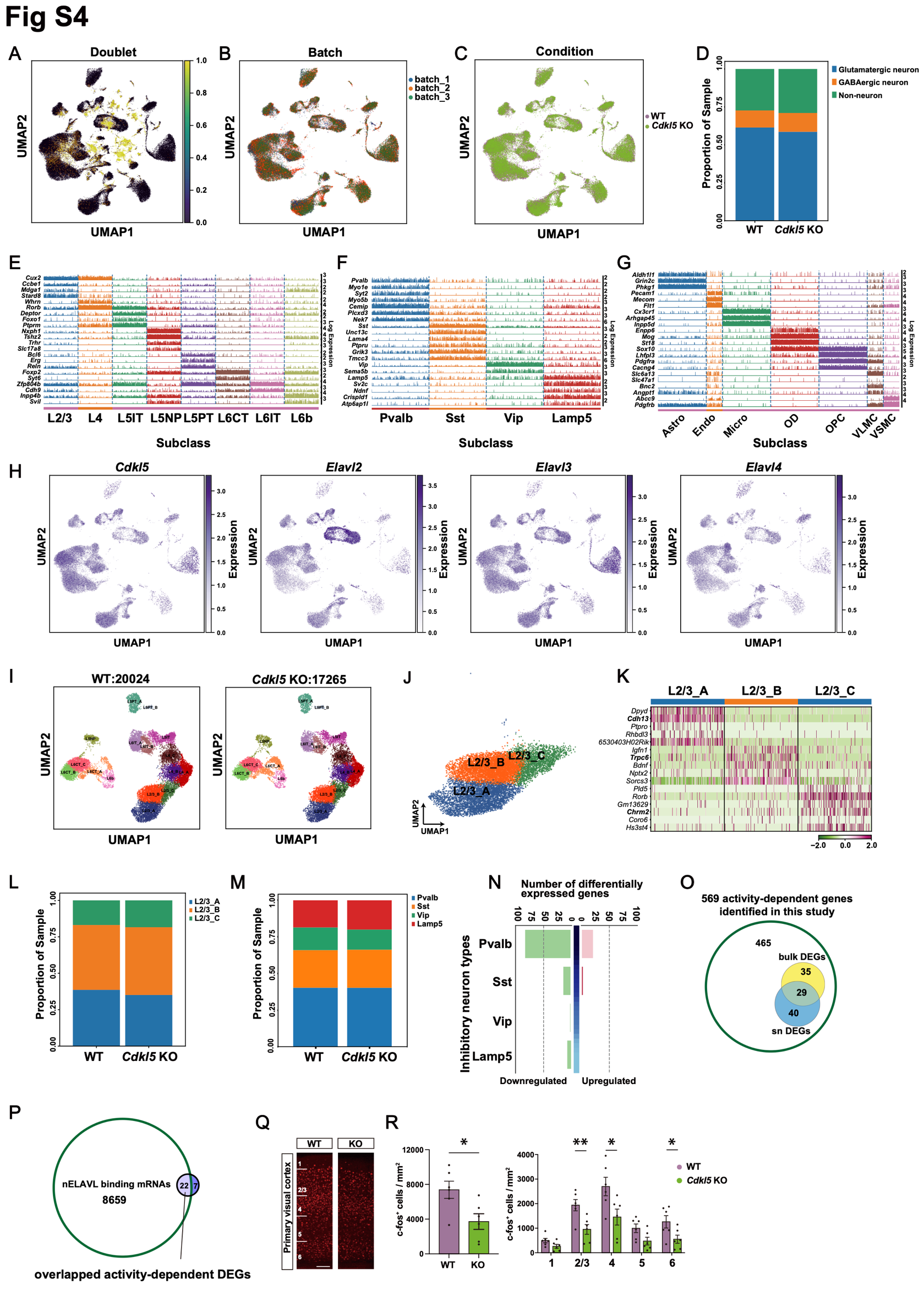

**Figure S4. Single nucleus transcriptomic profiling of WT and *Cdkl5* KO mice.**

(A) UMAP showing doublets (yellow) which were identified by Scrublet.

(B) UMAP showing batches of independent experiments in this study after batch correction.

(C) Same as (B), colored by Condition (WT and *Cdkl5* KO).

(D) Bar plot showing proportion of glutamatergic neurons, GABAergic neurons and non-neuronal cells in WT and *Cdkl5* KO mouse V1.

(E - G) Tracks plot showing subclass-specific markers (rows) in glutamatergic neurons (E), GABAergic neurons (F) and non-neuronal cells (G).

(H) UMAP showing the distribution of *Cdkl5* and *nElavls* gene expression across WT and *Cdkl5* KO mouse V1.

(I) UMAP visualizing annotations for subtypes of glutamatergic neurons in WT and *Cdkl5* KO mouse V1.

(J) UMAP visualizing sublayers (A, B, and C) in Layer 2/3 glutamatergic neurons of WT and *Cdkl5* KO mouse V1.

(K) Heatmap showing the z-scored expression of top five enriched genes for each sublayer of Layer 2/3 glutamatergic neurons.

(L) Bar plot showing the proportion of individual sublayer in Layer 2/3 glutamatergic neurons of WT and *Cdkl5* KO mouse V1.

(M) Bar plot showing proportion of GABAergic neurons including Pvalb, Sst, Vip and Lamp5 in WT and *Cdkl5* KO mouse V1.

(N) Number of DEGs between WT and *Cdkl5* KO mice V1 in GABAergic neurons, genes up- and down-regulated in *Cdkl5* KO compared with WT shown in pink and green bars, respectively. DEGs were defined as genes with |log_2_ fold change| > 0.25 and FDR-adjusted p-value < 0.05.

(O) Venn Diagram of activity-dependent genes identified in this study and overlapping numbers of DEGs in bulk RNA sequencing (yellow) and snRNA sequencing (blue).

(P) Twenty-two overlapped mRNAs between 8681 nELAVL binding genes on human brain (Scheckel et al., 2016) and 29 overlapped activity-dependent DEGs between bulk and sn sequencing from figure (O).

(Q & R) Decreased c-fos+ cells in L2/3, L4 and L6 in *Cdkl5* KO mouse V1. Scale bar, 100μm. WT: n=6 mice, *Cdkl5* KO: n=6 mice, Unpaired t-test (two-tailed). Data are presented as mean ± SEM, * p < 0.05. ** p < 0.01.

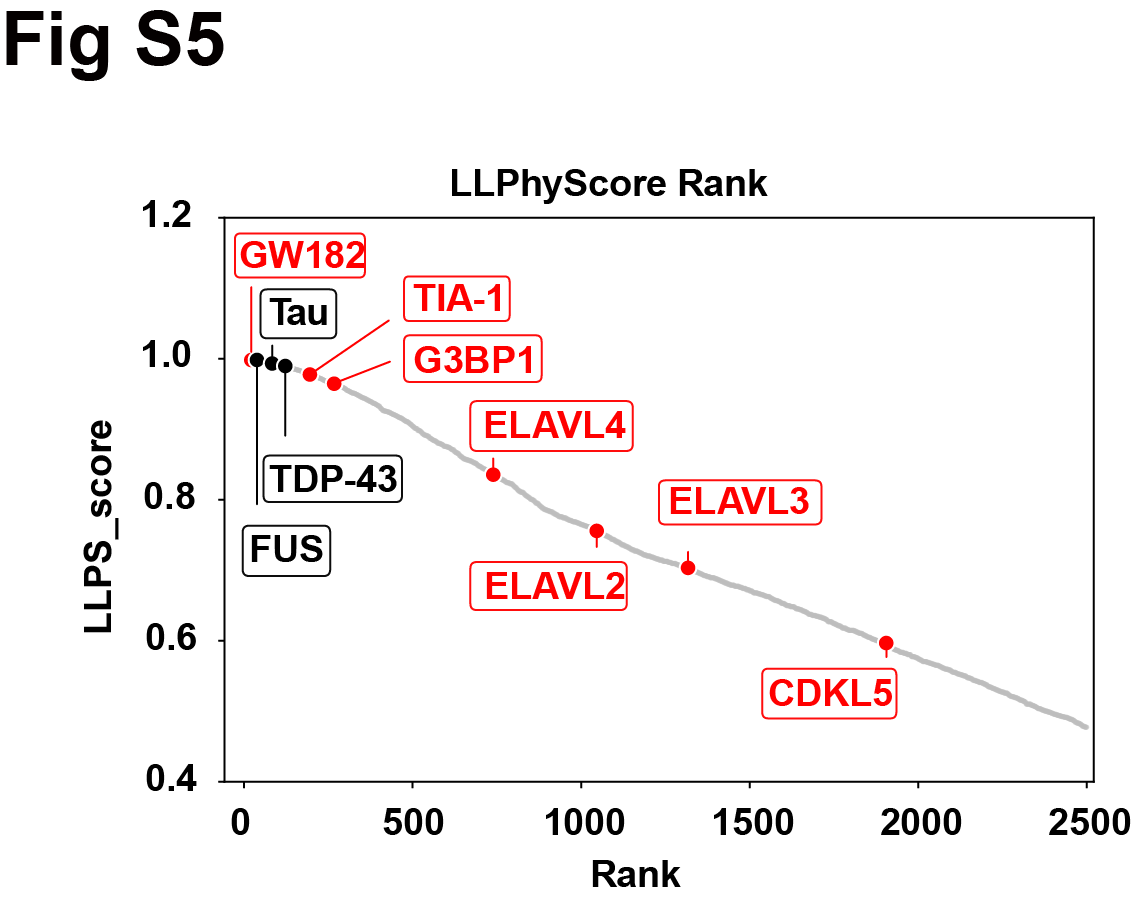

**Figure S5.** LLPhyScore analysis showing the phase separation scores and rank of nELAVLs and other cell condensates scaffold proteins such as GW182 and TIA1.

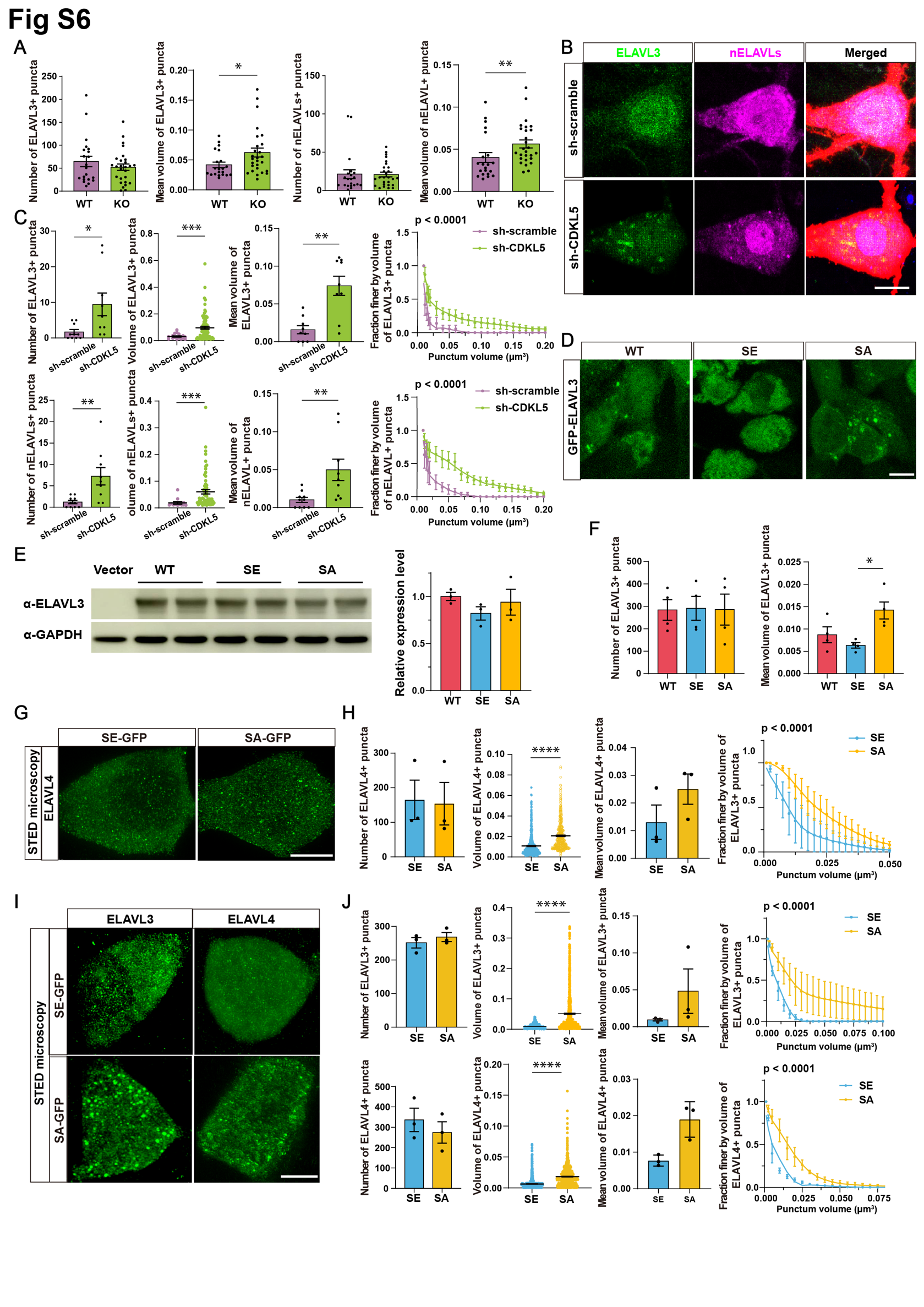

**Figure S6. Lack of phosphorylation by CDKL5 leads to altered nELAVL condensation.**

(A) Quantification analysis of ELAVL3 (magenta) and nELAVL (red) puncta in Figure 3A. While the puncta number showed no significant differences, the mean volume of ELAVL3 (magenta) and nELAVL (red) puncta showed significant increase in *Cdkl5* KO mice. WT: n=21 neurons from 3 mice; *Cdkl5* KO: n=27 neurons from 3 mice. Mann-Whitney test (two-tailed).

(B & C) Confocal imaging of ELAVL3 (green) and nELAVL (magenta) puncta in rat primary neurons with RFP denote sh-scramble or sh-CDKL5 transfection. Compared to sh-scramble control, sh-CDKL5 neurons harbored more and larger ELAVL3 (green) and nELAVL (magenta) puncta. Scale bar, 5 μm. n=10 (sh-scramble) and 9 (sh-CDKL5) cells from 3 batches of experiments. Mann-Whitney test (two-tailed). Volume fraction analysis: Two-way ANOVA with Sidak’s multiple comparisons test on main effect of sh-scramble and sh-CDKL5.

(D) Representative images of ELAVL3-WT/SE/SA overexpressing 293T cells. Scale bar, 10 μm.

(E) Exogenous ELAVL3 expression levels (left), and quantification of WB bands (right). n=3. One-way ANOVA followed by Tukey’s multiple comparisons test.

(F) Quantification analysis of ELAVL3 puncta in Figure 3C. While the puncta number showed no significant differences, the mean volume of ELAVL3 showed significant increase in SA group, compared to SE group. n=4 cells from 3 batches of experiments. One-way ANOVA followed by Turkey’s multiple comparisons test.

(G & H) STED microscopy of overexpressed SE or SA mutant of ELAVL4 in rat primary neurons. Compared to SE, the SA group had large-shifted punctum volume distribution. Scale bar, 5μm. n=3 cells from 3 baches of experiments. Mann-Whitney test (two-tailed). Volume fraction analysis: Two-way ANOVA with Sidak’s multiple comparisons test on main effect of SE and SA.

(I & J) STED microscopy of overexpressed SE or SA mutant of ELAVL3 and ELAVL4 in HEK 293T cells. Compared to SE, the SA mutant had large-shifted punctum volume distribution. Scale bar, 10 μm. n=3 cells from 3 batches of experiments. Mann-Whitney test (two-tailed). Volume fraction analysis: Two-way ANOVA with Sidak’s multiple comparisons test on main effect of SE and SA. Data are presented as mean ± SEM, * p < 0.05. ** p < 0.01. *** p < 0.001. **** p < 0.0001.

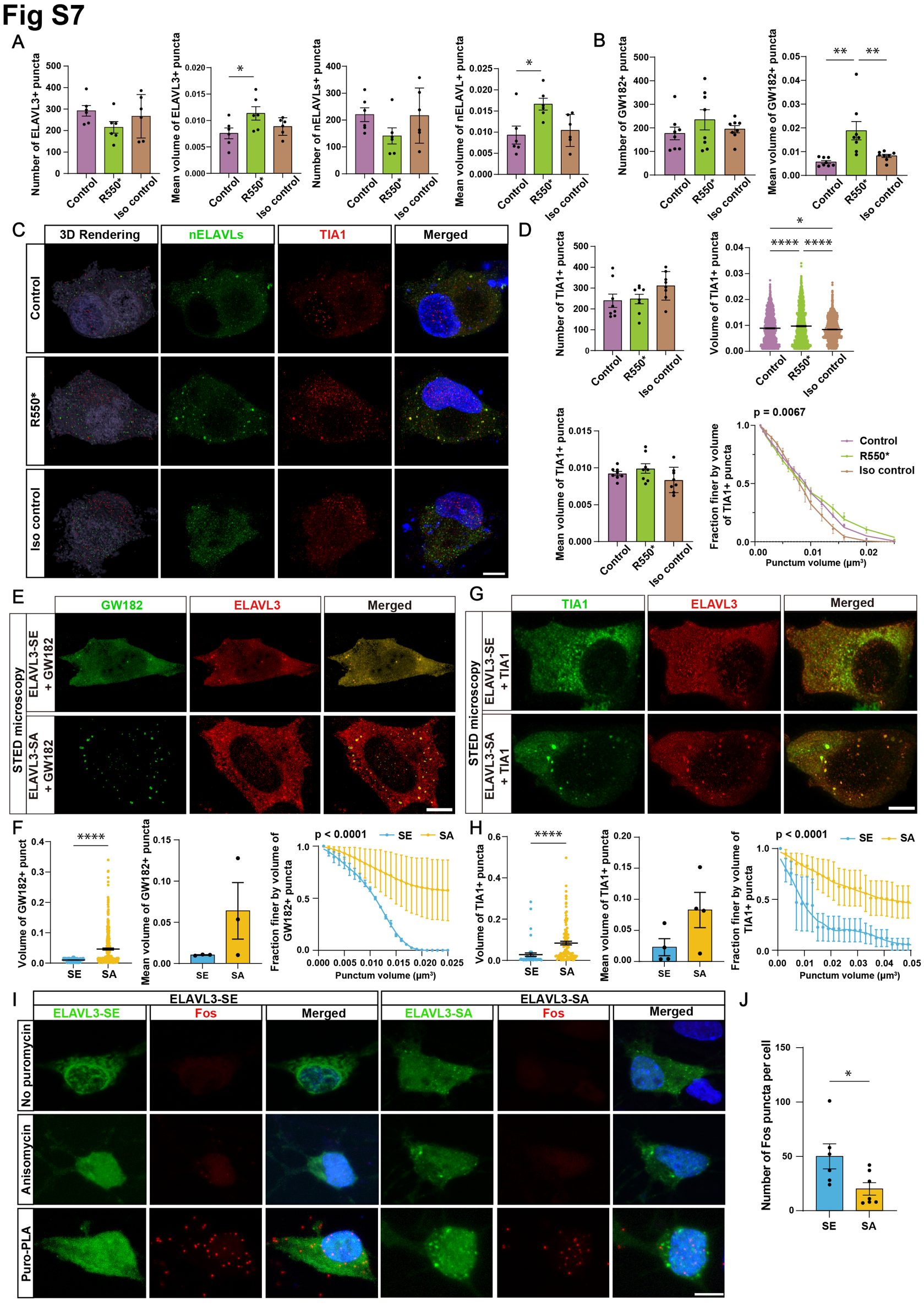

**Figure S7. CDKL5 mutation and nELAVL phosphomutants lead to aberrant nELAVL condensates and altered crosstalk with P-bodies and stress granules.**

(A) Quantification analysis of ELAVL3 (green) and nELAVL (red) puncta in iNeurons of Figure 5F. While the puncta number showed no significant differences, the mean volume of ELAVL3 (green) and nELAVL (red) puncta showed significant increase in R550* iNeurons, compared to control group. n=6 cells from 3 batches of experiments. One-way ANOVA with Tukey’s multiple comparisons test.

(B) Quantification analysis of GW182 (red) puncta in iNeurons of Figure 5H. While the puncta number showed no significant differences, the mean volume of GW182 (red) puncta showed significant increase in R550* iNeurons. n=6 cells from 3 batches of experiments. One-way ANOVA with Tukey’s multiple comparisons test.

(C & D) STED microscopy imaging of nELAVLs (green) and TIA1 (red) in control, R550* and isogenic control iNeurons. The TIA1 punctum volume was larger in R550* iNeurons. Scale bar, 10 μm. n=8 cells from 4 batches of experiments. One-way ANOVA with Tukey’s multiple comparisons test. Volume fraction analysis: Two-way ANOVA with Sidak’s multiple comparisons test on main effect of Control, R550* and Iso control; Multiple comparisons: p = 0.5457 for Control vs R550* p = 0.0095 for R550* vs Iso control.

(E & F) STED microscopy of co-expressed GW182 and SE/SA mutant of ELAVL3 in HT22 cells. Compared to SE, co-expressing with the SA mutant led to a large-shifted GW182 punctum volume distribution. Scale bar, 5μm. n=3 cells from 3 batches of experiments. Mann-Whitney test (two-tailed). Volume fraction analysis: Two-way ANOVA with Sidak’s multiple comparisons test on main effect of SE and SA.

(G & H) STED microscopy of co-expressed TIA1 and SE/SA mutant of ELAVL3 in HT22 cells. Compared to SE, co-expressing with the SA mutant led to a large-shifted TIA1 punctum volume distribution. Scale bar, 5μm. n=4 cells from 3 batches of experiments. Mann-Whitney test (two-tailed). Volume fraction analysis: Two-way ANOVA with Sidak’s multiple comparisons test on main effect of SE and SA.

(I & J) Confocal images showed rat primary neurons expressing ELAVL3-SE/SA stained with puro-PLA for the transcription factor Fos protein (red). The Fos puncta is significantly decreased in SA group. Scale bar, 5 μm. n=6 (SE) and 7 (SA) cells from 3 batches of experiments. Mann-Whitney test (two-tailed). Data are presented as mean ± SEM, * p < 0.05, ** p < 0.001, **** p < 0.0001.

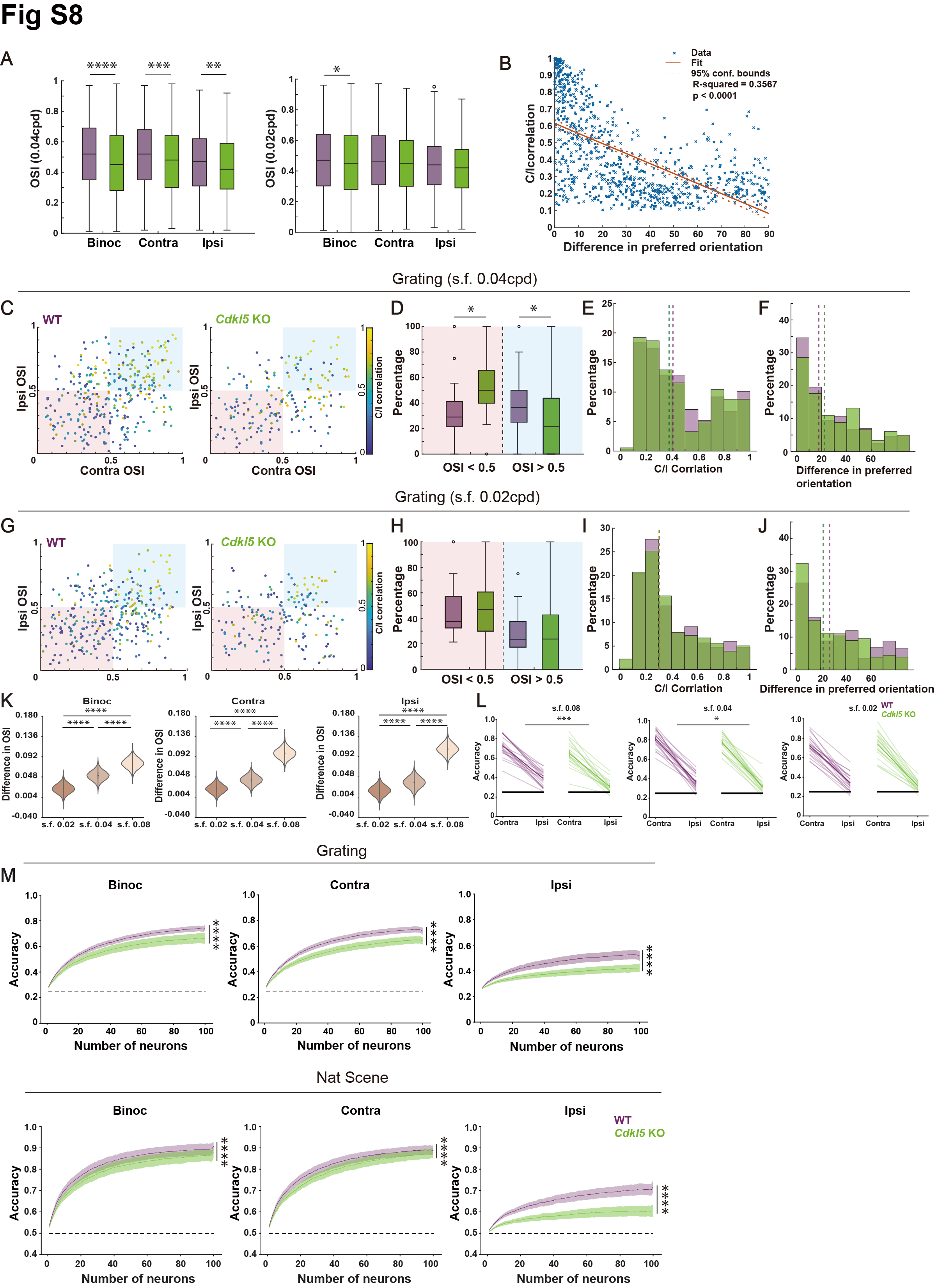

**Figure S8. Visual deficits caused by CDKL5 deletion.**

(A) (Left) Boxplots showing the orientation selective index (OSI) of *Cdkl5* KO mice were reduced across Binoc, Contra and Ipsi conditions under grating (s.f. 0.04cpd). Binoc OSI: 1070 neurons from 6 WT and 877 neurons from 6 *Cdkl5* KO mice. Contra OSI: 1037 neurons from 6 WT and 868 neurons from 6 *Cdkl5* KO. Ipsi OSI: 724 neurons from 6 WT and 435 neurons from 6 *Cdkl5* KO. (Right) Boxplots showing the orientation selective index (OSI) of *Cdkl5* KO mice were reduced in Binoc conditions under grating (s.f. 0.02cpd). Binoc OSI: 995 neurons from 6 WT and 1425 neurons from 6 *Cdkl5* KO mice. Contra OSI: 935 neurons from 6 WT and 1465 neurons from 6 *Cdkl5* KO. Ipsi OSI: 728 neurons from 6 WT and 942 neurons from 6 *Cdkl5* KO. The centerlines represent median values, and the whiskers connect the nonoutlier minimum and maximum values to 0.25 and 0.75 quartiles respectively. Outliers are values greater than 1.5 interquartile range away from the quartiles. Mann-Whitney U test (two-tailed).

(B) The linear regression relationship between the Contra and Ipsi eye correlation and the difference in preferred orientation. Blue “x” symbols represent the data points. The solid orange line indicates the least-squares fit. The dashed orange lines show the 95% confidence intervals of the fit.

(C) Scatter plots comparing OSI and correlation of grating (s.f. 0.04cpd) responsive neurons in WT mice and *Cdkl5* KO mice. Color bar shows the correlation between tuning curves. 327 neurons from 6 WT and 182 neurons from 6 *Cdkl5* KO. Mann-Whitney U test (two-tailed).

(D) Boxplots showing fraction of neurons per FOV within the blue region (OSI > 0.5) and red region (OSI < 0.5) from C. 26 FOV from 6 WT mice and 21 FOV from 6 *Cdkl5* KO mice. Outliers are values greater than 1.5 interquartile range away from the quartiles. Mann-Whitney U test (two-tailed).

(E) Distribution of Contra and Ipsi eye correlation between WT and *Cdkl5* KO under grating (s.f. 0.04cpd) stimuli. 327 neurons from 6 WT and 182 neurons from 6 *Cdkl5* KO. Mann-Whitney U test (two-tailed).

(F) Distribution of difference in preferred orientation between WT and *Cdkl5* KO under grating (s.f. 0.04cpd) stimuli. 327 neurons from 6 WT and 182 neurons from 6 *Cdkl5* KO. Mann-Whitney U test (two-tailed).

(G) Scatter plots comparing OSI and correlation of grating (s.f. 0.02cpd) responsive neurons in WT mice and *Cdkl5* KO mice. Color bar shows the correlation between tuning curves. 318 neurons from 6 WT and 179 neurons from 6 *Cdkl5* KO. Mann-Whitney U test (two-tailed).

(H) Boxplots showing fraction of neurons per FOV within the blue region (OSI > 0.5) and red region (OSI < 0.5) from G. 26 FOV from 6 WT mice and 21 FOV from 6 *Cdkl5* KO mice. Mann-Whitney U test (two-tailed).

(I) Distribution of Contra and ipsi eye correlation between WT and *Cdkl5* KO under grating (s.f. 0.02cpd) stimuli. 318 neurons from 6 WT and 179 neurons from 6 *Cdkl5* KO. Mann-Whitney U test (two-tailed).

(J) Distribution of difference in preferred orientation between WT and *Cdkl5* KO under grating (s.f. 0.02cpd) stimuli. 318 neurons from 6 WT and 179 neurons from 6 *Cdkl5* KO. Mann-Whitney U test (two-tailed).

(K) Difference in OSI values between WT and *Cdkl5* KO under different spatial frequencies. 26 FOVs from 6 WT mice and 21 FOVs from 6 *Cdkl5* KO mice. One-Way ANOVA followed by Tukey's HSD Pairwise Group Comparisons.

(L) At a spatial frequency of 0.08 and 0.04 but not 0.02, *Cdkl5* KO mice had a reduced decoder accuracy compared with WT mice. 26 FOV from 6 WT mice and 21 FOV from 6 *Cdkl5* KO mice. Two-way ANOVA on main effect of WT and *Cdkl5* KO.

(M) Reduced decoder accuracy for grating and natural scene stimuli in *Cdkl5* KO mice. 26 FOV from 6 WT mice and 21 FOV from 6 *Cdkl5* KO mice. Two-way ANOVA on main effect of WT and *Cdkl5* KO. Data are presented as mean ± SEM, * p < 0.05. ** p < 0.01. *** p < 0.001. **** p < 0.0001.

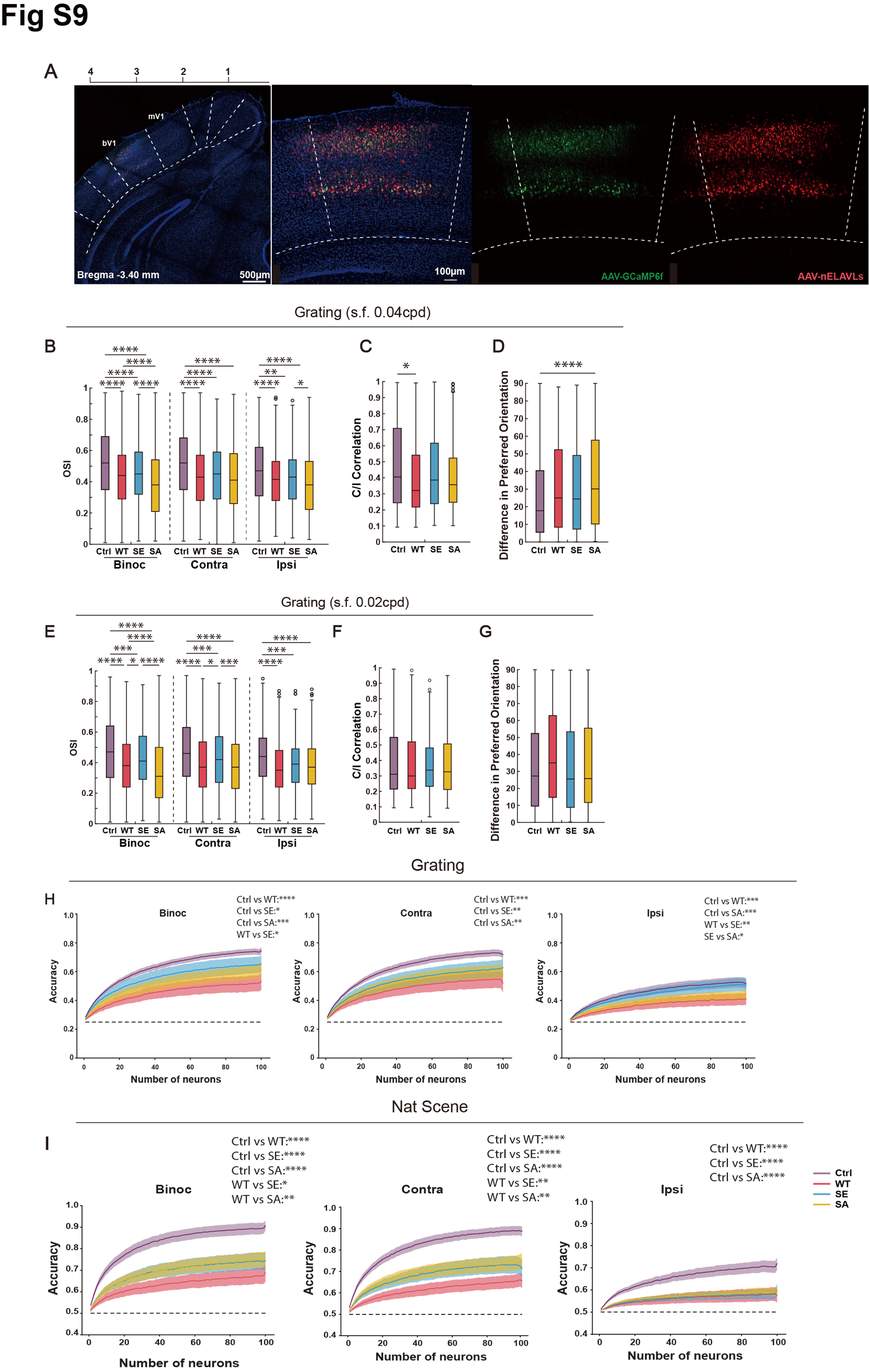

**Figure S9. nELAVL phosphorylation for visual processing under different spatial frequencies.**

(A) *Post-hoc* verification of the virus injection site following two-photon imaging. bV1 is 2.8-3.5 mm lateral from the midline and 3.4 mm posterior to Bregma in the checking image (left). The three images on the right are magnifications of bV1. Green channel indicates AAV-GCaMP6f, red channel indicates AAV-nELAVLs, and blue channel corresponds to DAPI staining.

(B) Boxplots showing OSI were reduced in WT, SE and SA mice across Binoc, Contra and Ipsi conditions under grating (s.f. 0.04cpd). Binoc OSI: 1070 neurons from 6 Ctrl, 822 neurons from 5 WT injected, 589 neurons from 4 SE injected and 1045 neurons from 5 SA injected mice. Contra OSI: 1037 neurons from 6 Ctrl, 791 neurons from 5 WT injected, 583 neurons from 4 SE injected and 1033 neurons from 5 SA injected mice. Ipsi OSI: 724 neurons from 6 Ctrl, 594 neurons from 5 WT injected, 414 neurons from 4 SE injected and 655 neurons from 5 SA injected mice. Kruskal-Wallis test followed by Tukey’s multiple comparisons test.

(C) Boxplots showing Contra and Ipsi correlation under grating (s.f. 0.04cpd) stimuli in Ctrl, WT injected, SE injected and SA injected mice. 327 neurons from 6 Ctrl, 234 neurons from 5 WT injected, 153 neurons from 4 SE injected and 293 neurons from 5 SA injected mice. Kruskal-Wallis test followed by Tukey’s multiple comparisons test.

(D) Boxplots showing difference in preferred orientation under grating (s.f. 0.04cpd) stimuli in Ctrl, WT injected, SE injected and SA injected mice. 327 neurons from 6 Ctrl, 234 neurons from 5 WT injected, 153 neurons from 4 SE injected and 293 neurons from 5 SA injected mice. Kruskal-Wallis test followed by Tukey’s multiple comparisons test.

(E) Boxplots showing OSI were reduced in WT, SE and SA mice across Binoc, Contra and Ipsi conditions under grating (s.f. 0.02cpd). Binoc OSI: 995 neurons from 6 Ctrl, 899 neurons from 5 WT injected, 573 neurons from 4 SE injected and 1107 neurons from 5 SA injected mice. Contra OSI: 935 neurons from 6 Ctrl, 896 neurons from 5 WT injected, 620 neurons from 4 SE injected and 1062 neurons from 5 SA injected mice. Ipsi OSI: 728 neurons from 6 Ctrl, 590 neurons from 5 WT injected, 349 neurons from 4 SE injected and 592 neurons from 5 SA injected mice. Kruskal-Wallis test followed by Tukey’s multiple comparisons test.

(F) Boxplots showing Contra and Ipsi correlation under grating (s.f. 0.02cpd) stimuli in Ctrl, WT injected, SE injected and SA injected mice. 318 neurons from 6 Ctrl, 267 neurons from 5 WT injected, 131 neurons from 4 SE injected and 280 neurons from 5 SA injected mice. Kruskal-Wallis test followed by Tukey’s multiple comparisons test.

(G) Boxplots showing difference in preferred orientation under grating (s.f. 0.02cpd) stimuli in Ctrl, WT injected, SE injected and SA injected mice. 318 neurons from 6 Ctrl, 267 neurons from 5 WT injected, 131 neurons from 4 SE injected and 280 neurons from 5 SA injected mice. Kruskal-Wallis test followed by Tukey’s multiple comparisons test.

(H) Decoder accuracy for Ctrl, WT injected, SE injected and SA injected mice under grating stimuli based on each FOVs under grating (s.f. 0.08cpd) stimuli. 22 FOVs from 6 Ctrl, 15 FOV from 5 WT injected, 16 FOVs from 4 SE injected and 21 FOVs from 5 SA injected mice. Mixed linear model followed by Tukey's HSD Pairwise Group Comparisons on main effect between groups.

(I) Decoder accuracy for Ctrl, WT injected, SE injected and SA injected mice under natural scene stimuli based on each FOVs under natural scene stimuli. 22 FOVs from 6 Ctrl, 15 FOV from 5 WT injected, 16 FOVs from 4 SE injected and 21 FOVs from 5 SA injected mice. Mixed linear model followed by Tukey's HSD Pairwise Group Comparisons on main effect between groups. Data are presented as mean ± SEM, * p < 0.05. ** p <0.01. *** p < 0.001. **** p < 0.0001.

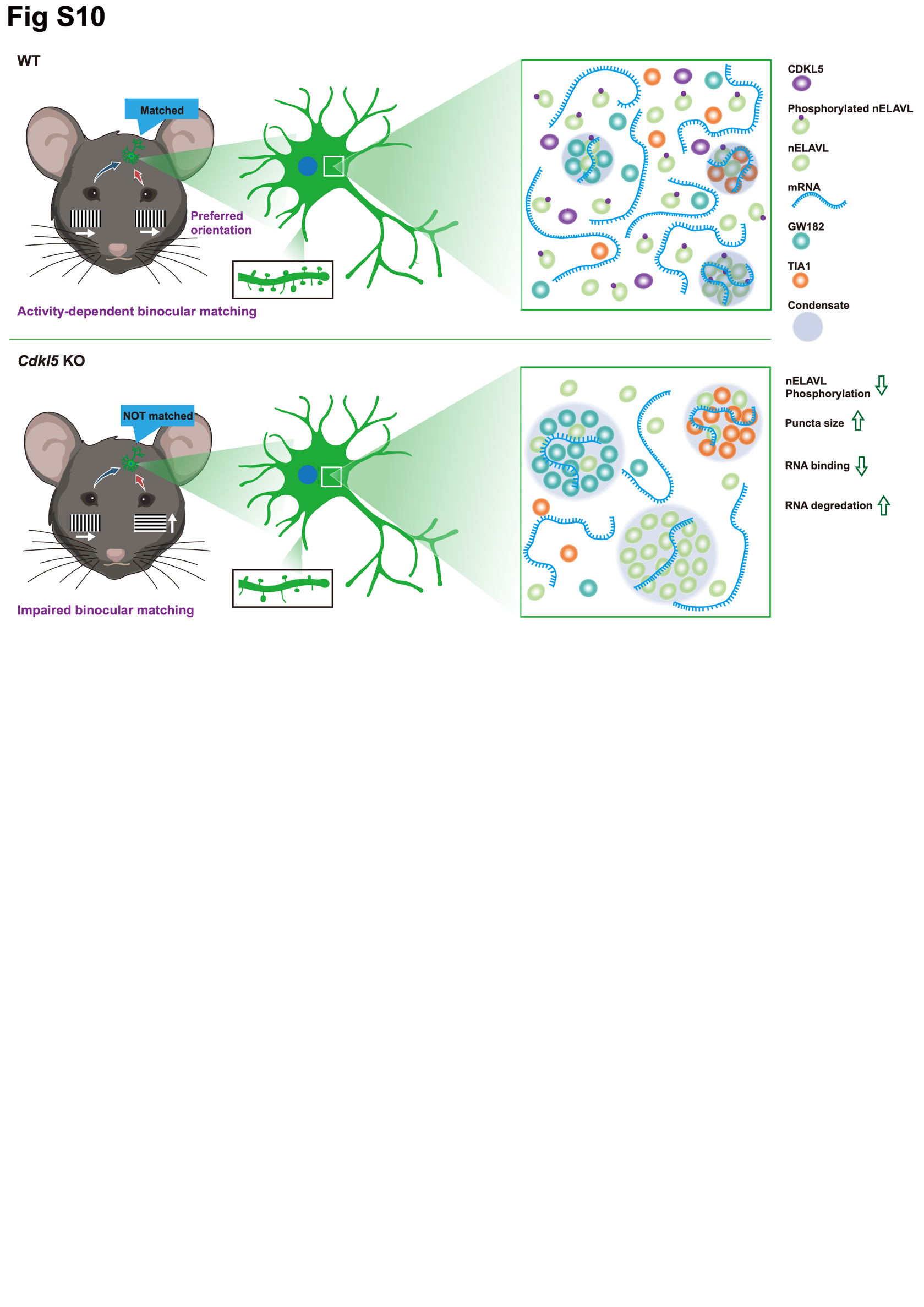

**Figure S10. Working model of CDKL5 regulating biomolecular condensates to guide binocular visual circuit refinement.**

In WT mice, CDKL5 phosphorylates nELAVLs in an activity-dependent manner and regulates the dynamics of nELAVL condensates to modulate the mRNA levels of activity-dependent genes, thereby refining binocular visual circuits in bV1. In *Cdkl5* KO mice, the lack of phosphorylation on nELAVLs leads to aberrant condensate formation and dysregulated crosstalk with other biomolecular condensates such as GW182+ P-bodies and TIA1+ stress granules. Consequently, *Cdkl5* KO bV1 neurons exhibit altered dendritic spine morphology and impaired visual cortical processing.

Supplementary Tables S1-S5 title:

Table S1. Phosphopeptides and summary statistics.

Table S2. DEG analysis from bulk RNA sequencing.

Table S3. DEG analysis from snRNA sequencing.

Table S4. DEG analysis in glutamatergic neurons.

Table S5. Percentage of responsive neurons in 2P imaging.

Table S6. Neurons response amplitude in 2P imaging
